# Oligonucleotide-directed proximity-interactome mapping (O-MAP): A unified method for discovering RNA-interacting proteins, transcripts and genomic loci *in situ*

**DOI:** 10.1101/2023.01.19.524825

**Authors:** Ashley F. Tsue, Evan E. Kania, Diana Q. Lei, Rose Fields, Christopher D. McGann, Elliot Hershberg, Xinxian Deng, Maryanne Kihiu, Shao-En Ong, Christine M. Disteche, Sita Kugel, Brian J. Beliveau, Devin K. Schweppe, David M. Shechner

## Abstract

Throughout biology, RNA molecules form complex networks of molecular interactions that are central to their function, but remain challenging to investigate. Here, we introduce Oligonucleotide-mediated proximity-interactome MAPping (O-MAP), a straightforward method for elucidating the biomolecules near an RNA of interest, within its native cellular context. O-MAP uses programmable oligonucleotide probes to deliver proximity-biotinylating enzymes to a target RNA, enabling nearby molecules to be enriched by streptavidin pulldown. O-MAP induces exceptionally precise RNA-localized *in situ* biotinylation, and unlike alternative methods it enables straightforward optimization of its targeting accuracy. Using the 47S pre-ribosomal RNA and long noncoding RNA *Xist* as models, we develop O-MAP workflows for unbiased discovery of RNA-proximal proteins, transcripts, and genomic loci. This revealed unexpected co-compartmentalization of *Xist* and other chromatin-regulatory RNAs and enabled systematic characterization of nucleolar-chromatin interactions across multiple cell lines. O-MAP is portable to cultured cells, organoids, and tissues, and to RNAs of various lengths, abundances, and sequence composition. And, O-MAP requires no genetic manipulation and uses exclusively off-the-shelf parts. We therefore anticipate its application to a broad array of RNA phenomena.

## MAIN

In the context of the cell, very little RNA is naked^1-3^. Direct binding interactions with other biomolecules (proteins, RNAs, genomic loci) regulate all aspects of an RNA’s lifecycle, including biogenesis, localization, turnover, and protein-coding or noncoding functions^4, 5^. Moreover, higher-order interactions between transcripts and their local microenvironment are critical for organizing subcellular architecture and compartmentalization^6, 7^. In humans, for example, RNAs are central determinants of chromatin folding^8-10^, and they nucleate and scaffold a host of biomolecular condensates that collectively control cellular metabolic, epigenetic, and stress-signaling pathways^11-13^. Yet, most of these critical structures have eluded detailed molecular characterization, due to a lack of methods for elucidating RNA interactions at compartment-level (nm–µm) distances^6, 14^.

Most state-of-the-art RNA interaction-discovery approaches use biotinylated antisense oligonucleotides to pull down a target RNA and its molecular partners from cell lysates^14-17^. While powerful, these approaches are limited in several key regards. Because it is challenging to optimize the specificity of the oligo probe pool, these approaches are limited by off-target RNA capture^18^. Moreover, enriching RNA from crude lysates can be plagued by artifactual interactions with abundant, nonspecific RNA-binding proteins, leading to false positives^19^. Overcoming this experimental background often requires large input masses (∼10^8^ cells)^17^, especially for low-abundance RNAs. Finally, because these techniques probe an RNA *ex vivo*, they cannot capture higher-order interactions that depend upon intact subcellular structure.

We hypothesized that many of these obstacles could be addressed using *in situ* proximity-biotinylation^20^. In this approach, promiscuous biotinylating enzymes (*e*.*g*. peroxidases like Horseradish Peroxidase, HRP, and APEX)^21-23^ are localized to a subcellular site of interest, typically by transgenic expression^20^. Upon addition of substrate, these enzymes generate reactive biotin species that diffuse outward and covalently tag nearby molecules *in situ*, enabling these molecules to be enriched by simple streptavidin pulldown and identified by mass-spectrometry or high-throughput sequencing^24-26^. Yet, while proximity-biotinylation has emerged as a powerful tool for protein-targeted interaction discovery, applying the technique to RNAs remains challenging. Unlike proteins, transcripts cannot be genetically fused to the biotinylating enzyme, and hence most strategies seek to engineer artificial complexes between the enzyme and its target RNA^21, 27, 28^. But, these transgenic approaches often produce substantial pools of mislocalized or unbound biotinylating enzymes, resulting in nonspecific background labeling that can blur the experimental signal^29^. These methods also require complex cell engineering to simultaneously overexpress multiple components, often including the RNA itself^27^, making them challenging to apply to low-abundance RNAs, and altogether impossible in difficult-to-engineer lines, organisms, and clinical samples. While a handful of methods (*e*.*g*. RNA-TRAP and HyPro)^30, 31^, potentially overcome this barrier by targeting individual transcripts without transgenic manipulation, these approaches rely on custom reagents that are cumbersome to synthesize, vet, and optimize^30, 31^.

To overcome these challenges, we here present Oligonucleotide-directed proximity-interactome mapping (O-MAP), a straightforward and flexible method for applying proximity-labeling to individual RNA targets in genetically unmodified samples. O-MAP utilizes the same peroxidase/tyramide chemistry used in APEX-based proximity-omics approaches^23, 24^, but it relies on programmable oligonucleotide probes, rather than transgenic expression, to deploy biotinylating enzymes to endogenous target RNAs (**Fig. 1a**). This approach enables precise RNA-targeting that can be easily optimized and experimentally validated, overcoming a chief limitation of oligo-pulldown based approaches. Building on this, we demonstrate O-MAP mass-spectrometry (O-MAP-MS), RNA-Sequencing (O-MAP-Seq), and DNA-Sequencing (O-MAP-ChIP) workflows for unbiased, compartment-wide RNA-interaction discovery. These require markedly fewer (20–100-fold) cells than established methods, and use only inexpensive, off-the-shelf parts and standard manipulations. Using these workflows, we discover new interactions within the nucleolus and Barr bodies—RNA-scaffolded compartments that are difficult to isolate biochemically. Finally, we demonstrate that O-MAP can be readily applied to cell lines, organoids, and tissues, as well as to diverse classes of target RNAs, suggesting its utility as a broad-use RNA interaction-discovery tool.

**Figure 1.**
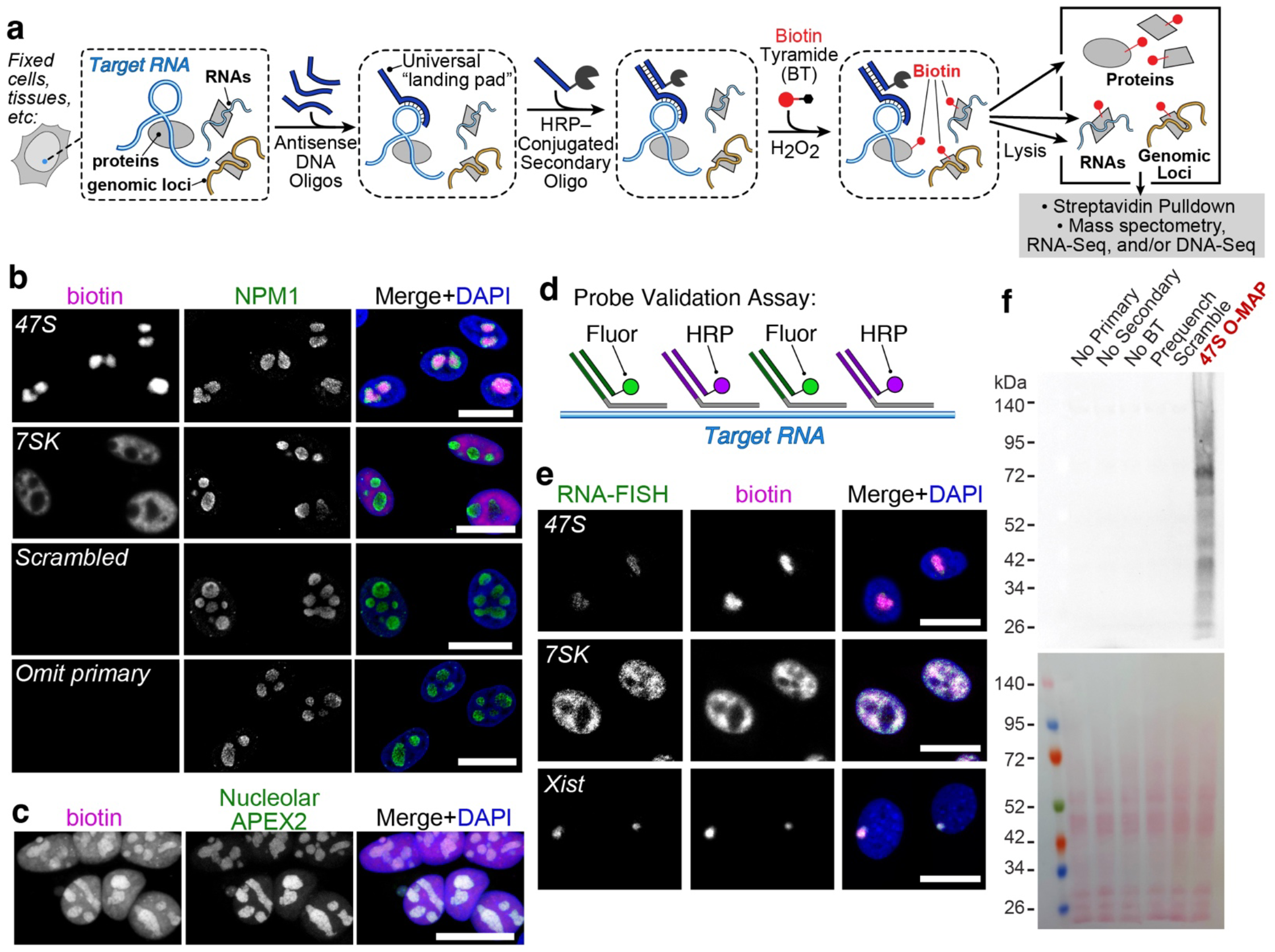
O-MAP Design and Implementation. **a**, Overview of O-MAP. Specimens are chemically fixed, and pools of antisense DNA probes are hybridized to the target RNA. These probes recruit a common, HRP-conjugated secondary probe that catalyzes *in situ* proximity-biotinylation. **b**, O-MAP enables precise RNA-targeted proximity labeling, enabling interaction-discovery. *In situ* biotinylation imaged by neutravidin staining; *NPM1* by immunofluorescence. Note nucleolar biotinylation in 47S O-MAP, nucleoplasmic biotinylation in 7SK O-MAP. HeLa cells. **c**, Nucleolar-targeted APEX2 exhibits substantial off-target, nucleoplasmic labeling^25^. **d**, O-MAP Probe Validation Assay. Primary probes are split into sub-pools that enable O-MAP and RNA-FISH to be performed simultaneously. Lack of co-localization suggests probe off-targeting. **e**, Probe Validation Assays on 47S-pre-rRNA and 7SK, in HeLa cells, and *Xist*, in mouse Patski cells. **f**, Recovery of O-MAP-biotinylated proteins. *Top:* Streptavidin-HRP blot of whole cell lysates. Note ladder of biotinylated proteins from 47S O-MAP *Bottom:* Ponceau stain. All scale bars: 20 µm.

## Results

### O-MAP Design and Implementation

Recent advancements in RNA-Fluorescent *In Situ* Hybridization (RNA-FISH) oligonucleotide probe design have dramatically improved the method’s sensitivity, flexibility, and specificity, enabling nearly any transcript to be imaged with high precision^32-35^. In developing O-MAP, we aimed to leverage these advancements to deploy the proximity-labeling enzyme HRP, rather than fluorophores, to a target RNA. As our primary model system, we chose the human 47S pre-ribosomal RNA, the long noncoding RNA that scaffolds the nucleolus^11, 36^. To specifically target the 47S precursor (and avoid mature ribosomes) we selected an established RNA-FISH probe set^37^ against ITS1, a “transcribed spacer” domain that is degraded during ribosome biogenesis and which never departs from the nucleolus (**Supplementary Fig. 1** and **Supplementary Table 1**)^38^. We used these probes to explore different HRP recruitment strategies, by appending them with functional modules that can be targeted by HRP conjugates and scoring the resulting nucleolar biotinylation by neutravidin staining (**Supplementary Fig. 2**). Several of these strategies used small-molecule haptens (*e*.*g*. Digoxigenin) to recruit an HRP-conjugated hapten-binding protein—similar to classical Tyramide Signal Amplification (TSA)-FISH and related interaction-discovery methods^26, 30, 31, 39, 40^. Yet, while these strategies induced prominent nucleolar biotinylation, they also exhibited variable off-target labeling that frequently rivaled the nucleolar signal (**Supplementary Fig. 2**).

In contrast, our optimized O-MAP strategy relies exclusively on programable oligonucleotide hybridization to precisely target HRP to an RNA of interest (**Fig. 1a**). In O-MAP, cells or tissues are chemically fixed with formaldehyde, and pools of oligo probes are annealed to the target RNA. These probes are chemically unmodified, but are appended with universal “landing pad” sequences originally developed for the FISH techniques Oligopaint^41^ and SABER^35^. In a subsequent hybridization step, these landing pads recruit a common secondary oligo that is conjugated to HRP^42^. Upon addition of biotin-tyramide and hydrogen peroxide, HRP generates short-lived, highly reactive phenoxyl radicals that diffuse outward and pervasively biotinylate molecules near (∼10 nm) the target RNA^24^, enabling their enrichment.

Unlike hapten-based HRP-recruitment strategies (**Supplementary Fig. 2**), 47S-targeted O-MAP induced prominent and precise nucleolar biotinylation with nearly undetectable background (**Fig. 1b**). Likewise, O-MAP targeting *7SK*—a small noncoding RNA thought to reside in nuclear speckles^43^—produced exclusively nucleoplasmic labeling. For each RNA, similar results were observed using multiple distinct probe sets, including probes designed using the OligoMiner Pipeline^34^ to have different hybridization parameters (**Supplementary Fig. 3 and Supplementary Table 1**). Importantly, omitting any component of the O-MAP workflow (primary oligos, the oligo-HRP conjugate, HRP substrates), or using scrambled primary probes, completely ablated biotinylation (**Figs. 1b** and **Supplementary Fig. 4; Supplementary Table 1**). Moreover, 47S-targeted O-MAP appeared markedly more spatially refined than the analogous genetically encoded approach—nucleolar-targeted APEX2^25, 44^—which generated substantial off-target, nucleoplasmic biotinylation (**Fig. 1c**).

O-MAP’s landing pad design also enables a straightforward strategy for optimizing the specificity of its probe pool, thus overcoming a longstanding challenge of oligo pulldown-based approaches^14-17^ **(Fig. 1d**). Derived from a standard RNA-FISH method^33^, our O-MAP Probe Validation Assay first groups probes into “odd” and “even” sub-pools. Each sub-pool is outfitted with a distinct landing pad that recruits either an HRP-conjugated or fluorescent secondary oligo, enabling O-MAP and RNA-FISH to be performed simultaneously. Off-targeting (*i*.*e*. non-colocalizing) probes are then readily identified and eliminated. We applied this assay to our 47S- and 7SK-targeting probe sets, and in both cases observed strong overlap between the O-MAP and FISH signals, further underscoring O-MAP’s precision (**Fig. 1e**). We then tested this assay on a less abundant

RNA with a more confined localization. For this we chose *Xist*, the long noncoding RNA (lncRNA) that drives X-chromosome inactivation, and which coats the inactive X-chromosome (Xi)^45^. As predicted, *Xist* O-MAP induced strong biotinylation in a single prominent nuclear punctum, and which exclusively co-localized with the *Xist* RNA-FISH signal (**Fig. 1e**). Importantly, O-MAP’s use of formaldehyde crosslinking and formamide denaturation did not appear to limit the recovery and enrichment of biotinylated proteins, as demonstrated with both 47S and 7SK (**Fig. 1f** and **Supplementary Fig. 5**). Taken with results above, this suggests that O-MAP generates exceptionally precise, RNA-targeted *in situ* biotinylation compatible with recovery and downstream analysis. We therefore sought to develop a suite of tools for discovering proteins (**O-MAP-MS**), transcripts (**O-MAP-Seq**), and genomic loci (**O-MAP-ChIP**) interacting with a target RNA.

### Mapping RNA-proximal proteomes with O-MAP-MS

To develop our O-MAP-MS pipeline our primary test case was the nucleolus—the subnuclear organelle that compartmentalizes and controls ribosome biogenesis—in HeLa cells^11, 36^. Our approach emulated the ratiometric quantification strategy used in APEX-based proximity-proteomics, which enhances precision by measuring the relative abundances of peptides within a target compartment and the adjoining subcellular space^46^ (**Fig. 2a**). In parallel experiments, we used 47S O-MAP to label nucleoli, 7SK-targeting probe sets (**Supplementary Fig. 3**) as proxies for the neighboring nucleoplasmic compartment, and scrambled probes to model nonspecific background biotinylation. After O-MAP labeling, biotinylated proteins were enriched under denaturing conditions, and protein abundances were measured by Tandem Mass Tag (TMTPro) quantitative mass spectrometry^47^. Owing to the reduced background labeling and tyramide amplification, our initial proof-of-principle O-MAP-MS experiments required 20–100-fold less starting material (5.5 million cells per replicate) than nucleolar biochemical fractionation^48^ or the oligo pulldown method ChIRP–MS^17^.

**Figure 2.**
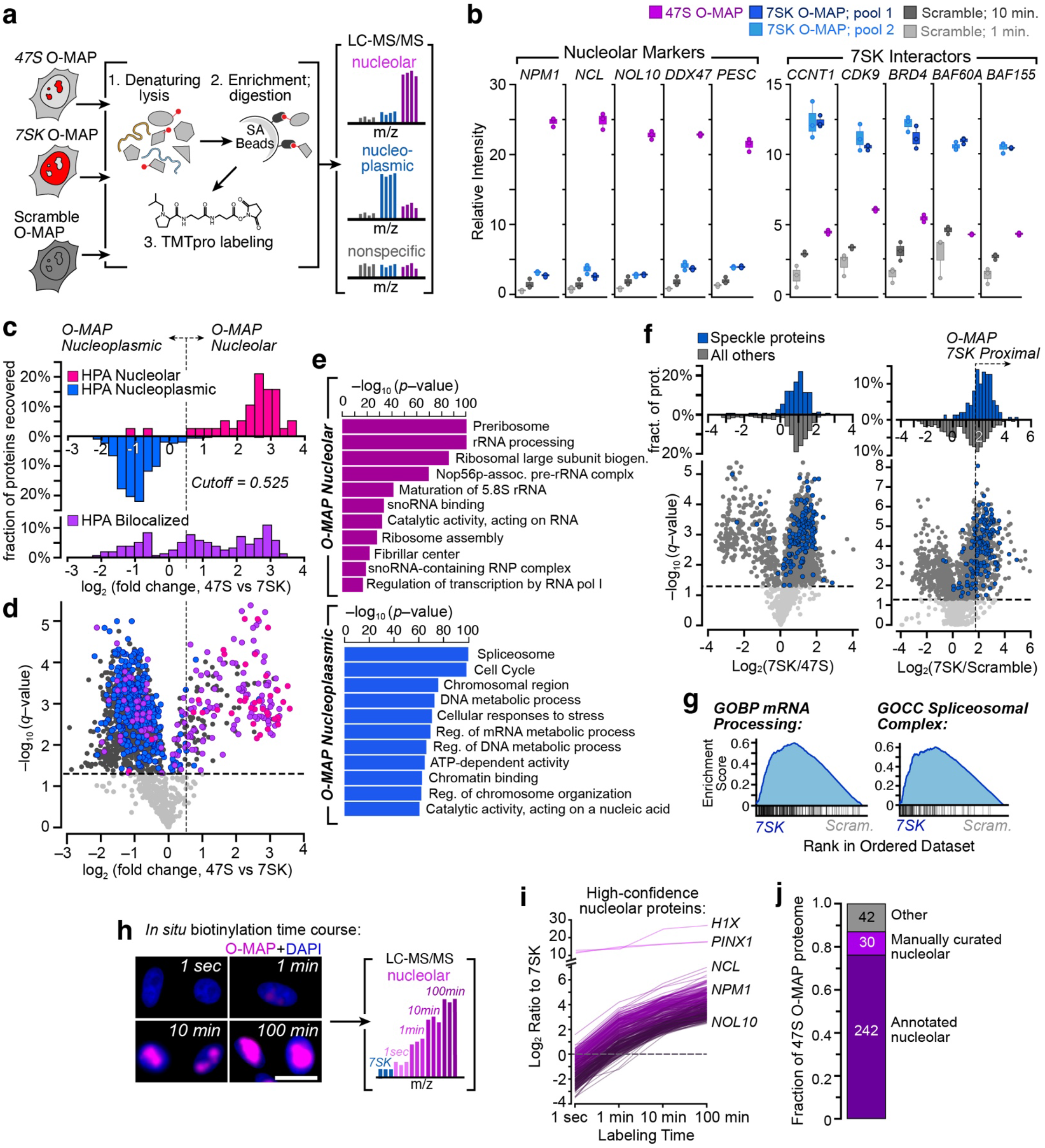
O-MAP-MS for probing RNA-proximal proteomes. **a**, Strategy for characterizing the HeLa nucleolar proteome. Parallel O-MAP experiments targeting the 47S (nucleolar), 7SK (nucleoplasmic) or using scrambled probes (background) were processed as indicated. Samples were quantified by TMTPro mass-spectrometry; each replicate (*n* = 3 biological replicates) was labeled with a unique mass-tag. **b–g**, analysis of a single-shot 47S/7SK O-MAP-MS experiment, using 5.5×10^6^ cells per replicate. **b**, Recovery of known nucleolar proteins (*left*) and 7SK interactors (*right*). Two different probe sets, requiring different labeling times, were used in 7SK experiments; these were time-matched to scrambled-probe controls. Box whisker plots with individual data points shown. **c–d**, Global enrichment of known nucleolar (*magenta*), nucleoplasmic (*blue*), and bi-localized (*purple*) marker proteins, defined by the Human Protein Atlas (HPA). Dotted line denotes the optimal threshold separating the compartments, determined by Receiver-Operating Characteristic (ROC) analysis (**Supplementary Fig. 8**). **c**, Histograms plotting the enrichment of each sub-compartmental proteome. Only proteins with *q-*value ≤ 0.05 are shown. Note conspicuous separation between nucleolar and nucleoplasmic markers, and multi-modal distribution of bi-localized proteins. **d**, Volcano plot showing all data. **e**, Gene Ontology (GO) analysis for the O-MAP-nucleolar (*top*) and O-MAP-Nucleoplasmic (*bottom*) proteomes, defined from ROC analysis. The top eleven most enriched terms are shown. **f**, 7SK O-MAP enriches the Nuclear Speckle proteome. Histograms and volcano plots as in **c**,**d**, showing enrichment of speckle proteins in both 7SK- vs-47S and 7SK-vs-Scramble comparisons. The latter comparison more clearly distinguished speckle proteins from the broader nucleoplasm and was used in ROC analysis to determine the optimal threshold cutoff (**Supplementary Fig 9**). **g**, Gene Set Enrichment Analysis (GSEA) of the “O-MAP 7SK-proximal” proteome reveals a striking enrichment in RNA splicing-relevant Gene Ontology Biological Process (GOBP) and Cellular Component (GOCC) terms. **h–j**, Higher coverage of the nucleolar proteome using an O-MAP *in situ* biotinylation time course. **h**, Approach. 47S O-MAP was performed in parallel for the indicated times, enriched, and TMTPro-labeled as above (*n* = 3). A single 7SK probe set and time point was used for normalization. *In situ* biotinylation visualized by neutravidin staining. Scale bars: 20 µm. **i**, k-medoid clustering yields a clade of 313 high-confidence nucleolar proteins. Notable marker genes are indicated. **j**, nearly all (87%) members of the nucleolar medoid group corresponds to annotated or manually curated nucleolar proteins.

Yet, even with this small input, O-MAP-MS was able to accurately capture targeted subcellular compartments at considerable depth (1954 total proteins detected) and reproducibility (Pearson’s correlations ranging from 0.77–0.99 between replicates; **Supplementary Fig. 6**). Stereotypic nucleolar markers (*e*.*g*., NPM1, NCL), were markedly enriched by 47S O-MAP (**Fig. 2b**, *left*), while 7SK O-MAP highly enriched both classical components of the 7SK RNP (*e*.*g*. CDK9; Cyclin T1)^49^, and recently discovered interactors like BAF155 and BAF160A^50^ (**Fig. 2b**, *right*, **and Supplementary Fig. 7**). To expand on this, we examined more extensive sets of nucleolar and nucleoplasmic marker proteins, using the Human Protein Atlas^51^ (HPA) to define markers exclusively localized within each compartment (38 nucleolar and 305 nucleoplasmic proteins observed in our data), and a cohort of proteins bilocalized between the two (155 in our data, **Supplementary Table 2**). As demonstrated in (**Fig. 2c**,**d**), we observed striking enrichment of resident nucleolar proteins exclusively from 47S O-MAP-MS (95% of markers, average of 5.1-fold enrichment), and converse enrichment of resident nucleoplasmic proteins exclusively from 7SK O-MAP-MS (98% of markers, average of 2.7–fold enrichment; *p =* 6.83×10^−41^, Fisher’s Exact Test). Bilocalized proteins exhibited a multi-modal distribution, with 64% being more strongly enriched by 47S-than 7SK-O-MAP (average 2.1-fold enrichment, **Fig. 2c**).

To further assess O-MAP’s spatial precision, we performed hypothesis-unbiased analysis of the putative nucleolar (47S-proximal) and nucleoplasmic (7SK-proximal) proteomes identified by our data. We used Receiver-Operating Characteristic (ROC) analysis to determine the optimal threshold separating these two groups (log_2_(47S/7SK) = 0.523, **Supplementary Figs. 8–9**), thereby defining candidate lists of 258 O-MAP-Nucleolar and 1396 O-MAP-Nucleoplasmic proteins (**Supplementary Table 2**). Gene Ontology (GO)^52^ and Gene Set Enrichment Analysis (GSEA)^53^ of candidate nucleolar proteins revealed substantial enrichment of factors driving the transcription and processing of pre-ribosomal RNA, ribosome assembly, and nucleolar architecture, as expected (**Fig 2e**, *top* and **Supplementary Fig. 10**). Likewise, the putative nucleoplasmic proteome was highly enriched in factors involved in chromatin organization, mRNA biogenesis and DNA metabolism (**Fig. 2e**, *bottom*). We furthermore examined if O-MAP might reveal more targeted analysis into the 7SK-proximal subnuclear compartment. 7SK has been suggested to accumulate in Nuclear Speckles— membraneless nuclear bodies with putative roles in mRNA biogenesis and processing^43, 54^. Remarkably, of the 117 established speckle factors observed in our data (**Supplementary Table 2**), nearly all were strongly enriched by 7SK O-MAP-MS, relative to either 47S O-MAP or scramble controls (97% and 79% enriched, respectively, **Fig. 2f**). We used ROC analysis to define an optimal threshold cutoff for the speckle factors, noting that these factors were significantly more enriched from the broader nucleoplasmic proteome when comparing 7SK O-MAP to scrambled controls, rather than to 47S O-MAP (**Supplementary Fig. 9**). This analysis revealed a list of 510 candidate 7SK-proximal proteins, which were highly enriched in factors involved in mRNA biogenesis, especially pre-mRNA splicing (**Fig. 2g** and **Supplementary Figs. 9–10**). This further reinforces 7SK’s proposed localization to nuclear speckles, and these bodies’ putative role as splicing hubs^43, 54^.

We next explored the temporal dynamics of O-MAP labeling. We reasoned that, during a labeling time course, proteins within a target compartment would be biotinylated with distinct kinetics from those adjoining it, allowing us to identify richer sets of compartment-specific factors by classifying common temporal profiles. To test this, we performed 47S-targeted O-MAP at times ranging from 1 second to 100 minutes, using a single 7SK time point as a normalization control (**Fig. 2h**). From K-medoid (k=12) clustering, four clusters were significantly enriched for HPA-defined nucleolar proteins (Fisher’s exact test < 0.05; **Supplementary Fig. 11**), generating an expanded list of 313 candidate nucleolar proteins (**Fig. 2i** and **Supplementary Table 3**). Recovery of these proteins appeared to spike after one minute of *in situ* biotinylation and plateau by 10 minutes (**Supplementary Fig. 12**). Encouragingly, 241 (77%) proteins in this cluster have established roles in nucleolar organization or ribosome biogenesis, as annotated by the GO, HPA and Uniprot databases^55^ (**Fig. 2j**). Manual curation of the remaining factors revealed 31 (10%) that likely also play roles in these processes, but were misannotated (**Fig. 2j** and **Supplementary Table 3**). This demonstrates the potential power of this temporal profiling approach, which would be intractable by live-cell proximity-labeling due to the confounding effects of diffusion^56^. Taken with the above, these data strongly support O-MAP’s ability to discover RNA-proximal proteomes with high precision.

### Mapping RNA-proximal transcriptomes with O-MAP-Seq

Having established O-MAP as an RNA-targeted proteomic tool, we next sought to expand the technique to transcriptomic analysis—mapping the transcripts localized near a target RNA. However, because tyramide-radical chemistry is markedly less efficient at labeling nucleic acids than proteins^26^, we anticipated that the direct capture of *in situ-*biotinylated RNA would be challenging, and require large-scale input cell growths^26^. Therefore, we adopted a strategy based on APEX-RIP, in which formaldehyde crosslinks are retained during cell lysis and enrichment, and RNAs within the target compartment are captured by pulling down the biotinylated proteins to which they’re covalently bound^23^ (**Fig. 3a**). The high capture efficiency of this approach enabled us to precisely map RNA-proximal transcriptomes from as few as 8.3×10^6^ cells, approximately 12-24–fold lower than that required by oligo-capture-based methods^14, 57^.

**Figure 3.**
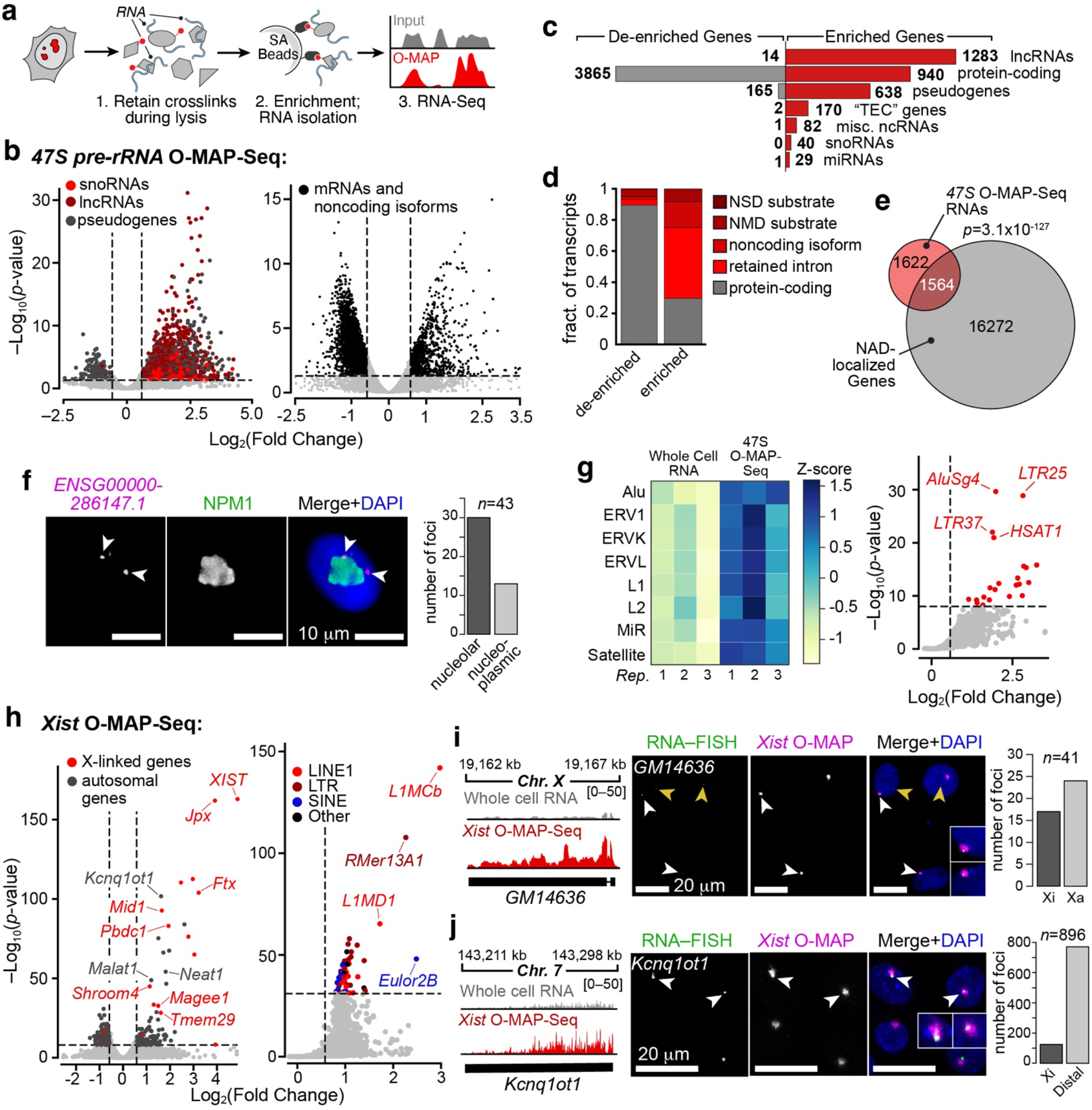
O-MAP-Seq for probing RNA-proximal transcripts. **a**, Approach. *In situ* biotinylated proteins are used to enrich nearby transcripts, which are then quantified by RNA-sequencing. **b–f**, O-MAP-Seq characterization of the HeLa nucleolar transcriptome, **b**, Volcano plots of 47S O-MAP-Seq data, demonstrating enrichment of snoRNAs, lncRNAs, pseudogenes (*left*), and de-enrichment of mRNAs (*right*) (*n* = 3 biological replicates). **c**, Summary of enriched and de-enriched RNA classes in the nucleolar transcriptome. **d**, Nucleolar transcripts expressed from protein-coding genes are predominantly noncoding variants; de-enriched transcripts are predominantly mRNAs. NSD: nonstop decay; NMD: nonsense-mediated decay. **e**, Nearly half of the nucleolar transcriptome is encoded from loci within nucleolar-associated chromatin loci (NADs; **Fig. 4**). Fisher’s exact test. **f**, O-MAP-Seq identifies novel nucleolar-localized transcripts. Note co-localization between *ENSG00000-286147*.*1* RNA-FISH and *NPM1* Immunofluorescence (*arrows;* highlighted in zoom insets), quantified on right. **g**, Nucleolar transcripts are enriched in Transposable Element (TE) domains. *Left:* Z-scores of variance stabilizing transformed (VST) data, corresponding to major TE families. *Right*: volcano plot of individual TE classes in *47S* O-MAP-Seq data. **h–j**, Characterizing transcripts near the inactive X-chromosome (Xi), in Patski cells. **h**, Volcano plots of *Xist* O-MAP-Seq data. *Left:* enrichment for X-linked transcripts that escape X-chromosome inactivation (XCI, *red*) and autosomal transcripts, including several chromatin-regulatory lncRNAs (indicated). *Right:* enrichment for several classes of X-linked transposable elements. **i**, *Gm14636* is a novel XCI-escape gene.*Left: Gm14636* enrichment by O-MAP-Seq. *Middle:* co-localization of *Gm14636* RNA-FISH and *Xist* O-MAP Note penetrant mono- or bi-allelic expression from the Xa (*gold arrows*) and/or Xi (*white arrowheads*), quantified on the right. **j**, *Kcnq1ot1* localizes near the Xi. Panel arrangement parallels that of **i**.

As a first test case, we applied O-MAP-Seq to the HeLa nucleolus, using the *47S*-targeting probe set established above (**Fig. 1**). The nucleolar transcriptome is thought to be predominantly noncoding^58^, comprising RNAs that regulate ribosome biogenesis (*e*.*g*., small nucleolar RNAs, snoRNAs)^59^, transcripts with putative nucleolar-architectural and chromatin-regulatory roles^60^, and a large cohort of long noncoding RNAs (lncRNAs) with yet-undefined functions^25^. Collectively, these transcripts arise from all three RNA polymerases and span lengths from 20 nucleotides^61^ to several kilobases. We attempted to capture these diverse species by using a non-poly(A)-selective, rRNA-depletion-based library preparation strategy similar to that established previously^62^. This yielded a catalog of *47S*-proximal RNAs that recapitulated much of the known nucleolar transcriptome, underscoring O-MAP-Seq’s precision and accuracy (**Figs. 3b–c, Supplementary Figs. 13–14**, and **Supplementary Table 4**). Of note, we observed conspicuous enrichment of every snoRNA detectable in our libraries (40 total, average 1.98 fold enrichment, *p=*0.011; FDR 0.05), the “transcribed spacer” domains within the *47S* pre-rRNA itself^38^, and nucleolar lncRNAs like SLERT^63^. As predicted, coding genes were broadly de-enriched, with only 940 (4.64% of detectable genes) exhibiting significant nucleolar enrichment (**Fig. 3b–c**; *p*_*adj*_ *= 0*.*009*; *FDR 0*.*05*). Moreover, isoform-level analysis revealed that most of these nucleolar-enriched, nominally protein-coding transcripts were in fact noncoding isoforms—most prominently, transcripts with retained introns (∼46% of enriched transcripts, comparted to ∼4% of the de-enriched pool, **Fig. 3d**). A similar intron-retention phenomenon has been recently observed in other subnuclear compartments^64^, suggesting that our observations represent *bona fide* biological regulation and not artifacts of the O-MAP-Seq pipeline. Likewise, the genomic localization and domain architecture of *47S* O-MAP-Seq-enriched transcripts suggest their validity as nucleolar RNAs. Nearly half (49.1%) of these transcripts are expressed from genes located within Nucleolar Associated chromatin Domains (NADs, **Fig. 3e**), compared to 7.7% of all genes. A similar enrichment of NAD-encoded transcripts was observed previously^25^, suggesting that these RNAs may be localized to the nucleolus cotranscriptionally. Nucleolar transcripts were also enriched for nearly every family of transposable element (TE), with the AluSg4 family of Alu SINEs being the most significant (**Fig. 3f**, *p*_*adj*_ = 2.22×10^−30^; *FDR 0*.*05*). This is consistent with previous work demonstrating that Alu repeats are particularly enriched in nucleolar- and lamina-associated RNAs^25^, and that these RNAs may play a role in stabilizing nucleolar architecture^65^. Finally, our *47S* O-MAP-Seq data also identified 3186 putative novel nucleolar transcripts (**Supplementary Table 4**). We confirmed by RNA-FISH that one such transcript, ENSG00000286147.1 (9.85-fold nucleolar enrichment; *p*_*adj*_ = 0.048), exhibited nucleolar and perinucleolar localization (69.77%, n=43; **Fig. 3g**). This further validates O-MAP-Seq’s accuracy and demonstrates its ability to discover novel RNAs within a target subcellular compartment.

Encouraged by these results, we next sought to apply O-MAP-Seq to a lower-abundance target with a previously uncharacterized RNA interactome. For this we chose the lncRNA *Xist*, the “master regulator” of mammalian X-chromosome inactivation (XCI)^45, 66-68^. Differentiated cells typically express only 100–200 copies of the *Xist* transcript^69^; these maintain epigenetic silencing by physically coating the inactive X-chromosome (Xi) and compacting it into a discrete subnuclear compartment^70^. The protein- and chromatin-interactions that enable *Xist* to drive this process have been extensively studied^66, 67^, but to our knowledge the RNA composition of the *Xist-*coated Xi compartment remains uncharacterized. We thus sought to elucidate the Xi transcriptome by O-MAP-Seq, using our validated *Xist-*targeting probe set, in mouse “Patski” cells^71^, and the optimized pipeline developed above (**Fig. 1d**). This revealed striking enrichment of the *Xist* transcript itself (29-fold enriched over input), and of lncRNAs *Jpx* and *Ftx* (average 12.1–fold enrichment), which are located from the same genomic neighborhood and which also contribute to XCI^67^ (**Fig. 3h**,*left*). Furthermore, we observed prominent enrichment of several X-linked genes that are known to escape XCI (*e*.*g. Mid1; Shroom4*)^68^, likely due to the capture of nascent transcripts expressed from the Xi (**Supplementary Fig. 15**). This was paralleled by the enrichment of the recently discovered noncoding RNA^72^ GM14636, suggesting that it too might escape XCI (**Fig. 3i**, *left*). RNA-

FISH analysis supported this hypothesis: though mono- and biallelic expression of GM14636 was variable in Patski cells, approximately 41.5% of observed foci co-localized with *Xist*, indicating penetrant expression from the Xi (**Fig. 3i**, *right*). Finally, we noted that *Xist* O-MAP transcripts were highly enriched for several classes of X-linked transposable elements—most notably, the L1MCb and L1MD1 subfamilies of LINE1 elements (**Fig. 3h**, *right*). These elements are overrepresented on the X-chromosome, and have been hypothesized to play a role in XCI^73^.

Intriguingly, *Xist* O-MAP-Seq also enriched several notorious chromatin-regulatory lncRNAs, including the imprinting regulator *Kcnq1ot1* (**Figs. 3h** and **3j**, *left*). Like *Xist, Kcnq1ot1* physically coats a Mb-scale domain near the site of its transcription, ultimately driving that domain’s heterochromatinization and silencing^74^. RNA FISH confirmed that *Kcnq1ot1* co-localizes with the *Xist* “cloud” at higher-than-expected frequency (14% of puncta, *n = 896*; **Fig. 3j**, *right*). This might indicate that *Xist* and *Kcnq1ot1* share a common pool of resources, or visit a shared subnuclear structure, that supports both of their epigenetic silencing functions.

### Mapping RNA-proximal genomic interactions with O-MAP-ChIP

We next examined if O-MAP could map the chromatin loci within a target transcript’s subnuclear compartment. As with O-MAP-Seq, our strategy relied on formaldehyde crosslinking and the capture of *in situ* biotinylated proteins to enrich nearby DNA, similar to Chromatin Immunoprecipitation (ChIP; **Fig. 4a**). We focused again on the lncRNA *Xist*, in mouse Patski cells. In differentiated cells like these, *Xist* is thought to physically coat the inactive X-chromosome^45^. Indeed, when we mapped our *Xist* O-MAP-ChIP data genome-wide, we observed robust enrichment along the entirety of chromosome X (**Fig. 4b**). Patski cells are a hybrid line derived from a cross between *M. musculus* and *M. spretus*, and they exhibit skewed XCI in which the *musculus* X-chromosome is constitutively inactivated^71^. This allowed us to quantify the allelic segregation of *Xist* O-MAP-ChIP reads, by measuring distinct SNPs between maternal and paternal alleles. Strikingly, the average allelic proportions across *Xist* O-MAP X-chromosomal peaks were highly specific to the inactive (*musculus*) allele (94%; **Fig. 4c**).

**Figure 4.**
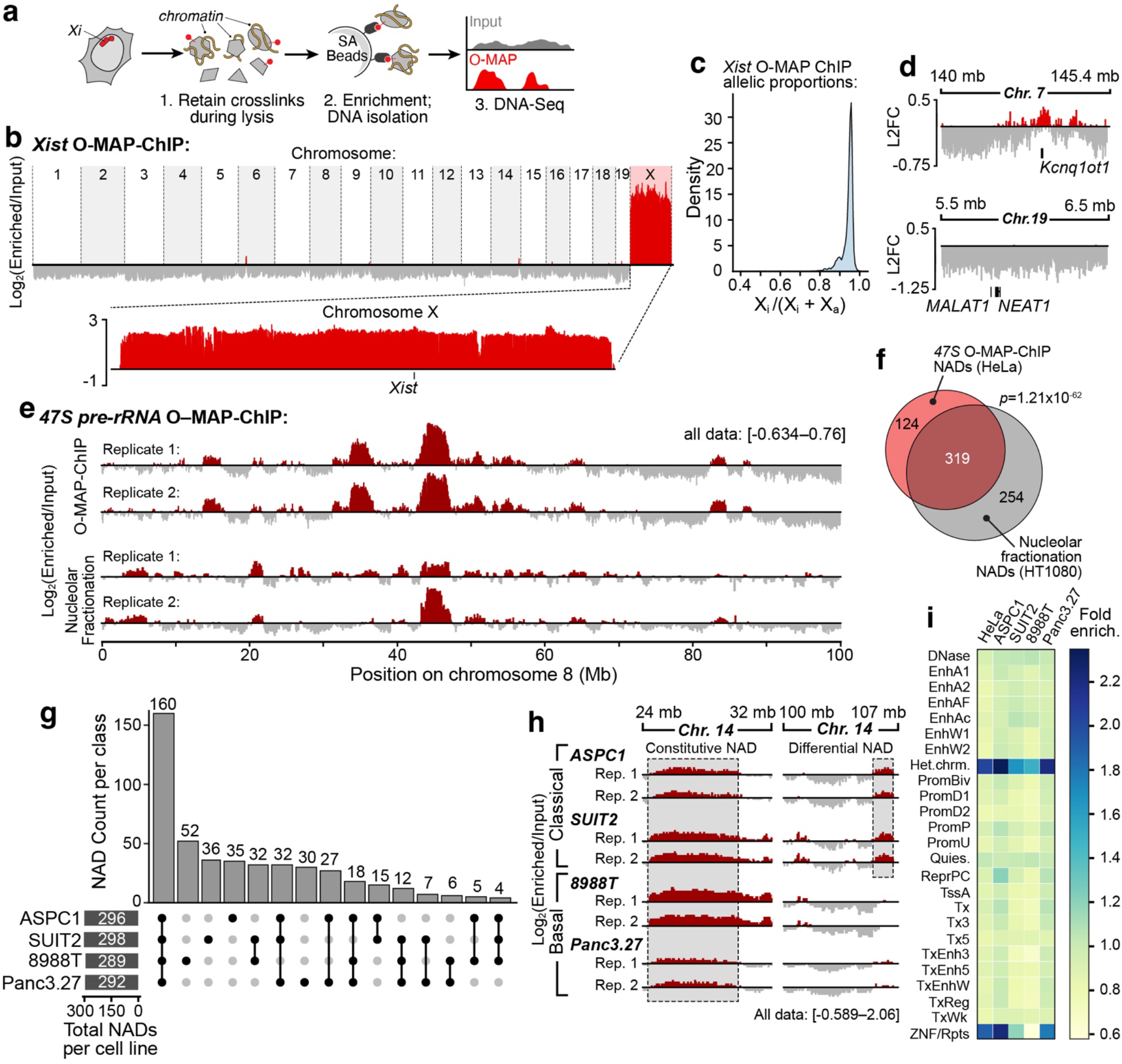
O-MAP-ChIP for probing RNA-proximal genomic loci. **a**, Approach. *In situ* biotinylated proteins are used to enrich nearby chromatin loci, which are then quantified by DNA-sequencing. **b–d**, O-MAP-ChIP characterization of *Xist* genomic interactions, in Patski cells. **b**, *Xist* O-MAP-ChIP predominantly labels the X-chromosome (*right*, and *inset*). Data for the entire mouse genome are shown; *Xist* genomic locus is noted below. **c**, *Xist* O-MAP-ChIP is specific to the inactive X-chromosome (Xi). Histogram of Allelic proportions for ChIP data, quantified using SNPs specific to the Xi and Xa^77^. **d**, Putative interactions between autosomal loci and the Xi. The *kcnq1ot1* locus—but not *MALAT1* and *NEAT1* loci—appear enriched in *Xist O-MAP-ChIP* data. L2FC: log_2_(Fold change, Enriched/Input) **e**–**i**, O-MAP-ChIP characterization of Nucleolar Associated Domains (NADs). **e**, 47S-targeted O-MAP in HeLa cells recapitulates the known human NAD architecture. Most of chromosome 8 is shown. Note higher reproducibility of O-MAP data. Nucleolar fractionation data taken from (Ref. ^78^). **f**, Conservation of NAD architecture between HeLa and HT1080 cells. **g**, Parallelized analysis of NAD architecture across four Pancreatic Ductal Adenocarcinoma (PDA) cell lines. Upset Plot summarizing NAD conservation, or lack thereof, between lines. The total number of NADs in each line appears nearly invariant. **h**, O-MAP-ChIP identifies NADs that are differentially regulated between Classical and Basal PDA subtypes. Examples of constitutive (*left*) and Differential (*right*) NADs on Chromosome 14 are shown. **i**, ChromHMM^79^ analysis reveals differential enrichment of chromatin signatures among HeLa and PDA cell line NADs.

Considerable evidence indicates that the *Xist* RNA makes direct chromatin contact exclusively with loci on the inactive X-Chromsome^75, 76^. Thus, we were curious about the autosomal peaks in our O-MAP-ChIP data which, while unlikely to be points of direct interaction with the free *Xist* transcript, might correspond to rare, *trans-* chromosomal contacts near the *Xist*-bound Xi. A similar profile of autosomal contacts has been observed by oligo-capture methods upon *Xist* overexpression, suggesting that O-MAP may be able to capture interactions for which other methods require signal amplification^80^. Notably, we observed one such putative *trans*-chromosomal interaction with the *Kcnq1ot1* locus (**Fig. 4d**, *top*), which is thought to be directly bound by the *Kcnq1ot1* RNA^74^. We also observed significant enrichment for this RNA by O-MAP-Seq (**Figs. 3h**,**j**), implying that the *Kcnq1ot1* locus—bound by its own transcript—is sometimes recruited near *Xist-*bound loci on the Xi. Importantly, the chromatin-regulatory lncRNAs *Malat1* and *Neat1*, which were highly enriched at the RNA level by *Xist* O-MAP-Seq (**Fig. 3h**) but which are not thought to bind their own loci^18^, were not enriched at the DNA level by *Xist* O-MAP-ChIP (**Fig. 4d**, *bottom*).

As a more challenging target, we next sought to use O-MAP-ChIP to profile nucleolar-chromatin interactions in HeLa cells, by targeting the *47S* pre-rRNA. Mammalian nucleoli are surrounded by megabase-scale chromatin structures termed Nucleolar Associated Domains (NADs) which comprise nearly half of all heterochromatin, and which are central to epigenetic programming^81^. Although NADs have been characterized by isolating and sequencing intact nucleoli^78^, this demanding approach has been challenging to apply in more than a handful of human cell lines. In contrast, *47S* O-MAP-ChIP enabled comprehensive, high-resolution maps of HeLa NADs from only 4×10^6^ cells, with approximately five days’ hands-on time (**Fig. 4e** and **Supplementary Fig. 16**). These O-MAP-ChIP data largely recapitulated the NAD architecture mapped by biochemical fractionation^78^, with nearly 72% overlap between the two data sets (*p*=1.21 × 10^−62^, Fisher’s exact test), even though they were acquired from different cell lines (**Fig. 4f**). O-MAP-ChIP also appeared markedly less noisy than fractionation-sequencing, with higher agreement between replicates (**Fig. 4e**).

NAD architecture is almost universally remodeled in cancer, potentially driving epigenetic and transcriptional changes that facilitate oncogenesis^11, 81, 82^. Yet, the functional impact of this dysregulation has been difficult to assess without robust methods for characterizing NAD architecture across cancer types. To demonstrate how O-MAP might facilitate this analysis, we applied *47S* O-MAP-ChIP across four Pancreatic Ductal Adenocarcinoma (PDA) cell lines, systematically interrogating NAD organization across both the “classical” (ASPC1 and SUIT2 lines) and “basal” (8988T and Panc3.27 lines) PDA subtypes^83^ (**Fig. 4g**). Although nucleolar morphology differs markedly between PDA subtypes^83^, *47S* O-MAP-ChIP revealed a set of invariant nucleolar-genomic interactions (160 NADs) conserved across all cell lines, suggesting a core PDA NAD architecture. These conserved domains comprise the most abundant NAD class (approximately 55% of NADs), though sizeable groups of cell line-specific domains (30–52 NADs; 10–18%), or domains uniquely absent from a cell line (4–32 NADs; 1.4–11%) were also observed. We also observed NADs unique to each PDA subtype— 15 and six NADs in classical and basal, respectively (**Figs. 4g**,**h**). These variable domains may be particularly important for PDA, in which epigenetic regulation is thought to play a key role during oncogenesis and subtype specification^84^. ChromHMM analysis to identify cell line specific NAD epigenomic signatures^79^. This showed clear differences in zinc-finger/repeats (ZNF/Repeats) and heterochromatin domains between NADs of different cell types, with a profound loss of ZNF/Repeats in 8988T NADs (**Fig. 4i**). To our knowledge, any one of these data sets would represent the highest-resolution map of human NADs reported to date. Furthermore, these data— combined with our nucleolar proteomic (**Fig. 2**) and transcriptomic analyses (**Fig. 3a**–**g**)—demonstrate O-MAP’s capacity for parallelized, “multi-omic” dissection of RNA-scaffolded compartments using a common workflow.

### O-MAP is portable across specimen types and target RNAs

A chief limitation of genetically encoded proximity-biotinylation methods is that they require the generation of a custom-built transgenic line for each new cell culture model, tissue, or organism under interrogation^20^. Because O-MAP doesn’t require genetic manipulation to program its spatial targeting, it may enable rapid parallelization. We demonstrated this using O-MAP to profile the 47S pre-rRNA across a panel of cultured mammalian cell lines. In all cases, we used the same hybridization- and *in situ*-labeling conditions optimized in HeLa cells, and analogous (species-specific) 47S*-*targeting probes (**Fig. 1b, Supplementary Table 1**). This revealed O-MAP to be remarkably modular, catalyzing precise, nucleolar-targeted biotinylation in every line tested (ASPC1, SUIT2, 8988t, Panc3.27, A375, HEK 293T, U2OS, and MEFs, **Fig. 5a** and **Supplementary Fig. 17**). Notably, we observed robust labeling irrespective of nucleolar volume, density, or morphology—factors that can limit affinity-purification- and fractionation-based approaches. This modularity enabled us to perform parallelized discovery of nucleolar-genomic interactions across multiple cell lines (**Fig. 4e**–**i**), which would be cumbersome by established methods.

**Figure 5.**
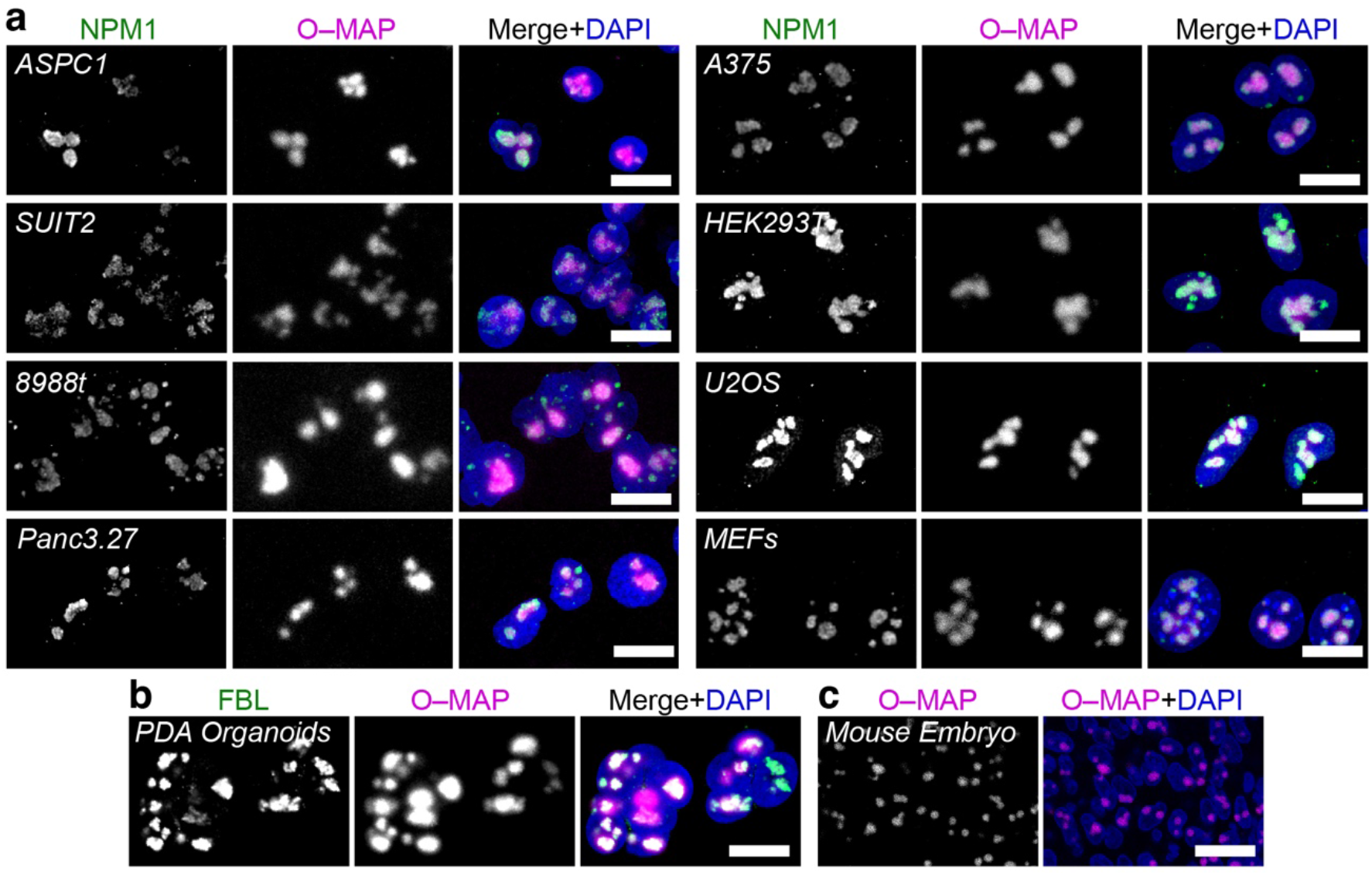
O-MAP is readily ported across specimen types. **a**, 47S-O-MAP in cultured mammalian cell lines. NPM1 immunofluorescence denotes nucleoli. All human-derived lines used the same probe set and hybridization conditions as HeLa cells (**Figs. 1–4**); MEFs used an analogous mouse-targeting probe set. **b**, 47S O-MAP in human patient-derived pancreatic ductal adenocarcinoma (PDA) organoids. *FBL* immunofluorescence denotes nucleoli. **c**, 47S O-MAP in cryo-preserved mouse tissue slices. All scale bars: 20 µm.

Next, we tested if O-MAP might be portable to specimens for which transgenic approaches would be more challenging or intractable, using patient-derived PDA tumor organoids^85^ and fixed mouse embryo tissue sections as models. Although each sample-type required modest re-optimization of hybridization- and *in situ*-labeling conditions, we were able to generate robust and specific 47S-targeted O-MAP in both (**Fig. 5b**,**c**). This demonstrates that O-MAP is extensible beyond two-dimensional cell culture, and implies its potential application for RNA-interaction discovery in clinically relevant contexts^85^.

We next examined O-MAP’s portability to different RNAs, by targeting an array of transcripts with diverse lengths, expression levels, sequence composition, biogenesis pathways, and localization. For each new target, probes were designed using the OligoMiner pipeline^34^, and the optimal *in situ* biotinylation time was determined empirically via a labeling time course (**Supplementary Table 5**). Encouragingly, in each case, O-MAP yielded prominent *in situ* biotinylation that recapitulated the target’s known subcellular localization. Our Probe Validation Assay (**Fig. 1d**) further confirmed the RNA-targeting accuracy of the probe pool: in all cases *in situ* biotinylation and RNA-FISH signals exhibited a high degree of overlap (**Fig. 6a**). This was observed with highly abundant transcripts like the chromatin-regulatory lncRNAs *MALAT1* and *NEAT1*, with modestly expressed targets like the *WDR7* mRNA^86^, and with low-abundance RNAs like *Firre* and *Kcnq1ot1*. We note that obtaining robust O-MAP at low-abundance targets required large primary probe pools (100–500 oligos) to bolster the number of HRP molecules recruited to the target transcript, as well as longer labeling times (60–120 minutes; **Supplementary Table 5**).

Finally, we hypothesized that O-MAP could be applied to RNA domains that uniquely mark nascent transcripts, thereby enabling analysis of an RNA’s subnuclear microenvironment during the early stages of its biogenesis. Having established this strategy with Transcribed Spacer domains on the 47S pre-rRNA (**Fig. 1** and **Supplementary Fig. 3**), we sought to generalize the approach with nascent Pol II transcripts, by targeting introns. As a first test case we selected *Xist*, since its nascent transcripts (“pre-*Xist*”) denote the X-chromosome

**Figure 6.**
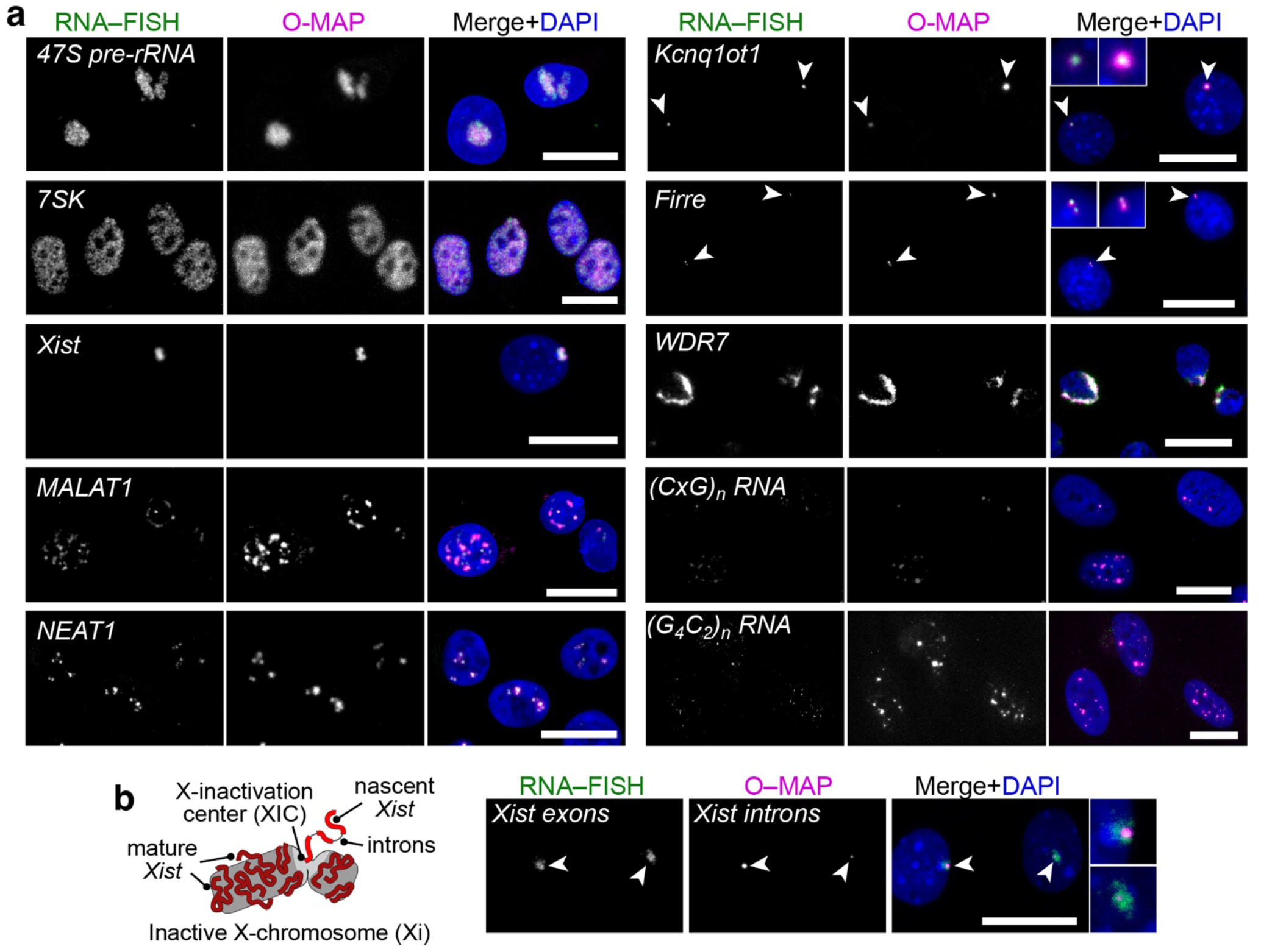
O-MAP is broadly applicable to different RNA targets. **a**, O-MAP Probe Validation Assay (**Fig. 1d**) applied to a compendium of target transcripts. Note conspicuous overlap between RNA–FISH (*green*) and O-MAP (*magenta*) signals. Images from HeLa (*47S, 7SK, MALAT1, NEAT1*), Patski (*Xist, Kcnq1ot1, Firre*), patient-derived fibroblast^86^ (*WDR7*), and U2OS cells (*(CxG)n* and *(G4C2)n RNAs*). *Firre* is expressed from a single-copy transgene; (CxG)n and (G4C2)n RNAs are artificial constructs under doxycycline-inducible expression^87^; all other targets are endogenous transcripts. Insets show zoomed-in sections of the same images, to highlight signal overlap. **b**, O-MAP at nascent transcripts probes subnuclear neighborhoods. *Left:* Although mature *Xist* coats the entire inactive X-chromosome (Xi), nascent *Xist* transcripts uniquely denote the X-inactivation center (XCI). *Right:* O-MAP targeting *Xist* introns induces biotinylation at confined foci within the Xi “cloud.” All scale bars: 20 µm.

Inactivation Center (XIC), a specialized chromatin domain that is critical for XCI initiation^45, 67^ (**Fig. 6b**, *left*). As expected, pre-*Xist* O-MAP induced small, bright biotinylation puncta localized within larger “clouds” of *Xist* RNA-FISH, suggesting precise biotinylation of a small subcompartment within the Xi (**Fig. 6b**). Such subcompartment labeling demonstrates that O-MAP can distinguish between populations of a target RNA’s isoforms (*i*.*e*., mature vs. nascent), and suggests a general strategy for precisely probing subnuclear micro-compartments that would be challenging to purify biochemically.

## Discussion

In this work, we have developed a nearly universal method for applying proximity-labeling to RNA, by co-opting the mechanics of RNA-FISH to precisely deploy biotinylating enzymes to a target transcript without genetic modification. This approach, O-MAP, supports a robust toolkit for elucidating the proteins (O-MAP-MS), transcripts (O-MAP-Seq), and genomic loci (O-MAP-ChIP) near an RNA of interest. We believe this toolkit holds considerable advantages over established RNA interaction-discovery methods, offering superior precision, biological context, flexibility, ease of use, and cost. Moreover, we anticipate that O-MAP’s ability to probe higher-order interactions within a transcript’s subcellular “neighborhood” will enable unprecedented analysis of RNA-mediated compartmentalization^6, 7^, a powerful new approach for spatial cell biology.

The most effective RNA interaction-discovery methods use antisense oligonucleotides to pull down a target RNA *ex vivo*^14-17^, or transgenic manipulation to localize proximity-labeling enzymes to that RNA in live cells^21, 27, 28^. Compared to oligo pulldown-based approaches (*e*.*g*. ChIRP, ChART, RAP)^14, 17, 88^, O-MAP offers more precise optimization, higher sensitivity, and broader biological context. A chief limitation of these pulldown-based techniques is their inability to optimize the targeting precision of the capture-oligo pool^14, 18^, leading to off-targeting noise. O-MAP overcomes this using a straightforward validation assay that rapidly identifies and eliminates off-targeting probes (**Figs. 1d**–**e**, and **6**). Furthermore, oligo pulldown techniques capture their target RNA in crude cell lysates, where artifactual interactions with abundant, promiscuous RNA-binding proteins can confound analysis^19^. Overcoming this issue typically eliminates all but the target RNA’s most direct interactors^14^, thus eliminating higher-order biological context. Moreover, the inefficiency of target capture, typically necessitate large inputs (>10^8^ cells per replicate)^14, 17, 88^. In contrast, because O-MAP enzymatically amplifies the number of biotins per oligo probe, and enables interacting molecules to be directly captured (rather than co-purified by pulldown), it is markedly more efficient. Of note, our multi-omic analysis of HeLa nucleoli (**Figs. 2, 3a**–**g**, and **4e**– **i**) collectively required fewer cells than would a single replicate of ChIRP-MS^17^. This higher efficiency, and the ease with which O-MAP is deployed to different specimens (**Fig. 5**), could enable parallelized analysis across tissues, cell types, or other experimental variables (*e*.*g*., **Fig. 4g–i**), which would be unwieldly using oligo-capture methods. And unlike those methods, proximity-biotinylation approaches like O-MAP are uniquely suited to probe dynamic, higher-order interactions that are too transient or fragile to survive a pulldown.

Compared to RNA-targeted live-cell proximity-labeling methods, O-MAP is more spatially precise, more applicable across transcripts and specimens, and is simpler to design and execute. Live cell approaches (*e*.*g*., RapID, MS2/Cas13-APEX, CRUIS) assemble artificial complexes between the labeling enzyme and its RNA target, by transgenically overexpressing these components fused to localization sequences or binding motifs^21, 27, 28, 89^. Although such transgenic approaches have had some success, they must contend with substantial background labeling from unbound enzymes^27, 28^ (**Fig. 1c**), and with artifacts from overexpressing the target RNA outside of its native context^90^. These issues are particularly problematic with low-abundance transcripts. In contrast, O-MAP doesn’t require transgenic manipulation to control enzyme localization. The sequences, abundances, and hybridization parameters of its HRP-targeting oligos can be explicitly controlled to minimize off-targeting^35^, and residual, mislocalized probes can be removed by stringent washing. This strategy enabled endogenous RNAs—even low abundance transcripts—to be biotinylated with nearly undetectable background (**Figs. 1b** and **6**). It furthermore enables O-MAP analysis in contexts where genetic manipulation would be challenging or altogether impossible, including pre-fixed tissue samples (**Fig. 5c**). Applying O-MAP in samples like these, such as clinical isolates, may provide a powerful avenue for biomarker and therapeutic target discovery.

O-MAP is conceptually similar to other methods that use affinity reagents, rather than transgenic expression, to localize proximity-labeling enzymes. Several established ‘omics tools (*e*.*g*., EMARS, SPPLAT, BAR, TSA-Seq, and TSA-MS)^26, 40, 91-93^ use antibodies to deploy HRP to a target protein *in situ*, essentially adapting an immunofluorescence workflow for novel interaction discovery. A similar tactic underlies RNA-TRAP^30^ and HyPro^31^, which use Digoxigenin (DIG)-modified oligos to recruit HRP to a target RNA—either via an anti-DIG antibody-conjugate^30^ or linked to a custom-made DIG-binding protein^31^. In our experience, these hapten-recruitment strategies are markedly noisier than O-MAP’s “landing-pad” approach (**Supplementary Fig. 2**), and heir lack of modularity precludes strategies like our Probe Validation Assay (**Figs. 1d–e** and **6**), a vital optimization tool. Moreover, unlike the current implementation of HyPro, O-MAP is compatible with quantitative tandem mass-tag-based proteomics (**Fig. 2**), enables ChIP-like genomic interaction discovery (**Fig. 4**), and can be used with conventional formalin-fixed tissues (**Fig. 5**). O-MAP is also considerably cheaper, using inexpensive, chemically unmodified primary oligos, in lieu of DIG-conjugated probes. And critically, O-MAP uses entirely off-the-shelf parts, and doesn’t require the synthesis of custom proteins.

Our work also illustrates O-MAP’s unique ability to discover compartment-level interactions that are opaque to current methods. For example, while 7SK’s direct binding partners are well established^49, 50^, our O-MAP-MS analysis represents, to our knowledge, the first proteomic characterization of this enigmatic RNA’s subnuclear compartment (**Fig 2f**,**g**). O-MAP-Seq likewise enabled first-of-its-kind characterization of the *Xist-* proximal transcriptome, revealing a new XCI-escape gene and the unanticipated co-compartmentalization of *Xist* with other chromatin-regulatory RNAs (**Fig. 3h**–**j**). Notably, the novel interaction between *Xist* and *Kcnq1ot1* (**Fig. 3h**,**j**) appears to occur at distances that would be intractable by most RNA-RNA interaction-discovery tools, which predominantly query direct base-pairing contacts^94^. This interaction is also paralleled in our O-MAP-ChIP analysis, which revealed a cohort of putative long-range interactions between the inactive X chromosome and autosomal loci (**Fig. 4d**). *Trans*-chromosomal contacts of this sort are notoriously challenging to probe using conventional, genomics tools^95^. This suggests that O-MAP-ChIP might enable characterization of subnuclear compartments at currently unattainable distance scales, providing powerful new insight into higher-order genome organization. O-MAP-ChIP also enabled a parallelized analysis of NAD structure across cell lines, revealing both conserved and variable elements of NAD architecture (**Fig. 4e–i**). Extending this analysis to other lines, and complementing it with O-MAP-MS and O-MAP-Seq, will enable an unprecedented molecular characterization of nucleolar architectural remodeling during oncogenesis^82^.

We anticipate that O-MAP will be applicable to nearly any target RNA, and in any biological setting, for which RNA-FISH can be performed. However, we note some of the technique’s current limitations. O-MAP will likely be challenging with difficult FISH targets (*e*.*g*., small, low-abundance RNAs), requiring more advanced probe designs to amplify signal^35^. Moreover, formaldehyde fixation—though a mainstay of high-resolution imaging—can induce aberrations that perturb the local subcellular structure^96^, and which might be improved using alternate fixation strategies. Finally, like all peroxidase-based proximity-labeling strategies, O-MAP’s labeling chemistry identifies all molecules within its labeling radius, including both the target’s direct and nearby interactors. Future development of O-MAP’s probe design and labeling chemistry will enable more precise control over its labeling radius, allowing “shells” of RNA-interactions to be probed independently. Regardless, in its current implementation we believe that O-MAP’s ability to elucidate RNA interactions *in situ* overcomes a technical roadblock that has long challenged the RNA community. Given the extraordinarily broad scope of cellular functions performed by RNA^97^, this advancement will catalyze fundamental discoveries into countless biological phenomena.

### Data availability and accession codes

Mass spectrometry data will be deposited to ProteomeXchange. RNA- and DNA-Sequencing data have been deposited to the Gene Expression Omnibus under SuperSeries accession number GSE217566. Raw imaging files and custom code used in this work is available upon request.

## Acknowledgements

We thank J. Gianopulos, C. Hsu, C. Anderson, and J. Cabarrus for general technical assistance; A. Jain, M. Das, R. Akilesh, P. Valdmanis, and S. Smukowski for the kind donation of materials; T. Moss for rRNA computational tools; N. Peters, K. Collins, M. Soruco, E. Hacisüleyman, and A. Fenix for imaging assistance; A. Bertero, S. Attar, E. Nichols, for help with cell-based models; H-T. Lau and M. G. Golkowski for mass-spec assistance; G. Yardimci, G. Bonara, A. Mishra, R.D. Hawkins, and W. Noble for sequencing assistance; Y. Sancak, R. Gardner, and members of their labs for thoughtful discussions and critiques. This work was supported by National Institutes of Health Grants 1R01GM138799-01 and T32GM007750, a Safeway Albertsons Early Career Award in Cancer Research, a Brotman Baty Institute Catalytic Collaborations Award, the UW Royalty Research Fund (RRF), and the UW Student Technology Fund. E.E.K. was supported by the NSF DEB2016186 and the AHA 902616. Imaging at the UW Keck Center was supported by NIH S10 OD016240 and the UW Student Technology Fee. NGS data analysis was facilitated through the use of advanced computational, storage, and networking infrastructure provided by the Hyak supercomputer system and funded by the STF at the University of Washington.

## Author Contributions

A.F.T., E.E.K. and D.M.S. conceived of, designed, and analyzed all experiments, which were principally performed by A.F.T and E.E.K., assisted by D.Q.L. and M.K. Mass-spectrometry was assisted by R.F., C.D.M., S-E.O., and D.K.S. S.K. assisted with PDA- and organoid experiments. X.D. and C.M.D. assisted with all *Xist*- and *Firre*-related experiments. B.J.B. and E.A.H. assisted with oligonucleotide probe design. D.M.S. directed research. All authors contributed to writing.

## Competing Financial Interests

The authors declare competing financial interests. A.F.T, E.E.K, B.J.B, and D.M.S. have filed for a patent concerning the use of oligonucleotide-directed proximity-labeling to elucidate and visualize subcellular interactions *in situ*.

## ONLINE METHODS

### Cultured cells, tissue sections and organoids

HeLa, HEK 293T, U2-OS (*gifts from Dr. Y. Sancak, UW*), patskis and female mouse embryonic fibroblasts (fMEFs, *both gifts from Dr. C. Disteche, UW*), and SUIT2 cells (*a gift from Dr. S. Kugel, Fred Hutch*) were cultured in High Glucose DMEM with Pyruvate (Thermo Fisher; 11995073), supplemented with 10% (v/v) Fetal Bovine Serum (FBS, Thermo Fisher; 26140079), 100 units/mL penicillin and 100 µg/mL streptomycin (Thermo Fisher; 15140122), and 1x GlutaMAX (Thermo Fisher; 35050061). For Transgenic (CxG)_n_ and (G_4_C_2_)_n_ U-2 OS cells^87^ (**Fig. 6a**, *gifts from A. Jain, MIT*) qualified tetracycline-free FBS (Gibco; 26140079) was used. For these lines, transgenic RNA expression was induced by addition of doxycycline to a final concentration of 1 mg/mL for 24 hours prior to cell fixation. A375 (*a gift from Dr. J. Scott, UW*), 8988T, ASPC1, and Panc 3.27 cells (*gifts from Dr. S. Kugel*) were cultured in RPMI 1640 media (Thermo Fisher; 11875093), supplemented with 10% (v/v), FBS, 100 units/mL penicillin, and 100 µg/mL streptomycin. Patient-derived lymphoblast cell lines^86^ (*a gift from Dr. P. Valdmanis, UW*) were cultured in IMDM (Thermo Fisher; 12440053), supplemented with 20% (v/v) FBS, 100 units/mL penicillin, 100 µg/mL streptomycin, and 2.5 µg/mL amphotericin B. In all cases, cells were maintained at 37°C, under 5% CO_2_. Cell lines were authenticated by STR testing (ATCC), when possible.

For imaging experiments, cells were cultured in two-well Lab-Tek borosilicate glass #1.0 chambers (Thermo Fisher; 155380). To improve HEK293T adherence, chambers were treated with gelatin (0.5% (w/v), in water, Sigma; G7765) for 30 minutes at 37°C, prior to plating. For biochemistry, proteomic, and high-throughput sequencing experiments, cells were cultured in six-well plates. When necessary, material from multiple wells was harvested and merged into a single lysate, as described below.

Human Pancreatic Ductal Adenocarcinoma organoids (**Fig. 5b**) were prepared as described^85^. Mouse tissue sections (**Fig. 5c**, *a gift from Dr. E. Nichols, UW*) were prepared from day E13.5 embryos (C57BL/6J wild type mice; Jackson Laboratory) by drop fixing in 4% (v/v) formaldehyde, followed by sucrose equilibration, and thin-sectioning in OCT compound (VWR 25608-930. Cryosections (approximately 10 µm) were prepared at the UW Histology and Imaging Core and stored on glass slides at –80°C until use.

### O-MAP probe design and synthesis

Probes targeting the human 47S pre-rRNA ITS1 domain were taken from (Ref. 37). All other probes were designed using OligoMiner pipeline^34^ using the following settings. The blockParse script was run using the settings: -l 30 -L 37 -t 42 -T 47 -s 390 -F 40. Bowtie2 was used with settings: -U --no-hd -t -k 100 --very-sensitive-local -S. The outputClean script was run with the -u argument; the structureCheck script was run with the settings: -F 40 -s 390 -m dna1998 -T 42. K-mer filtering was performed in Jellyfish version 2.2.10, using a Jellyfish file for the corresponding genome (human or mouse), and using the kmerFilter function with the -m 18 and -k 1 arguments. Jellyfish files were generated for each genome assembly (hg38 for human; mm10 for mouse), using a hash size set to the appropriate size of the genome assembly^34^. For example, the command -s 3300M -m 18 -o hg38_18.jf --out-counter-len 1 -L 2 hg38.fa was used to generate the 18mer dictionary for hg38. For most targets, all probes that passed this final filtering step—typically 10–150 probes per target—were used. For *kcnq1ot1*, a set of 200 k-mer-filtered probes were used (**Supplementary Table 1**).

For the O-MAP Probe Validation Assay (**Fig. 1d**), probes were divided into sub-pools along the length of the target RNA, appended on their 3’ termini with SABER1 or SABER2 “Landing-pad” sequences^35^ (**Supplementary Table 1**). Once probes were validated, the complete pool was reformulated, appended with SABER1 and used for O-MAP alone.

Probe sets consisting of fewer than 20 probes were ordered as individual oligos (Sigma; 0.025–0.05 µg synthesis scale, standard desalting), and further purified from preparative polyacrylamide gels, as described previously^98^. Purified oligos were resuspended in nuclease-free water, quantified by UV-vis spectroscopy, pooled to a final aggregate concentration of 20 µM and stored at –20°C. Probe pools requiring more than 20 oligos were purchased as oPools (IDT; 50 pmol/oligo scale, unmodified), and dissolved to approximately 100 µM in nuclease free-water. 20 µL were desalted using the Oligo Clean & Concentrator Kit (Zymo Research; D4060), following the manufacturer’s instructions. Pools were quantified by UV-vis spectroscopy, diluted to 5 µm and stored at – 20°C. Fluorophore-conjugated secondary probe used in RNA-FISH (“SABER2–AF647”, **Supplementary Table 1**) was purchased from IDT (100 nmol scale; HPLC purification), resuspended to 100 µM in nuclease-free water, and stored in a light-tight container at –20°C. The HRP-conjugated secondary probe (“SABER1–HRP,” **Supplementary Table 1**) was purchased from Bio-Synthesis, resuspended to 10 µM in resuspension buffer (10mM NaH_2_PO_4_, 150 mM NaCl, pH 7.2), allotted into 10 µL single-use aliquots, flash-frozen and stored at – 20°C.

In some cases, the RNA-FISH signal was amplified by extending the FISH probe subpool with concatemers of additional “SABER2” Landing-pads (**Supplementary Table 1**). These were enzymatically added via the Polymerase Exchange Reaction (PER), essentially as described^35^. Briefly, pooled probes (5 µM, aggregate) were incubated with 0.5 µM Template hairpin and 0.1 µM Clean.G DNA hairpin (IDT; **Supplementary Table 1**), in 10 mM MgSO_4_, 0.6 mM each ATP, CTP, and TTP, 4 U *Bst* 2.0 DNA Polymerase (NEB; M0537), 1x PBS, in a final volume of 50 µL. Reactions were incubated at 37°C for two hours in a thermocycler, heat-inactivated at 80°C for 20 minutes, and chilled on ice. The resulting PER-extended oligos were then purified with OligoClean and Concentrator Kits, eluting into nuclease-free water, and their length was examined on denaturing 10% Polyacrylamide TBE-Urea gels, stained with SYBR-Gold (Thermo Fisher; S11494).

### O-MAP core protocol

The following protocol was used for omics-scale O-MAP, using cells grown in six-well dishes (3.5×10^5^ cells/well; plated one day prior to harvest). For imaging-only experiments, cells were plated at 7×10^4^ cells per well, in two-chamber LabTeks. In all cases, RNase-free reagents and manipulations were used throughout.

The core O-MAP workflow is split over two days. The first day comprises fixation, permeabilization, peroxidase inactivation, and primary probe hybridization; the second day comprises secondary probe hybridization, an optional endogenous biotin blocking step, and *in situ* biotinylation. Thereafter, the protocol varies depending on the endpoint assay—imaging, proteomics, RNA-Seq, or ChIP-Seq.

### O-MAP Day 1

All manipulations were performed at room temperature, unless noted. Cells were washed briefly with 1x Ca- and Mg-free DPBS (Thermo Fisher; 14190250) and fixed with freshly prepared 2% (v/v) formaldehyde (Electron Microscopy Sciences; 15710) in 1x PBS (Sigma; 6506), for 10 minutes without agitation. The formaldehyde solution was aspirated and the crosslinking reaction terminated by two washes with 250mM glycine in 1x PBS, five minutes each, with gentle rocking (3 rpm on a platform rocker). Cells were briefly washed with DPBS, permeabilized with 0.5% (v/v) triton-X 100 in PBS (10 min; gentle rocking), and washed three times with DPBS. Next, to inactivate endogenous peroxidases, samples were treated with 0.5% (v/v) H_2_O_2_, in 1x PBS, for 10 minutes with gentle rocking, and washed twice with PBS. Samples were then equilibrated in Formamide Wash Buffer (10–40% (v/v) deionized formamide (Thermo Fisher; AM9342); 2x SSC (Thermo Fisher; AM9765); 0.1% (v/v) Tween-20), for five minutes with gentle rocking. The formamide concentration was matched to the primary probe hybridization mix, as determined by the binding parameters of the primary probe pool (**Supplementary Table 5**). This buffer was then aspirated, and each sample was treated with 115 µL of Probe Mix (0.1 µM primary oligo probe pool, in 1x Hybridization Buffer: 10–40% deionized formamide; 2x SSC; 0.1% (v/v) Tween-20; 10% (w/v) dextran sulfate (SIGMA; D8906); in nuclease-free water) and this mix was gently spread over the sample by covering with a clean, 30 mm diameter #1.5 thickness glass cover slip (Bioptechs; 30-1313-03192). A 2x SSC-soaked kimwipe was placed between the wells to maintain humidity during hybridization. Plates were then sealed with Parafilm and incubated without agitation for 8 hours at 37°C or 42°C, depending on the probe set (**Supplementary Table 5**).

### O-MAP Day 2

Following primary hybridization, coverslips were removed and cells were washed three times with pre-warmed 30% Formamide Wash Buffer, 10 minutes per wash, at 37°C with gentle rocking. For imaging experiments, this was followed by a blocking step to mask endogenous biotinylated proteins, as described below (*see* “O-MAP Imaging”). In all cases, subsequent manipulations were carried out in the dark, to avoid photooxidation of the HRP conjugate. Each well was treated with 115 µL O-MAP Secondary Probe Mix (100 nM SABER1–HRP oligo (**Supplementary Table 1**), in 30% Formamide Hybridization Buffer), and covered with a clean coverslip. Samples were incubated for 1 hour at 37°C, with gentle rocking. Coverslips were then removed, and samples were washed four times with PBST (0.1% (v/v) Tween-20 in 1x PBS), 15 minutes per wash, with gentle rocking. Buffer was aspirated, and *in situ* biotinylation induced by addition of 1 mL Labeling Solution (0.8 µM biotinyl-tyramide (Sigma; SML2135), 1 mM H_2_O_2_, 1x PBST), and incubation at room temperature. Labeling times varied between RNA targets—ranging from 1 second (**Fig. 2h**, *top left*) to 120 minutes (**Fig. 2h**, *bottom right*)—and were determined empirically using the O-MAP Imaging assays described below. In all cases, biotinylation was halted by addition of Sodium Azide and Sodium Ascorbate (10 mM each, final) in 1x PBST, for three washes of five minutes each.

### O-MAP imaging

For imaging experiments, the background signal from endogenous biotinylated proteins was blocked after the secondary probe hybridization step. Briefly, samples were washed three times in 1x PBST and incubated in pre-blocking solution (1% (w/v) nuclease-free BSA (VWR; 97061-420) in 1x PBST) at room temperature for 30 min with gentle rocking. Samples were then blocked with 1 mL of Neutravidin Blocking Solution (10 µg/mL neutravidin (Thermo Fisher; 31000), 1% (w/v) nuclease-free BSA, in 1x PBST) for 15 min with gentle rocking at room temperature, and washed three times with PBST. To saturate free streptavidin binding sites, samples were next treated with 10 µg/mL D-biotin (Thermo Fisher; B20656) in 1x PBST, for 15 minutes with gentle rocking, followed by three washes with room temperature PBST. Thereafter, *in situ* biotinylation and quenching proceeded as described above, using 50 µL volumes for primary and secondary hybridization buffers. After biotinylation and quenching, samples were stained with 1 mL 1 µg/mL neutravidin-DyLight 550 conjugate (Thermo Fisher; 84606), in 1% BSA pre-blocking solution, for one hour at room temperature with gentle rocking, followed by three washes with 1x PBST. Samples were counterstained with DAPI (5 µg/mL, in 1x PBST) and imaged immediately, or were mounted in Vectashield (Vector Labs; H-1900-10) and stored at 4°C.

### O-MAP-MS

O-MAP-labeled cells (approximately 5.5×10^6^ cells per replicate; three replicates per experimental condition) were harvested by scraping into 1x PBST, supplemented with 10 mM Sodium Azide. After pelleting by centrifugation at 800 x *g* for 10 minutes at 4°C, remaining buffer was aspirated and the pellets were flash frozen in liquid nitrogen and stored at –80°C until use. All subsequent steps were performed at room temperature, unless noted. Cell pellets were lysed in 800 µL of MS Lysis Buffer (4% (w/v) SDS in 1x PBS, with 10 mM Sodium Azide, 1x Halt EDTA-Free Protease Inhibitor Cocktail) for five minutes. Samples were then sonicated using a Branson Digital Sonifier 250 outfitted with a double stepped microtip (Emerson Industrial Automation) at 10–12 Watts for 30 seconds (0.7 s on; 1.3 s off) for one cycle. Lysates were clarified by centrifugation at 15,000 x *g* for 10 minutes and soluble protein was quantified using the Pierce BCA Protein Assay Kit (Thermo Fisher; 23225). For each sample, 300 µg of protein was diluted to 1% SDS by the addition of three volumes Dilution Buffer (1x PBS, supplemented with 10 mM Sodium Azide and 1x Protease Inhibitors). Protein samples were then reduced with TCEP (Thermo Fisher; 77720, 10 mM final concentration) for 60 minutes with gentle rotation. To alkylate free thiol groups, samples were treated with iodoacetamide (Sigma; I1149, 20 mM final concentration) and rotated for 60 minutes at room temperature, before quenching by addition of DTT (5 mM final) and incubation for 15 minutes. For streptavidin pulldown, to each sample was added 100 µL Pierce Streptavidin Magnetic Bead slurry (Thermo Fisher; 88817) that had been equilibrated in Diluted Lysis Buffer (1% (w/v) SDS, 1x PBS, 10 mM Sodium Azide, 1x Protease inhibitors). Samples were rocked end over end for two hours at room temperature, and streptavidin beads were then washed with the following buffers (5 minutes per wash; rocking end over end at room temperature): (1–2) Two washes in Diluted Lysis Buffer, (3) 1x PBS (4–5) Two washes in KCl Buffer (1 M KCl in 1xPBS), (6–7)Two washes in Urea Buffer (2 M Urea in 1xPBS), (8–9) Two washes in 200 mM EPPS (pH 8.5). Beads were then resuspended in 15 µL 200 mM EPPS (pH 8.5), and bound proteins were eluted by on-bead proteolytic digestion, as follows. Lysyl endopeptidase (Lys-C, Fujifilm Wako; 121-05063) was added at a ratio of 1 µg of enzyme per 100 µg of input protein, and samples were incubated for three hours at 37°C, with shaking at 500 rpm. Trypsin (Thermo Fisher; 90057) was then added at a ratio of 1 µg of enzyme per 100 µg of input protein, and digestion continued overnight at 37°C, 500 rpm shaking.

### O-MAP-MS LC-MS/MS Sample Preparation

Eluted peptides were labeled with TMTpro reagents using established protocols^47^. Briefly, eluted peptides were supplemented to with acetonitrile to a final concentration of 30% (v/v), in 200 mM EPPS buffer (pH 8.5). TMTpro reagents in 100% anhydrous acetonitrile were then added to each sample at approximately a 2.5:1 (w/w) excess. The labeling reaction was allowed to continue for 1.5 hours, quenched with 5% hydroxylamine, and the labeled peptides were mixed. Pooled peptides were then dried by vacuum centrifugation. Dried, labeled peptides were resuspended in 100 µl of (5% acetonitrile, 1% formic acid) and cleaned using in-house assembled stage-tips^99^. Pooled peptides were eluted in (70% acetonitrile, 1% formic acid). Eluates were then dried to completion and stored at -80°C until analyzed by LC-MS/MS.

### O-MAP-MS data acquisition and analysis

Pooled, labeled peptides were resuspended in (5% acetonitrile, 2% formic acid) and eluted over an in-house pulled 25 cm C18 column (Accucore, Thermo Fisher Scientific) throughout a 180 minute gradient from (6% acetonitrile, 0.125% formic acid) to (32% acetonitrile, 0.125% formic acid). Peptides were analyzed using an SPS-MS3 method on a Thermo Fisher Orbitrap Eclipse to quantify TMTpro reporter ions. Briefly, the duty cycle consisted of three FAIMS (FAIMSpro, Thermo Fisher Scientific) mobility regions at Compensation Voltages (CV)=-40/-60/-80V. At each CV the following were collected within a duty cycle: an MS1 scan (R=120,000, MaxIT=50ms), six MS2 scans (Ion trap, Turbo scan speed, MaxIT=50ms, AGC=200%, CID NCE = 35%), and six SPS-MS3 scans (R=50,000, MaxIT=86ms, HCD NCE = 45%, AGC = 400%). A single dynamic exclusion of 90s was used across all CVs.

Resulting spectra were analyzed using the Comet search algorithm^100^, searched against a full human protein database with forward and reverse protein sequences (Uniprot 10/2020). Precursor monoisotopic peaks were estimated using the Monocle package^101^. Peptides and proteins were filtered to a 1% false discovery rate using the rules of parsimony and protein picking^102^. Protein quantification was done using signal-to-noise estimates of reporter ions. After filtering for contaminants, we performed a two-sided t-test comparing each O-MAP condition using Benjamini-Hochberg adjusted *p* values (i.e. *q*-values). Log 2 fold change of the mean of the biological replicates were also calculated for each biological condition. Nucleolar and nucleoplasmic were extracted from the Human Protein Atlas (accessed 12/13/2022) as defined by the subcellular proteomes lacking multi-compartment overlap. Our nucleolar list is merged from the HPA “nucleoli,” “nucleoli fibrillar center,” and “nucleolar rim” categories. Bi-localized proteins were also obtained using the HPA network plot of the nucleolar and nucleolar proteomes. The consensus speckle proteome was merged between the HPA compartment-unique “nuclear speckles” proteome, and the top 100 hits identified by speckle-targeted TSA-MS^40^. All marker gene sets are provided in (**Supplementary Table 2**). These lists were used to define True Positive (TP) and False Positive (FP) gene sets for Receiver Operating Characteristic (ROC) analysis, as detailed in the text. ROC analysis was performed in Microsoft Excel, as described previously^23, 46^—first by examining the relative recovery rates of TP’s and FP’s as a function of Log_2_(Fold Change), calculated on 0.25-fold increments, and calculating the area under the resulting curve (AUC, *e*.*g*., **Supplementary Fig. 8a**, *left*). The optimal enrichment threshold was defined as the Log_2_(Fold Change) at which the maximum (TP rate) – (FP rate) is obtained (*e*.*g*., **Supplementary Fig. 8a**, *right*). Gene Ontology analysis was performed using MetaScape^52^, and GSEA^53^.

For the labeling time course, k medoid analysis was performed on the mean of three biological replicates of each condition, using Partitioning Around Medoids (PAM) algorithm^103^. The optimal total cluster number was selected based on a maximum of the gap statistic optimization. Nucleolar clusters were chosen by the conspicuous enrichment of the Human Protein Atlas-defined nucleolar proteins (annotated as “enhanced,” “supported,” or “approved” by immunofluorescence data) in four k-medoid clusters. Analysis was based on Fisher’s exact test, with Benjamini-Hochberg adjusted *p* values of each cluster (*q*-values,) less than 0.05. Merging these clusters yielded the speculative nucleolar proteome shown in (**Fig. 2i**,**j**), and listed in (**Supplementary Table 3**). “Annotated nucleolar” proteins were so defined by the HPA, by nucleolar Gene ontology annotation (*i*.*e*., GO:0001650 and GO:0005730), or by Uniprot subcellular nucleolar annotation (from 12/13/2022). Manual curation was performed by literature review.

### O-MAP–Seq

O-MAP-labeled cells (approximately 9×10^6^ cells—one six-well dish—per replicate; three biological replicates per experimental condition) were harvested by scraping into PBSTq (1x PBST, supplemented with 10 µM Sodium Azide, 10 µM sodium ascorbate) and pelleted by centrifugation at 3,000 x *g* for 10 minutes. Buffer was aspirated and cells were flash frozen in liquid nitrogen and stored at –80°C until use. Pellets were thawed on ice and resuspended by gentle pipetting in 1000 µL ice cold RIPA Buffer (50 mM Tris-HCl pH 7.5, 150 mM NaCl, 0.1% (w/v) SDS, 0.5% (w/v) Sodium Deoxycholate, 1% (v/v) Triton X-100, 5 mM EDTA, 0.5 mM DTT), supplemented with 1x EDTA-Free Halt Protease Inhibitor Cocktail, 0.1 U/µL RNase-OUT (Thermo Fisher; 10777019)), and 10mM Sodium Azide, and rocked end-over-end at 4°C for five minutes. Samples were then sheared using a Branson Digital Sonifier 250 outfitted with a double stepped microtip, at 10–12 Watts for 30 s intervals (0.7 s on; 1.3 s off), with 30 s resting steps between intervals, seven intervals total. Samples were held in ice-cold metal thermal blocks throughout sonication. Lysates were then clarified by centrifugation at 15,000 x *g* for 10 min at 4°C, moved to fresh tubes and diluted with 1 mL Native Lysis Buffer (NLB: 25 mM Tris-HCl pH 7.5, 150 mM KCl, 0.5% (v/v) NP-40, 5 mM EDTA, 0.5 mM DTT), supplemented with 1X Halt Protease Inhibitor Cocktail, 0.1 U/µL RNase-OUT and 10 mM sodium azide. For each sample, 5% was removed as “input,” and to the remainder was added 100 µL of Pierce streptavidin magnetic bead slurry that had been equilibrated by two washes in 1:1 RIPA:NLB supplemented with 10mM sodium azide, 0.1 U/µL RNase-OUT, and 1X Halt Protease Inhibitor Cocktail. Samples were incubated for 2 hr at room temperature with end-over-end agitation. Beads were then washed with the following series of buffers (1 mL each, 5 min per wash at room-temperature with end-over-end agitation). All buffers were supplemented with 1x EDTA-Free Halt protease inhibitor cocktail, 0.1 U/µL RNase-OUT, and 0.5 mM DTT, unless otherwise noted: (1) RIPA, supplemented with 10 mM Sodium Azide; (2) RIPA alone (3) High Salt Buffer (1 M KCl, 50 mM Tris-HCl pH 7.5, 5 mM EDTA) (4) Urea Buffer (2 M Urea, 50mM Tris-HCl pH 7.5, 5 mM EDTA) (5) 1:1 RIPA:NLB, without protease inhibitors (6) NLB, without protease inhibitors, (7– 8) two washes in TE buffer (10 mM Tris-HCl pH 7.5, 1 mM EDTA), without protease inhibitors.

RNA was isolated from both input and O-MAP-enriched samples by proteolysis in 100 µL Elution Buffer (2% (v/v) N-lauryl sarcoside, 10 mM EDTA, 5 mM DTT, 200 μg proteinase K (Thermo Fisher; AM2548), in 1x PBS). Reactions were shaken at 700 rpm in a Mixer HC (USA Scientific) for 1 hour at 42°C, followed by 1 hour at 60°C. RNA was then extracted once with 1 volume of phenol pH 4.3, and twice thereafter with an equal volume of absolute chloroform. Reactions were supplemented with 15 µg Glycoblue (Thermo Fisher; AM9515) and NaCl to 300 mM, and ethanol precipitated at –20°C overnight. RNA was harvested by centrifugation at 15,000 x *g* for 30 minutes at 4°C, washed twice with 70% ethanol, and resuspended in 84.75 µL nuclease free water.

Contaminating DNA was removed by digestion with 5 U RQ1 RNase-free DNase I (Promega; M6101) in 100 µL of the manufacturer’s supplied buffer (1x final concentration) at 37°C for 30 min, and this reaction was terminated by addition of EDTA to 15 mM, final. RNA was purified by phenol extraction and ethanol precipitation, as described above, and resuspended in 15 µL nuclease free water. Sample concentration was measured using a Nanodrop One (Thermo Fisher).

### O-MAP-Seq library preparation and sequencing

Ribosomal RNA was first depleted by RNase-H digestion, using pools of antisense DNA oligonucleotides that targeted mature rRNAs, but not the pre-rRNA “transcribed spacer” domains^62^ as described previously^62^. For Patski cells (**Fig. 3h–j**), rRNA was removed using a NEBNext rRNA Depletion Kit (NEB; E6310), according to the manufacturer’s protocol, and RNA was further purified using acidic phenol extraction, chloroform cleanup, and ethanol precipitation. The antisense oligos used for HeLa cells (**Fig. 3a–g**), described previously^62^, were synthesized as an oPool (IDT; 50 pmol per oligo, unmodified), desalted using a Zymo OligoClean and Concentrator Kit, following the manufacturer’s instructions, and resuspended into nuclease-free water. 1 µg of antisense probes were added to 1 µg of RNA (whole cell, or O-MAP-enriched), in 200 mM NaCl, 100 mM Tris-HCl, pH 7.4, at a final volume 10 µL. This solution was heated to 95°C for 2 minutes and then slowly cooled to 45°C at a rate of –0.1°C/s, using a ProFlex PCR system (Thermo Fisher). Reactions were then supplemented with 10 µL of preheated RNase H mix (10 U Hybridase Thermostable RNase H (Lucigen; H39500), 20 mM MgCl_2_). Reactions were incubated at 45°C for 30 minutes and placed on ice. RNA was then purified by acidic phenol-chloroform extraction and ethanol precipitation, residual DNA was removed using RQ1 DNase, and RNA was again purified by phenol-chloroform extraction and ethanol precipitation, as described above.

Each sample was quantified on a Nanodrop One (Thermo Fisher). Sequencing libraries were prepared from 300 ng RNA using the NEBNext Ultra II Directional RNA Library Prep Kit and NEBNext Multiplex Oligos for Illumina (NEB; E7760 and E7735), according to the manufacturer’s instructions. Three biological replicates were used for each experimental condition; each library was given a unique index. Libraries were quantified using the NEBNext Library Quant Kit for Illumina (NEB; E7360), following the manufacturer’s instructions, and the quality of these libraries was confirmed using an Agilent 4200 TapeStation with an “DNA High Sensitivity” kit (Fred Huch Genomics Core). Libraries were pooled in equimolar concentrations to 20 nM aggregate concentration in nuclease-free water, with no more than 12 libraries per pool. These were subjected to 150 cycles of paired-end sequencing, followed by indexing, on one lane of an Illumina HiSeq 4000 per pool, run in high output mode (Azenta Life Sciences).

### O-MAP-Seq data analysis

For gene- and isoform-specific expression analyses (**Figs. 3b-d and 3h**, *left*), raw RNA-seq FASTA files were aligned to reference genomes using HISAT2 version 2.2.1, in the paired-end setting with default parameters^104^ For 47S-O-MAP, reads were mapped to a modified GRCh38 genome assembly (courtesy of T. Moss, Université Laval) in which all rDNA repeats are replaced by a single copy of the consensus rDNA locus as an extra chromosome (“GRCh38_rDNA”)^105^. For *XIST-*O-MAP, reads were mapped to mm10. The resulting SAM files were converted to BAM format and sorted using Samtools^106^ version 1.15.1. Bigwig files for visualizing strand-specific information were created using deepTools^107^ version 3.5.1 with parameters: --filterRNAstrand forward/reverse --binSize 1 --normalizeusingBPM. Mapped reads were quantified using StringTie^108^ version 2.2.1 and the StringTie output was prepared for differential expression analysis using the prepDE.py function. The resulting gene and transcript count matrices were used for differential expression analysis using DESeq2 (Ref. ^109^) with a FDR cutoff of 0.05.

Transposable element (TE) expression analysis (**Figs. 3g** and **h**, *right*) was performed using the TEtranscripts pipeline^110^. Briefly, raw RNA-seq fasta files were mapped to GRCh38 (for 47S) or mm10 (for *XIST*), using STAR^111^ version 2.7.10a, allowing multi-mapped reads with the following settings: --sjdbOverhang 149 -- winAnchorMultimapNmax 200 --outFilterMultimapNmax 100 --outSAMtype BAM Unsorted. TE expression was then quantified using TEtranscripts^110^ version 2.2.1, with the following parameters: --stranded reverse –mode multi --minread 1 --padj 0.05 -i 100. Differentially expressed TEs were identified using DESeq2 version 1.34.0 with a FDR cutoff of 0.05.

Volcano plots were generated using EnhancedVolcano version 1.12.0. All statistical analysis (Fisher’s exact test, hypergeometric distribution test, or Student’s t-test, where appropriate) was performed in R or in python using the ggplot2 (https:// https://ggplot2.tidyverse.org/), or seaborn^112^ and matplotlib^113^ modules.

### O-MAP-ChIP

O-MAP-labeled cells (approximately 5×10^6^ per replicate; two biological replicates per experimental condition) were harvested in PBSTq (1x PBST, supplemented with 10 µM Sodium Azide, 10 µM sodium ascorbate) by scraping, and pelleted by centrifugation at 3,000 x *g* for 10 minutes. Buffer was aspirated and cells were frozen in liquid nitrogen and stored at –80°C until use. Pellets were thawed on ice and gently resuspended by pipetting in 1 mL CLB (20 mM Tris pH 8.0, 85 mM KCl, 0.5% (v/v) NP-40), supplemented with 1x Halt EDTA-Free protease inhibitor cocktail and 10 mM Sodium Azide, for 10 minutes. Lysates were then clarified by centrifugation at 3,000 x *g* for five minutes at 4°C. Supernatant was aspirated, and samples were subjected to another round of CLB extraction, clarification, and supernatant aspiration. Pellets were then lysed by gentle pipetting in 1 mL of NLB (10 mM Tris-HCl pH 7.5, 1% (v/v) NP-40, 0.5% (w/v) Sodium Deoxycholate, 0.1% (w/v) SDS) and incubated on ice for 10 minutes. Samples were then sheared using a Branson Digital Sonifier outfitted with a double stepped microtip, at 10–12 Watts over 30 s intervals (0.7 s on; 1.3 s off), with 30 s resting steps between intervals, 18 intervals total. This resulted in an average shearing size of approximately 200 bp, as gauged on an agarose gel. Samples were held in ice-cold metal thermal blocks throughout sonication. Lysates were then clarified by centrifugation at 15,000 x *g* for 10 minutes at 4°C and supernatants were moved to fresh tubes. For each sample, 10% was removed as ‘input’; to the remainder was added 100 µL of streptavidin-coated magnetic bead slurry that had been equilibrated by two washes in NLB. Samples were incubated for 2 hours at room-temperature with end-over-end agitation. Beads were subsequently washed with the following series of buffers (1 mL each, 5 minutes per wash, at room-temperature, with gentle end-over-end agitation): (1) NLB, supplemented with 5 mM EDTA, 10 mM Sodium Azide and protease inhibitors (1x Halt EDTA-free Protease Inhibitor Cocktail), 150 mM NaCl; (2) NLB, supplemented with 5 mM EDTA, 10mM Sodium Azide and protease inhibitors, (3–4) two washes in 1 M KCl, 10 mM Tris-HCl pH 7.5, 5 mM EDTA, (5–6) two washes in 2 M Urea, 10 mM Tris-HCl pH 7.5, 5 mM EDTA, (7) 10 mM Tris-HCl pH 7.5, 1% (w/v) SDS, (8–9) 10 mM Tris-HCl pH 7.5, 1 mM EDTA.

DNA was isolated from both input and enriched samples by proteolysis in 100uL of Elution Buffer (2% (v/v) N-lauryl Sarcoside, 10 mM EDTA, 5 mM DTT, in 1x PBS, supplemented with 200 μg proteinase K). Samples were shaken for 1 hour at 700 rpm in a Mixer HC at 65°C. Supernatants were transferred to 0.2 mL tubes and incubated at 65°C overnight in a thermocycler. DNA was then extracted with an equal volume of phenol pH 6.6, followed by two extractions in equal volumes of absolute chloroform. Samples were supplemented with 1 µg GlycoBlue and NaCl to 300 mM final, and ethanol precipitated at –20°C overnight. DNA was harvested by centrifugation at 15,000 x *g* for 30 minutes at 4°C, washed twice with 70% ethanol, and resuspended into 20 uL nuclease free water. To remove residual RNA, each sample was supplemented with 10 µg RNase Cocktail Enzyme Mix (Thermo Fisher; AM2286) and incubated at 37°C for 1 hour. DNA was then purified by phenol extraction and ethanol precipitation as described above and resuspended in 20 µL nuclease-free water.

### O-MAP-ChIP library preparation and sequencing

DNA samples were quantified using a NanoDrop One. 300 ng DNA of each sample was used for library preparation, using the NEBNext Ultra II DNA Library Prep Kit and NEBNext Multiplex Oligos for Illumina (NEB; E7645 and E7335), according to the manufacturer’s instructions. Two biological replicates were used per experimental condition; each library was given a unique index during synthesis. Library concentrations were measured using the NEBNext Library Quant Kit for Illumina, and the quality of each sample was confirmed using an Agilent 4200 TapeStation with a “DNA High Sensitivity” kit (Fred Hutch Genomics Core). Libraries were pooled in equimolar concentrations to 20 nM aggregate concentration in nuclease-free water, with no more than eight libraries per pool. These were subjected to 150 cycles of paired-end sequencing, followed by indexing, on one lane an Illumina HiSeq 4000 per pool, run in high output mode (Azenta Life Sciences).

### O-MAP-ChIP data analysis

Deep sequencing reads were trimmed using TrimGalore! (https://www.bioinformatics.babraham.ac.uk/projects/trim_galore/) with parameters -q 30 --phred33, and mapped to the appropriate reference genome using Bowtie2 version 2.4.4(Ref. ^114^). For 47S-O-MAP-ChIP, reads were mapped to GRCH38_rDNA^105^; for *XIST*, reads were mapped to mm10. Duplicate reads were removed with the Picard MarkDuplicates function (http://broadinstitute.github.io/picard) before peak calling. O-MAP-ChIP data were normalized to replicate-matched input samples. For 47S-O-MAP, nucleolar associated domains were called by merging peaks from Enriched Domain Detector (EDD)^115^ and epic2 (Ref. ^116^). EDD was performed using default settings and— because NADs are enriched for highly repetitive sequences like centromeres—an empty BED file for the blacklist region. Epic2 peaks were called with the settings --bin-size 50000 -g 2. Peaks from EDD and epic2 were first concatenated and then merged with the BEDtools^117^ merge function, using the default settings. For *XIST*-O-MAP, regions of enrichment were called using MACS2 (Ref. ^118^) with the broadPeak setting. Further statistical analysis was perfomed in R or python, as described for O-MAP-Seq, above.

Epigenomic analysis of NADs was performed using ChromHMM version 1.22. The OverlapEnrichment function was called using the E117_25_imputed12marks_hg38lift_segments.bed file from the RoadMap Epigenomics Project and the final NAD calls in BED file format. Raw enrichment of each epigenomic signature for each cell line was plotted as a heatmap using seaborn version 0.10.1.

SNP analysis of the allelic segregation of *XIST-*interacting chromatin regions (**Fig. 4b**) was performed as previously described, using a method optimized for the Patski cell line^77^. Briefly, analysis was restricted to MACS2-enriched regions along chromosome X with a coverage of at least five reads. Each region’s allelic proportion was then calculated by computing (X_i_/(X_a_+X_i_)) for each read within the region. Regions with an allelic proportion >0.7 were designated as X_i_-specific; those with an allelic proportion <0.3 were designated as X_a_-specific; those between 0.3 and 0.7 were classified as common to both alleles.

### Streptavidin blotting

O-MAP labeled samples were lysed, sonicated, and clarified as described for O-MAP-MS. Samples were quantified by BCA, supplemented with 2x laemmli loading buffer and heated to 95°C for 10 minutes. Samples, standardized by protein mass, were loaded and separated on 10% SDS-PAGE gels, transferred onto PVDF membranes and stained with Ponceau S (Sigma P7170). Membranes were blocked with 5% (w/v) powdered milk in TBS-T (20 mM Tris, pH 7.5, 150 mM NaCl, 0.1% (v/v) Tween-20) for one hour with rocking at room temperature. Membranes were washed three times in TBS-T, 5 minutes per wash, and blotted with streptavidin– HRP conjugate (Cell Signaling Technology, 3999S, diluted 1:4,000 in TBS-T, supplemented with 5% (w/v) BSA) vernight at 4°C. Blots were washed three times for five minutes in TBS-T, developed using the SuperSignal West Pico PLUS Chemiluminescent Kit (Thermo Fisher 34580), and imaged.

### RNA-FISH, O-MAP Probe-Validation assays, and immunofluorescence

For RNA-FISH, cells were fixed in formaldehyde, permeabilized with Triton-X 100, equilibrated in formamide wash buffer, and hybridized to primary probes as described above (*see: O-MAP Day 1*), but omitting the peroxidase inactivation step. Thereafter, samples were washed three times in 30% Formamide Wash Buffer (five minutes per wash; room temperature) and incubated with 50 µL FISH Secondary Probe Mix (100 nM SABER2– AF647 oligo (**Supplementary Table 1**), in 30% Formamide Hybridization Buffer), and covered with a clean coverslip in a hybridization chamber. Samples were incubated for 1 hour at 37°C in the dark. Samples were washed three times with PBST (five minutes per wash), counterstained with DAPI solution (5 µg/mL, in 1x PBST) and either imaged immediately, or stored sealed in vectashield at 4°C.

For combined O-MAP and RNA-FISH experiments (**Figs. 1d–e, 3i**–**j**, and **6**), including the Probe-Validation Assay (**Fig. 1d**), RNA-FISH signal was dramatically diminished when HRP- and Fluorophore-conjugated secondary oligos were hybridized simultaneously, presumably due to fluorophore damage during the *in situ* biotinylation reaction. To avoid this, O-MAP and RNA-FISH were performed sequentially. O-MAP was performed first, as described above (*see: O-MAP Core Protocol*), including the endogenous biotin blocking step (*see: O-MAP Imaging*). After quenching the O-MAP biotinylation reaction, samples were washed three times in 30% Formamide Wash Buffer (5 minutes per wash; room temperature), and incubated with 50 µL FISH Secondary Probe Mix (100 nM SABER2–AF647, in 30% Formamide Hybridization Buffer) for 1 hour at 37°C. Samples were washed four times with PBST (five minutes per wash), stained with neutravidin–fluorophore conjugate, counterstained with DAPI solution for two minutes, and either imaged immediately in 1x PBST or mounted in vectashield.

When combined with immunofluorescence (**Figs. 1b, 3f**, and **5a–b**), O-MAP or RNA-FISH were performed prior to immunostaining. Cells were subjected to the RNA-FISH or O-MAP pipelines, as described above. Following secondary probe hybridization (RNA-FISH) or *in situ* biotinylation and quenching (O-MAP), cells were washed three times in 1x PBST, and then blocked with 5% (w/v) nuclease-free BSA (VWR 0332) in 1x PBST for one hour at room temperature. Samples were then incubated with rabbit anti-*NPM1* (Thermo Fisher; PA517742, used at 1:100 dilution) or rabbit anti-*FBL* (Abcam; ab166630; used at 1:100 dilution), in 1% (w/v) BSA, 1x PBST, for one hour at room temperature with gentle rocking. Samples were washed four times with 1x PBST and then incubated with either AlexaFluor 488-conjugated anti-rabbit (Thermo Fisher; A32731TR, 1:1000 dilution), with neutravidin-DyLight 550 conjugate (Thermo Fisher; 84606, 1:1000 dilution) as appropriate, in 1% (w/v) nuclease-free BSA, 1x PBST, for one hour at room temperature. Samples were washed four times with 1X PBST (15 minutes per wash), counterstained with DAPI for two minutes, and either imaged immediately in 1x PBST or mounted in vectashield.

Fluorescence widefield microscopy was performed on a Leica DM IL, equipped with a HC Fluotar 100x oil immersion objective with a 1.32 numerical aperture and planar correction (Leica; 11506527), a white LED light source (Leica; EL6000) and a DFC365 FX digital camera (Leica; 11547004). The following filter cubes were used: Texas Red (Leica TX2 ET; 11504180; used with Dylight-550 conjugates), Cy5 (Leica Y5 ET; 11504181, used for Alexafluor-647), GFP (Leica GFP ET; 11504174, used for Alexa Fluor-488), and DAPI (Leica DAPI ET; 11504204). Illumination intensity was adjusted using the light source manual control; acquisition times ranged from 40–2000 ms, as controlled by the Leica LASX software. Fluorescence confocal microscopy was performed on a Leica SP8X microscope (UW Keck Imaging Center), outfitted with a HC CS2 63x oil immersion objective, with 1.40 numerical aperture with both planar and apochromatic correction. The average voxel size was 0.06 × 0.06 × 0.346 µm. Samples were illuminated using a 470–670nm tunable White Light Laser system, with a typical laser power of 0.1% for DAPI, 3% for 550 nm, and 30% for 647 nm. Gain and offset settings were adjusted to avoid pixel saturation. Images were line-averaged twice, with an average pixel dwell time of 1.58 µs. A bit-depth of 8 or 16 was used and zoom factor between 1-3 was used for all images.

### Image processing

Images were processed using Fiji^119^ and ImageJ^120^, and multicolor overlays were made using the screen setting in Adobe Photoshop^35^. Most confocal images are maximum projections of z-stacks; the remainder correspond to single z-slices. Brightness and contrast were adjusted for display purposes using Fiji and ImageJ or Adobe Photoshop. In all cases, contrast adjustment was applied to improve signal visibility, by changing the minimum (black) and maximum (white) values only. Automated despeckling was applied when necessary (*e*.*g*. in RNA-FISH images with weak, diffuse speckling in between cells) to reduce residual background signal. Colocalization analysis (**Fig. 3f**,**i**,**j**) was performed using Fiji^121^.

## Notes

### Competing Interest Statement

A.F.T, E.E.K, B.J.B, and D.M.S. have filed for a patent (provisional application 63/300,125) concerning the use of oligonucleotide-directed proximity-labeling to elucidate and visualize subcellular interactions in situ.

