## Supplementary_Figures for "Oligonucleotide-directed proximity-interactome mapping (O-MAP): A unified method for discovering RNA-interacting proteins, transcripts and genomic loci *in situ*"

**Supplementary Figure 1.** RNA-FISH validation of the 47S pre-rRNA-targeting probe set
**Supplementary Figure 2.** Overview and limitations of preliminary O-MAP designs.
**Supplementary Figure 3.** Reproducibility of O-MAP labeling across primary probe sets.
**Supplementary Figure 4.** More O-MAP controls.
**Supplementary Figure 5.** Enrichment of biotinylated proteins for O-MAP-MS.
**Supplementary Figure 6.** Reproducibility of O-MAP-MS.
**Supplementary Figure 7.** Further validation of the one-shot 47S/7SK O-MAP-MS dataset
**Supplementary Figure 8.** Probing the nucleolar proteome with 47S O-MAP-MS
**Supplementary Figure 9.** Probing the 7SK-proximal proteome with O-MAP-MS
**Supplementary Figure 10.** Representative highly ranked Gene Set Enrichment Analysis (GSEA) results.
**Supplementary Figure 11.** K-medoid clustering
**Supplementary Figure 12.** Coverage of the nucleolar proteome during the 47S O-MAP labeling time course.
**Supplementary Figure 13.** 47S O-MAP-Seq enriches known and novel nucleolar transcripts.
**Supplementary Figure 14.** 47S O-MAP-Seq and HyPro-Seq enrich common transcripts.
**Supplementary Figure 15**. *Xist* O-MAP-Seq enriches nascent transcripts of XCI-escape genes.
**Supplementary Figure 16.** Genome maps of HeLa Nucleolar Associated Domains (NADs)
**Supplementary Figure 17.** 7SK O-MAP in cultured Pancreatic Ductal Adenocarcinoma (PDA) cell lines.

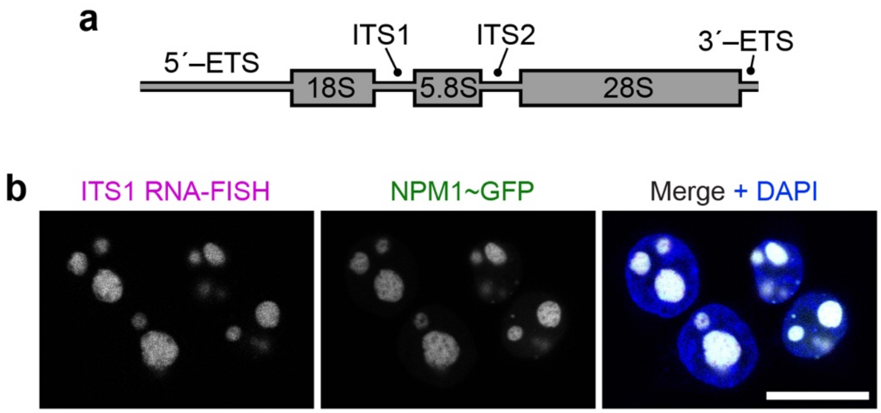


**Supplementary Figure 1. RNA-FISH validation of the 47S pre-rRNA-targeting probe set. a,** Schematic of the 47S pre-rRNA. During ribosome biogenesis, the 18S, 5.8S, and 28S domains are processed and incorporated into mature ribosomes, while the 5´– and 3´–External Transcribed Spacers (5´–ETS and 3´–ETS) and Internal Transcribed Spacers (ITS1 and ITS2) are cleaved from the precursor transcript and degraded within the nucleolus. The 47S pre-rRNA probe set used in the initial stages of O-MAP development target ITS1. **b,** validation of this probe set. HEK293T cells were transiently transfected with GFP-tagged *NPM1*, a nucleolar marker, and subjected to conventional RNA-FISH. Note conspicuous overlap between the *NPM1*~GFP and ITS1 RNA-FISH signals. Scale bar: 20 μm.

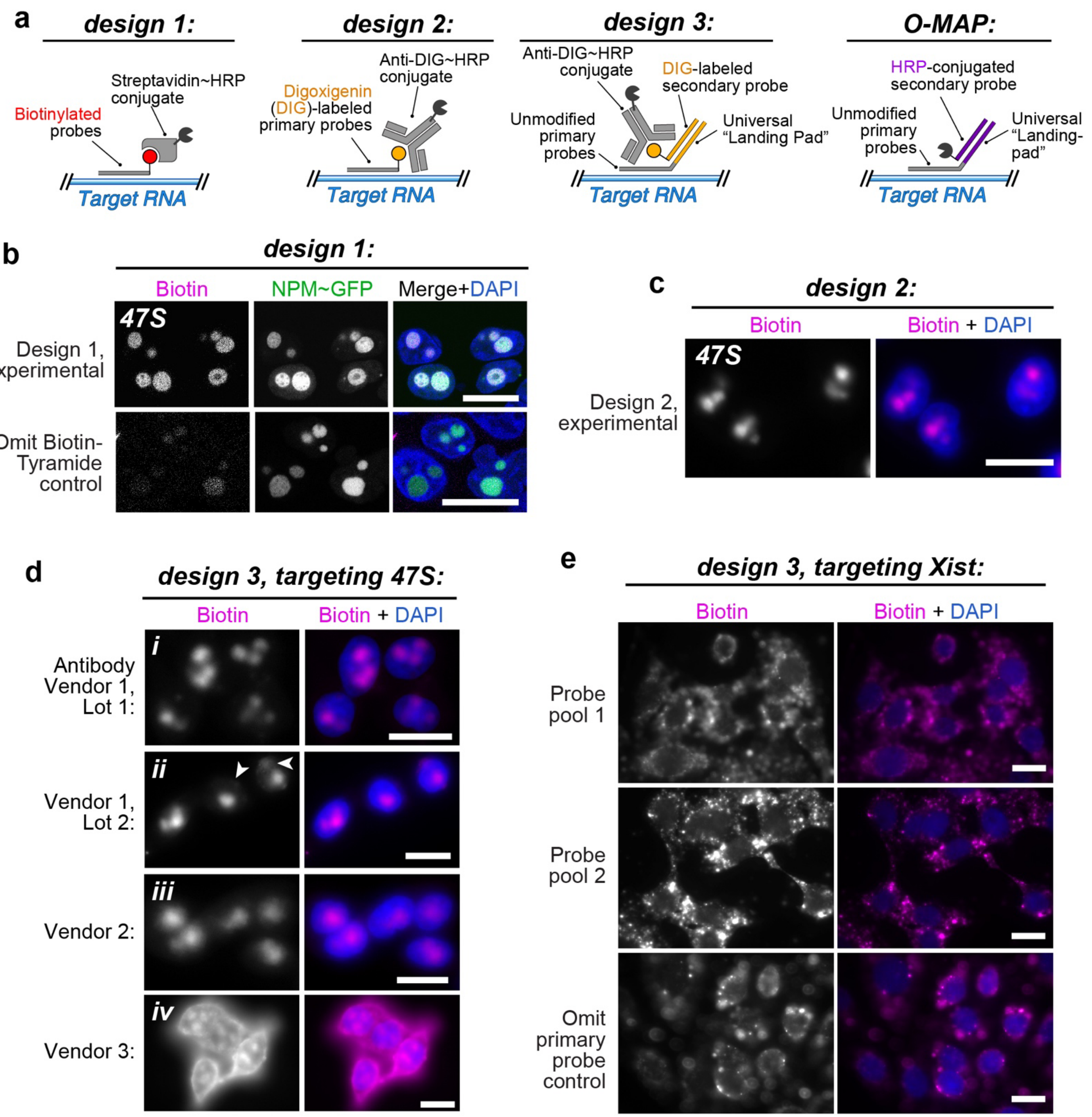


**Supplementary Figure 2. Overview and limitations of preliminary O-MAP designs. a,** summary of RNA-targeted HRP-recruitment strategies tested. Design 1 uses biotinylated primary probes to recruit a streptavidin-HRP conjugate. Designs 2 and 3 use Digoxigenin (DIG)-labeled primary or secondary probes to recruit an HRP–conjugated anti-DIG antibody. Our final O-MAP design, which uses HRP-conjugated oligo probes, is shown for comparison (*see also*, **Fig. 1a**). The same anti-DIG antibody is used in designs 2 and 3; the same "universal landing pad" sequences are used in designs 3 and 4. **b–e**, limitations of Designs 1–3. In all cases biotin was visualized by staining with a fluorescent neutravidin conjugate. **b,** Design 1 was disfavored because *in situ* biotinylation cannot be unambiguously distinguished from biotinylated primary probes. HeLa cells over-expressing NPM1~eGFP were probed using the same 47S-targeting probes as in (**Supplementary Fig. 1**) and the main text, appended on their 3´–termini with biotin. Note nucleolar biotin signal even in the absence of biotin-tyramide (*bottom panels*). We anticipated that this background signal would be especially problematic with low-abundance target RNAs. **c,** Design 2—analogous to HyPro (PMID: 35457249)—was sometimes capable of producing well-resolved nucleolar-targeted biotinylation. HeLa cells are shown. This approach was eventually disfavored due to antibody irreproducibility issues described below, and because the high cost of DIG-labeled oligos would limit its use with low-abundance transcripts, which can require dozens to hundreds of probes. **d,** Design 3 overcomes the oligo cost issue but still suffers from antibody background binding and irreproducibility. HeLa cells were probed with the same 47S-targeting primary probe sets used in the main text (appended with the same "landing pad" modules), a DIG-labeled secondary oligo, and four different lots or vendors of commercial HRP-conjugated antibodies. In some cases, we observed well-resolved RNA-targeted biotinylation (*panel i*), though other lots from the same vendor exhibited off-target labeling (*panel ii, arrows*). Regents from other vendors exhibited varying degrees of spatial blurring (*panel iii*), or conspicuous off-target biotinylation that rivaled or exceeded the target signal (*panel iv*). Compare these results to (**Fig. 1b**). **d,** Design 3 is particularly problematic with lower-abundance RNA targets. Patski cells were probed with the same *Xist*-targeting, landing-pad-extended probes as used in the main text, divided into two sub-pools. Anti-DIG-HRP was from Vendor 1. Note that all conditions—both probe sub-pools and the omit-primary negative control–induced substantial off-target biotinylation. Compare these results to (**Fig. 1e**). All scale bars, 20 µm.

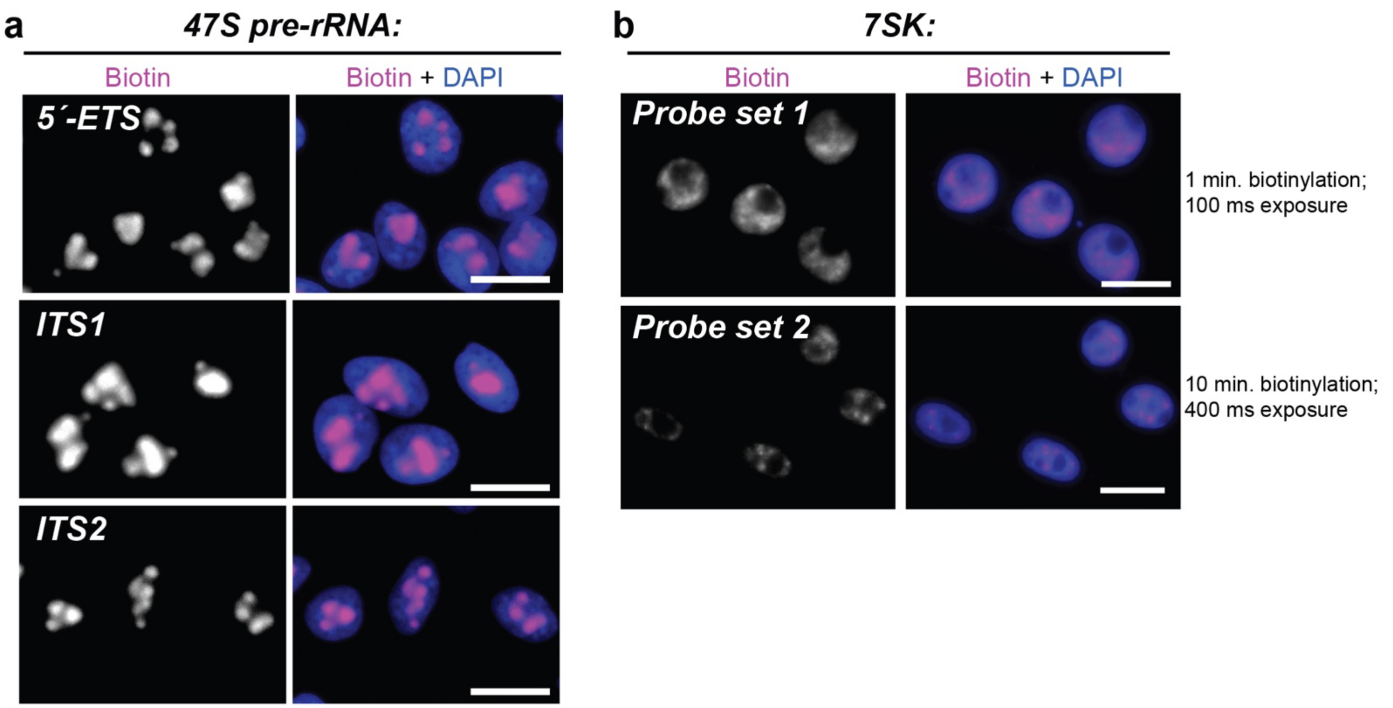


**Supplementary Figure 3. Reproducibility of O-MAP labeling across primary probe sets. a,** Targeting the 47S pre-rRNA. Probe sets targeting the 5´–ETS, ITS1, and ITS2 Transcribed Spacer domains (**Supplementary Fig. 1a**) produced similar patterns of nucleolar *in situ* biotinylation. Probes targeting the 3´–ETS yielded no signal (*data not shown*). **b,** Targeting 7SK. Each set targets the entirety of the 7SK transcript, but were designed to have different hybridization parameters. Note that *in situ* biotinylation and exposure times differed between 7SK probe sets, as indicated (*right*). Biotin was visualized by staining with fluorescent neutravidin, in HeLa cells. Scale bar: 20 μm.

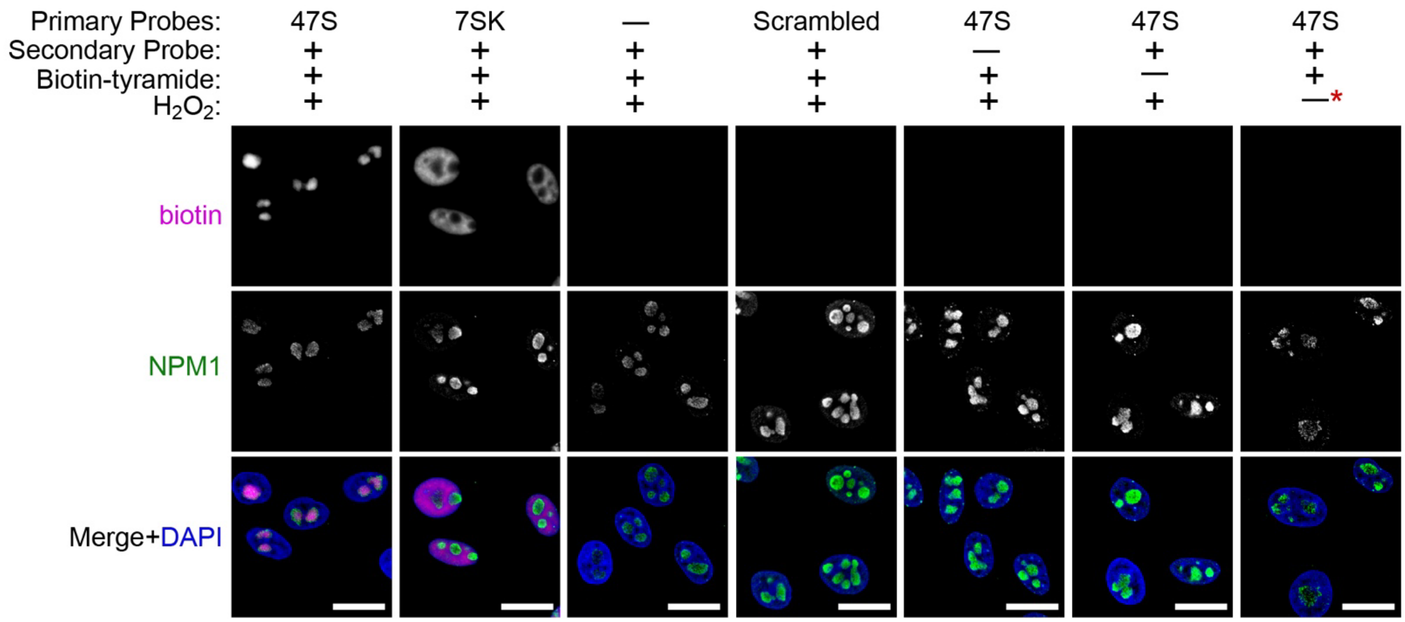


**Supplementary Figure 4. More O-MAP controls.** O-MAP and negative control experiments were performed in HeLa cells, as indicated. Biotin was imaged using a fluorescent neutravidin conjugate; *NPM1* via immunofluorescence. Note that omitting any component of the O-MAP pipeline ablated biotinylation signal. The 47S-O-MAP, 7SK-O-MAP, omit primary and scrambled primary conditions (*left four columns*) are the same images presented in (**Fig. 1b**). In the "omit-$H_2O_2$" condition (*far right, marked* * ), cells were pre-quenched with sodium azide and ascorbic acid prior to the addition of biotin-phenol and $H_2O_2$. Simply removing $H_2O_2$ from the O-MAP protocol still resulted in targeted *in situ* biotinylation (*i.e.* nucleolar labeling using 47S probes), presumably due to photoactivation of HRP. Scale bars, 20 μm.

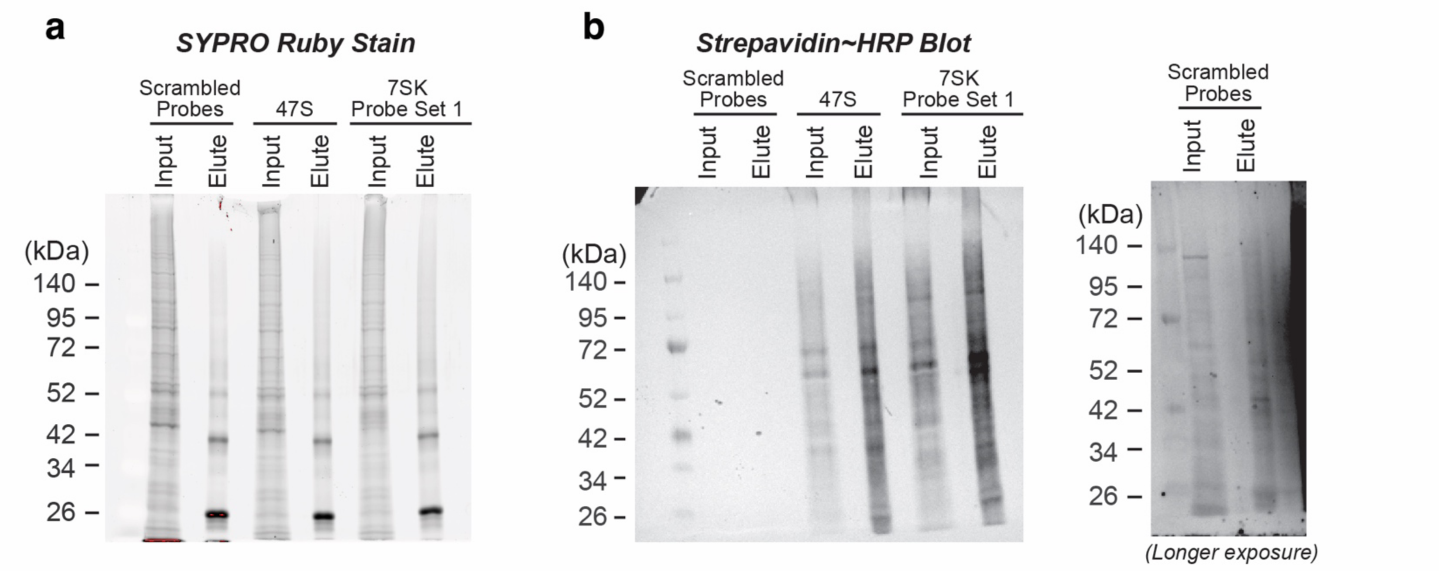


**Supplementary Figure 5. Enrichment of biotinylated proteins for O-MAP-MS.** HeLa cells were O-MAP labeled using 47S-targeting, 7SK-targeting, or scrambled probes, as indicated. For 7SK, probe set 1 was used (**Supplementary Table 1**). Cells were lysed by boiling in SDS (*see methods*), enriched by streptavidin pulldown, and released from the beads by again boiling in SDS. 10 µg samples of the starting lysates ("input") or eluted proteins ("Elute") were separated on 10% SDS PAGE gels. **a,** SYPRO Ruby$^{TM}$-stained gel. The blurriness of bands in the input samples is a consequence of formaldehyde crosslinking. **b,** Streptavidin~HRP blotting of the same gel, to visualize biotinylated proteins. Note the ladder of biotinylated products in both 47S- and 7SK- conditions, and the enrichment of these proteins in the eluted samples. Longer exposure (*right*) reveals a weak background of biotinylated material in the Scrambled probe control.

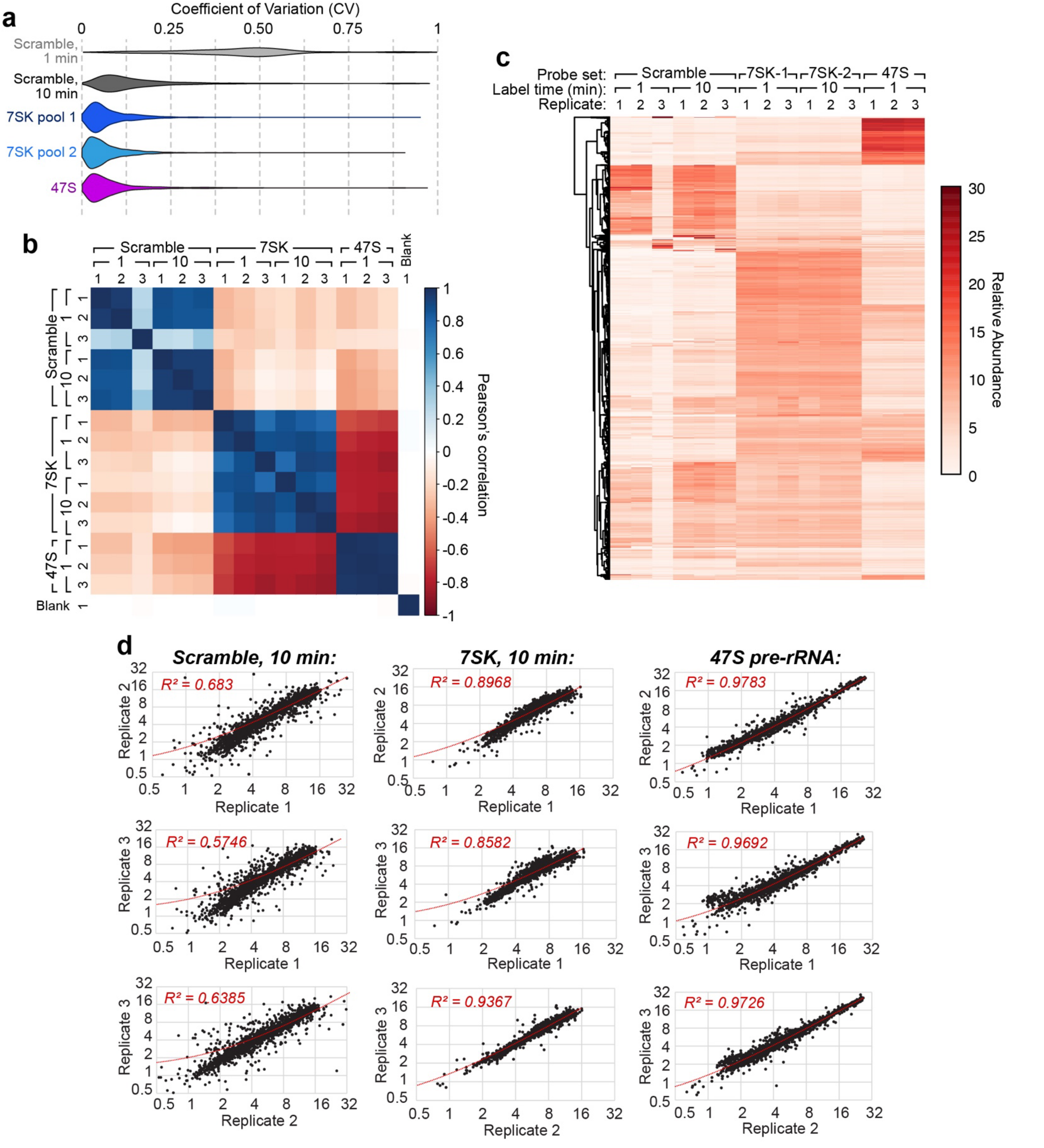


**Supplementary Figure 6. Reproducibility of O-MAP-MS. a,** Average coefficients of Variation (CV) between biological replicates (*n* = 3). Conventionally, CV's below 0.2 are considered acceptable. **b,** Correlation matrix between all samples. Note high positive correlations between biological replicates, highlighting O-MAP's reproducibility, and negative correlations between different sample types (*e.g.* 47S vs 7SK 1 minute), highlighting the unique composition of the proteome captured for each target. **c,** Heatmap of all proteomic data, clustered by unsupervised hierarchical clustering, showing defined clades of 47S-proximal and 7SK-proximal proteomes. Note that the 7SK 1-minute labeling condition used probe set 1; the 10-minute condition used probe set 2. **d,** Scatter plots comparing scaled protein abundances for individual biological replicates for the Scramble negative control (10 minutes' labeling time, *left*), 7SK probe set two (*center*), and 47S (*right*). Data are plotted on log2 scale and fit by linear regression. $R^2$, Pearson correlations.

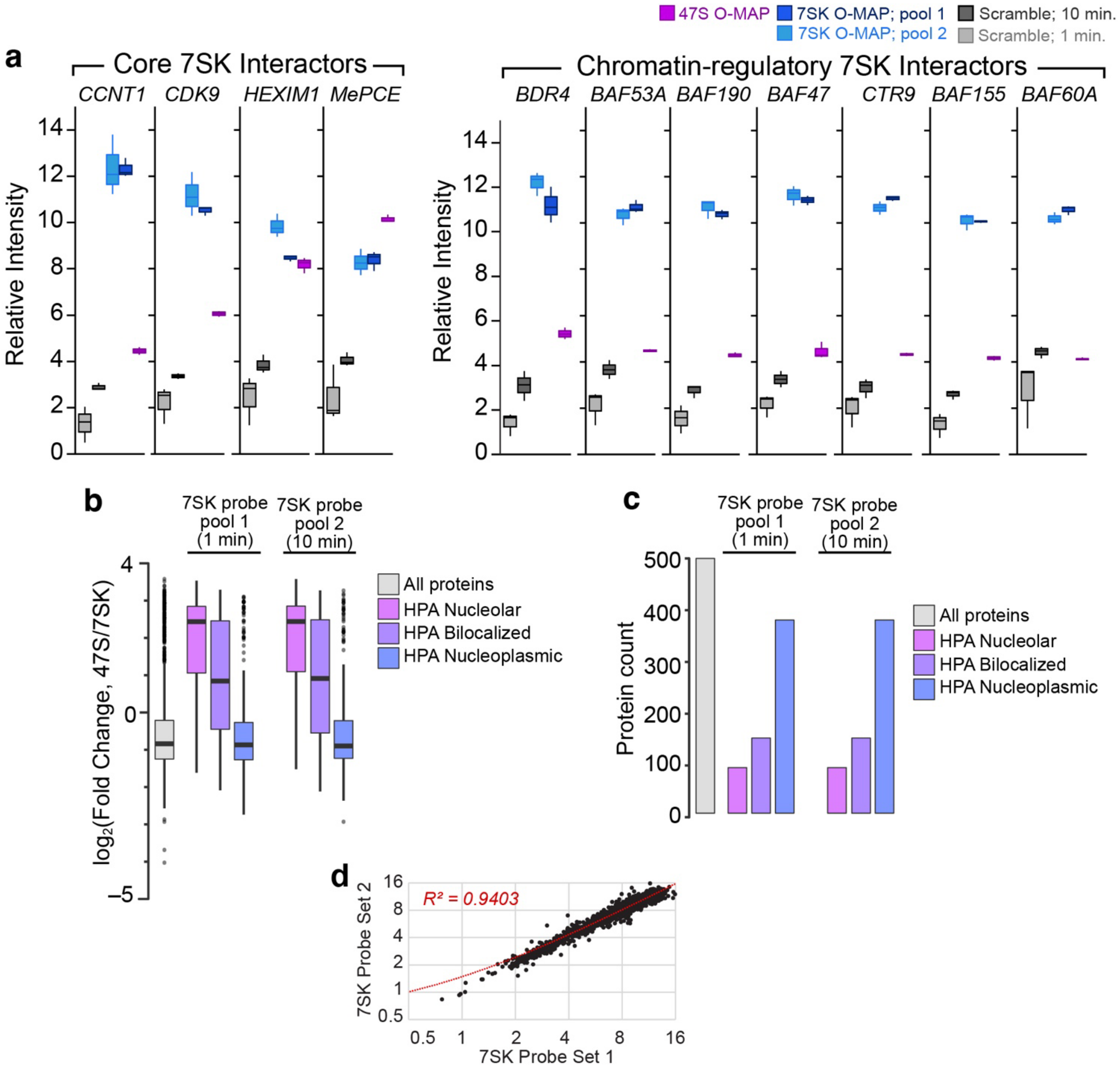


**Supplementary Figure 7. Further validation of the one-shot 47S/7SK O-MAP-MS dataset. a,** Enrichment profiles (as in **Fig. 2b**) of core components of the 7SK particle (*left*) and of recently discovered 7SK-interactors, including members of the BAF complex (*right*). *n = 3* biological replicates. Core interactors *LARP7* and *HEXIM2* were not detectable above noise. **b,** Differential enrichment of the nuclear proteome, defined by the Human Protein Atlas (HPA) as being strictly nucleolar (*magenta*), strictly nucleoplasmic (*blue*) or bilocalized between the two compartments (*purple*). Fold changes calculated using the same 47S-targeting probe set, relative to either 7SK-targeting probe pool 1 (*left*), or 2 (*right*). In both cases, note the strong enrichment of nucleolar proteins, intermediate enrichment of bilocalized proteins, and de-enrichment of nucleoplasmic proteins. Note that the two 7SK probe sets gave nearly identical results. This is further supported by: **c,** the total number of proteins from each class that were observed with both probe sets, and **d,** correlation between average protein abundances observed using each probe set, plotted and fit as in (**Supplementary Fig. 6d**).

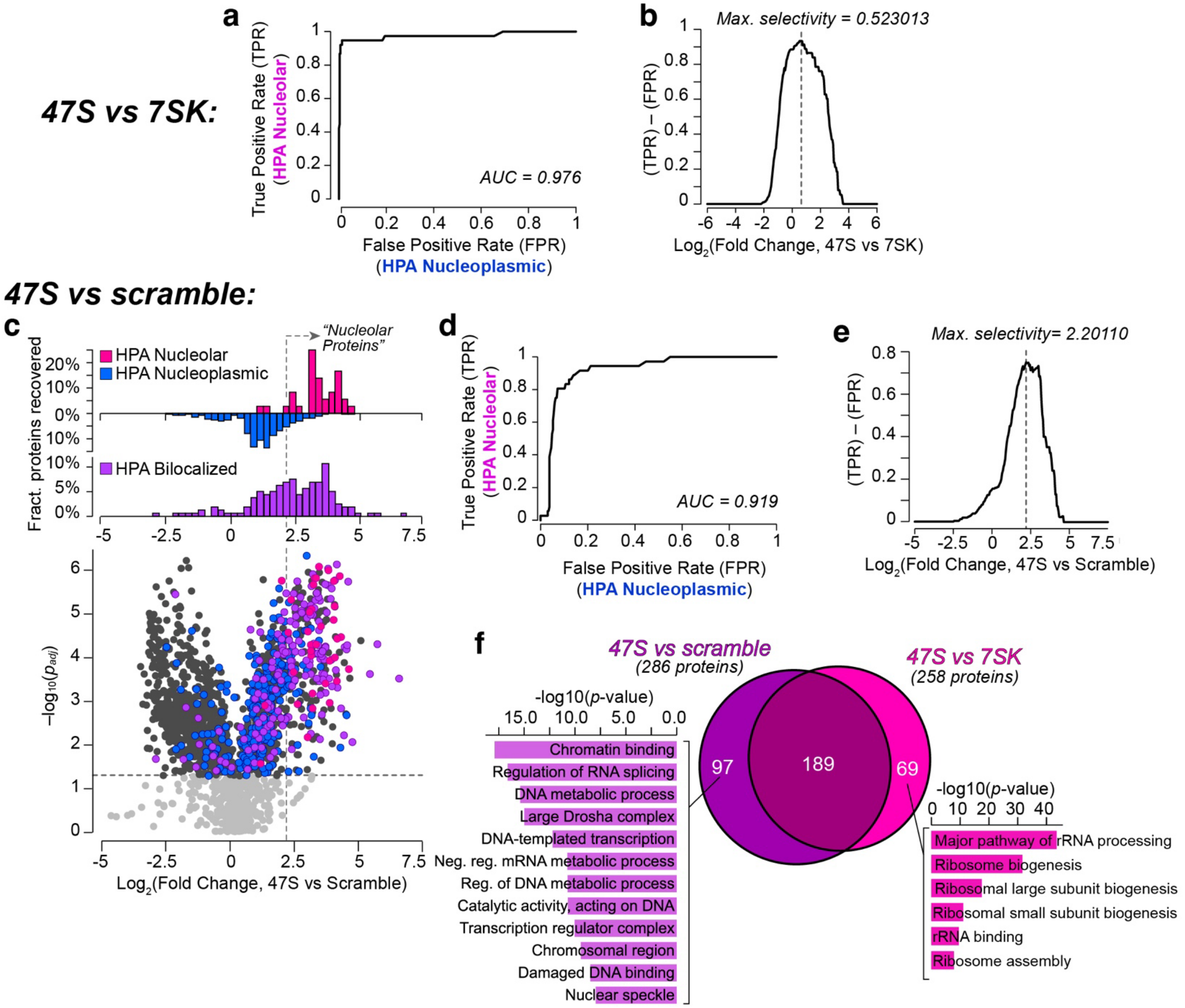


**Supplementary Figure 8. Probing the nucleolar proteome with 47S O-MAP-MS. a,** Receiving-Operator Characteristic (ROC) analysis of the 47S vs 7SK O-MAP-MS experiment. True Positive and False Positive proteins were defined using lists of exclusively nucleolar and exclusively nucleoplasmic proteins, respectively, as reported by the Human Protein Atlas (HPA). An Area Under the Curve (AUC) of nearly 1.0 suggests strong and highly sensitive selectivity for nucleolar proteins over the nucleoplasmic proteome. **b**, These data were used to derive an optimal $Log_2$(fold change, 47S/7SK) cutoff value of 0.523, and to define a putative list of 258 O-MAP core nucleolar proteins, as described in (**Figure 2**). **c–e**, parallel analysis using (47S/Scramble controls), instead of (47S/7SK). **c,** Volcano plot and histograms of showing the enrichment of HPA-nucleolar, HPA-Nucleoplasmic, and HPA-bilocalized proteins, using the same protein marker reference lists as in (**a–b**), and (**Fig. 2c**,**d**). **d,** ROC analysis of the (47S/Scramble) data demonstrates slightly lower sensitivity than that of the (47S/7SK) analysis, though still exceptionally. **e**, As in (**b**), these data were used to determine an optimal $Log_2$(fold change, 47S/Scramble) cutoff value of 2.201, defining a putative cohort of 286 O-MAP core nucleolar proteins. **f,** the putative nucleolar proteomes derived from the (47S/7SK) and (47S/Scramble) ROC analyses show considerable overlap (66–73%). Outliers were used for Gene Ontology (GO)-term analysis. Factors uniquely captured by the (47S/7SK) analysis were highly enriched for ribosome biogenesis factors, while those unique to the (47S/Scramble) analysis were enriched for nucleoplasmic functions. This suggests that the (47S/7SK) comparison more precisely captures the nucleolar proteome.

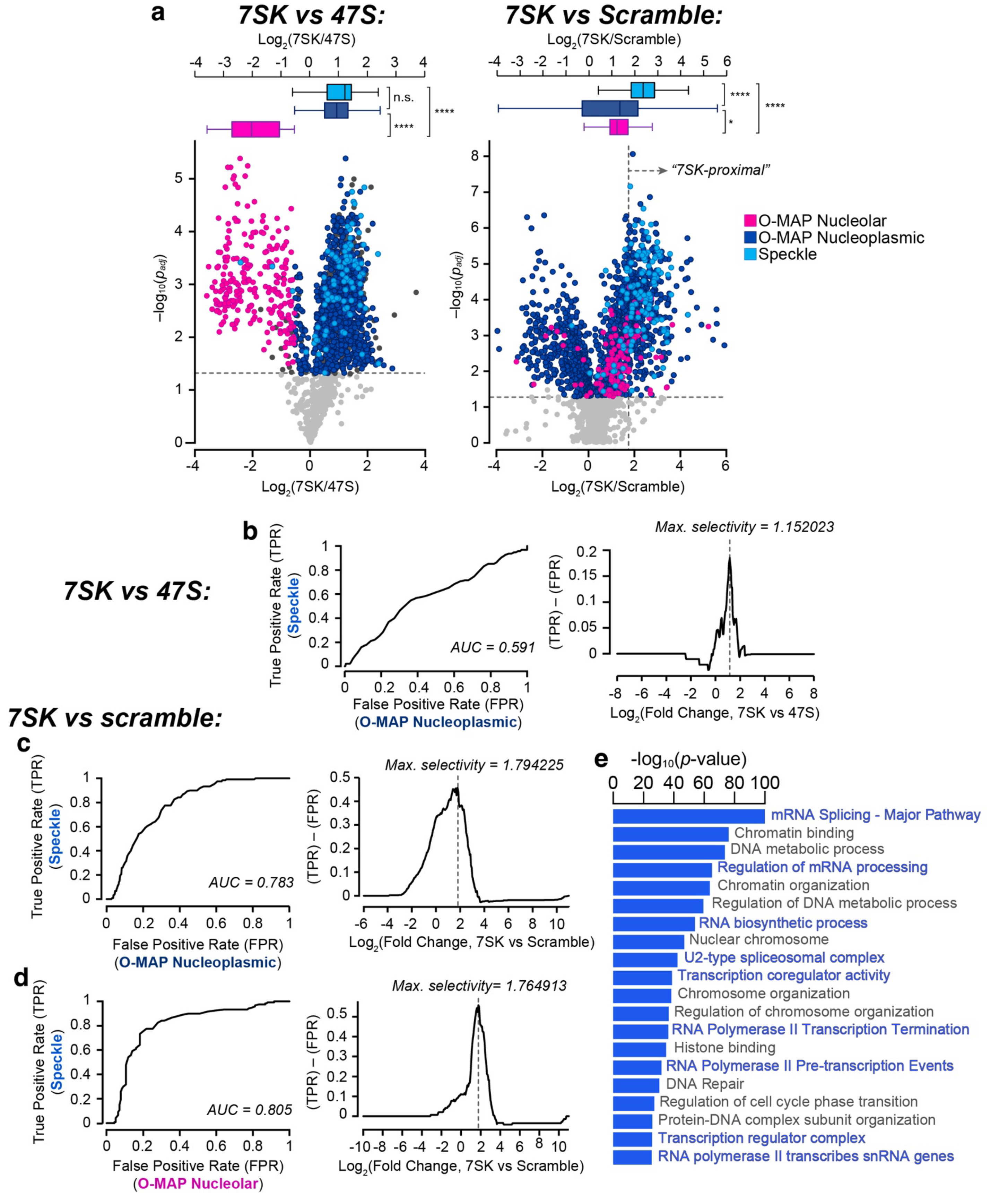


**Supplementary Figure 9. Probing the 7SK-proximal proteome with O-MAP-MS. a,** Identifying the optimal basis of comparison for probing the 7SK-proximal compartment: (7SK/47S) or (7SK/Scramble). For each comparison, volcano plots demonstrate the enrichment of the nucleolar- and nucleoplasmic- proteomes, as derived from O-MAP analysis (**Fig. 2** and **Supplementary Fig. 8**), and of known Nuclear Speckle proteins, markers of the 7SK-proximal compartment. Box-whisker plots (*above*) summarize the distribution of each protein group (significance testing: two-tailed, heteroscedastic Student's t-tests; n.s: not significant, * $p$=0.05; **** $p < 1x10^{-5}$). In each comparison, speckle proteins are significantly enriched relative to the nucleolar proteome, but in the (7SK/47S) comparison (*left*) these proteins are indistinguishable from the broader nucleoplasmic proteome ($p = 0.14$). In contrast, in the (7SK/Scramble) comparison (*right*) Speckle proteins are significantly enriched relative to the nucleoplasmic outgroup ($p = 5 \times 10^{-10}$). This suggests that (7SK/Scramble) is more selective for 7SK-proximal interactors over the broader nucleoplasmic proteome. This is corroborated by Receiver-Operating Characteristic (ROC) analysis, as described below, which was used to define the "7SK-proximal" cutoff in the righthand panel. **b,** ROC analysis of (7SK/47S), using speckle proteins as true positives and O-MAP-nucleolar proteins as false positives. An area under the curve (AUC) of approaching 0.5 suggests a nearly complete absence of signal. **c,** ROC analysis of (7SK/Scramble), using the same True Positive and False Positive lists as in (**b**), suggests a strong and specific separation between the protein populations, and an optimal $Log_2$(fold change) cutoff of ≥ 1.794. **d**, A nearly identical result is obtained when using O-MAP-nucleolar proteins as False Positives. These cutoffs define a cohort of 510 putative 7SK-proximal proteins. **e,** Gene Ontology (GO)-term analysis on this cohort is highly enriched for biological processes involved in pre-mRNA biogenesis, highlighted in blue.

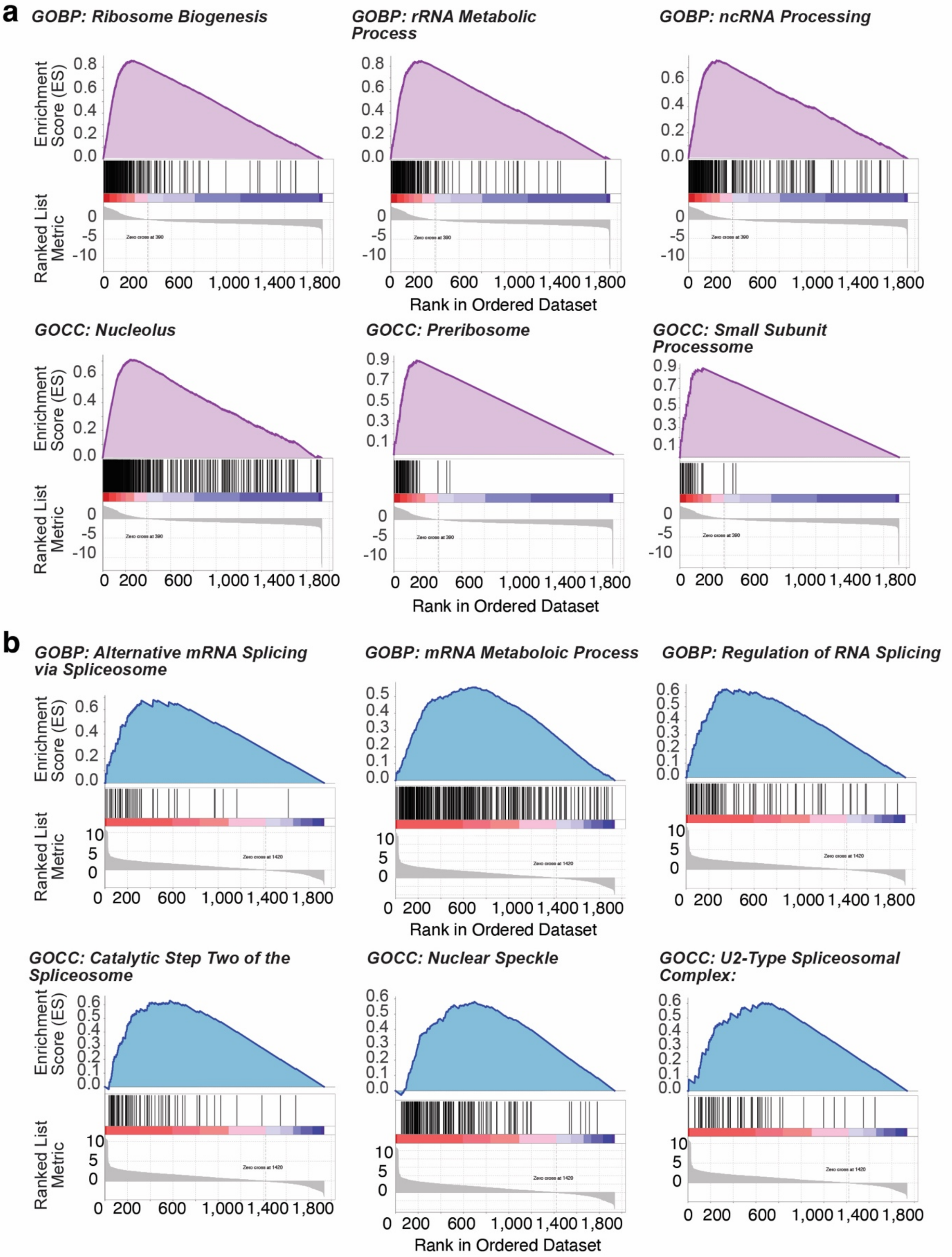


**Supplementary Figure 10. Representative highly ranked Gene Set Enrichment Analysis (GSEA) results. a,** GSEA results comparing 47S and 7SK O-MAP-MS. The same ranked gene list was used for all queries. Top Gene Ontology Biological Process (GOBP) and Cellular Component (GOCC) terms are shown. Rank ordering was determined by $Log_2$(Fold Change, 47S/7SK). For all, left-ranking genes (*red*) were more highly enriched from 47S O-MAP than 7SK O-MAP; those ranked toward the right (*blue*) were more enriched from 7SK O-MAP. **b,** GSEA results comparing 7SK O-MAP--MS to Scrambled controls. Gene rank was determined by $Log_2$(Fold Change, 7SK/Scramble). The same gene list was used for all queries. Leftmost genes were 7SK-enriched; rightmost were Scramble-enriched.

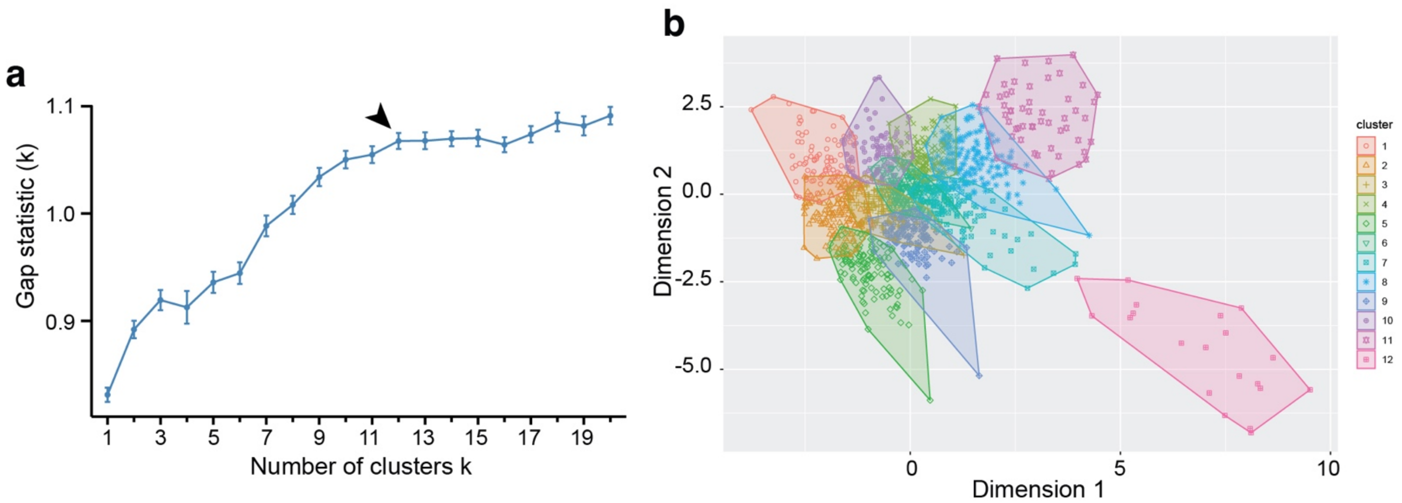


**Supplementary Figure 11. k-medoid clustering. a.**, k-medoid analysis was performed using a Partitioning Around Medoids (PAM) algorithm on the mean of the biological triplicates for each condition. Ultimately, 12 clusters were chosen based on gap statistic optimization (*arrowhead*). **b,** Principle component analysis (PCA) of the twelve k-medoid clusters.

**a**

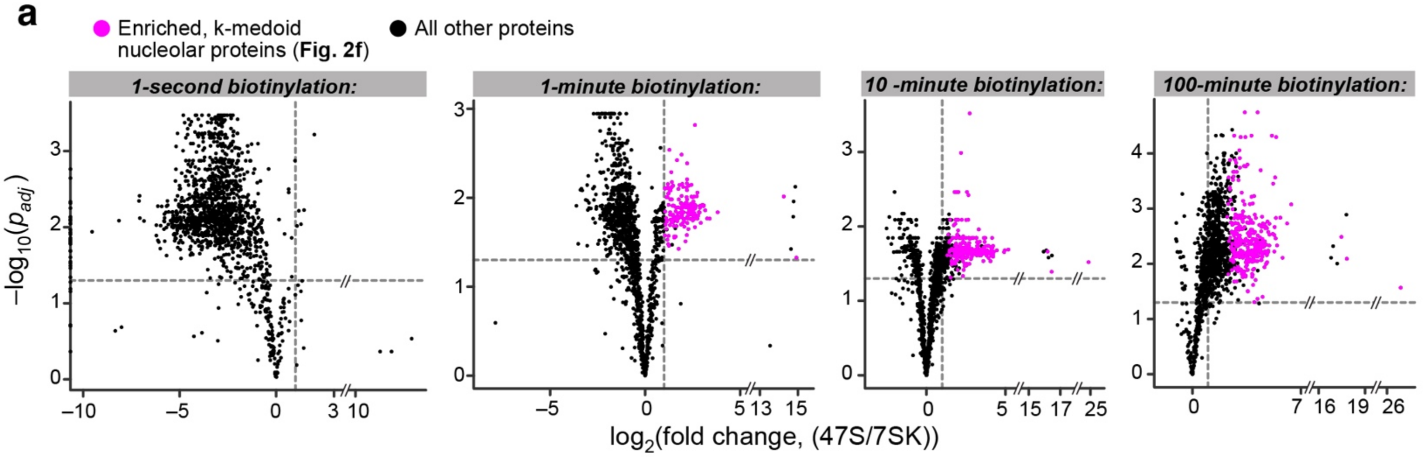


**b**

| | Total enriched proteins | Enriched k-medoid nucleolar proteins (313 total; **Fig. 2f**) |
|---|---|---|
| 1-second biotinylation | 5 | 0 (0%) |
| 1-minute biotinylation | 189 | 178 (56.9%) |
| 10-minute biotinylation | 490 | 313 (100%) |
| 100-minute biotinylation | 994 | 313 (100%) |

**Supplementary Figure 12. Coverage of the nucleolar proteome during the 47S O-MAP labeling time course. a,** Volcano plots for each 47S O-MAP labeling point, calculated relative to the 7SK/10-minute label condition. Enrichment of nucleolar proteins derived from our k-medoid analysis (**Fig. 2e–g; Supplementary Fig. 10**) are highlighted in pink. Significance cutoffs were assigned at $p_{adj} \leq 0.05$ and Fold Change (47S/7SK) ≥ 2.0. (*dotted lines*) **b,** Table summarizing the recovery of the nucleolar proteome at each labeling time point. Note that coverage appears to plateau at 10 minutes.

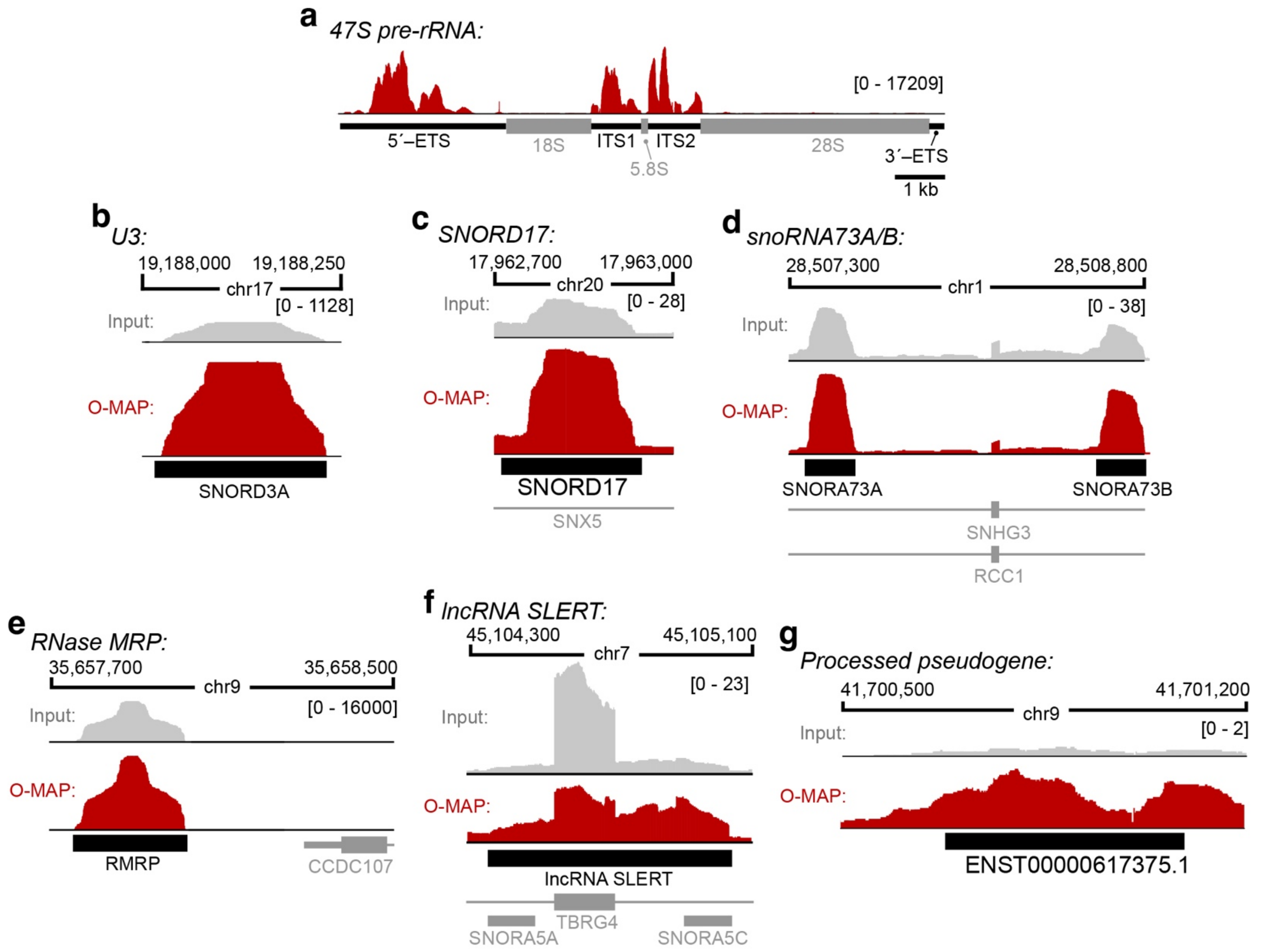


**Supplementary Figure 13. 47S O-MAP-Seq enriches known and novel nucleolar transcripts. a,** the 47S pre-rRNA. Note prominent enrichment for the 5´–ETS, IT1, and ITS2 "transcribed spacer" domains. Sequences corresponding to the mature 18S, 5.8S, and 28S rRNAs are removed during sequencing library preparation. Reads are aligned to a custom genome assembly containing a single copy of the rDNA consensus sequence (courtesy of T. Moss, U. Laval) annotated as a unique chromosome. **b,** the U3 noncoding RNA, which directs key cleavage events during ribosomal biogenesis, **c–d,** exemplar Box C/D (**c**) and H/ACA (**d)** small nucleolar RNAs (snoRNAs). SnoRNAs are often expressed within the introns of protein-coding genes (*gray*). **e,** RNase MRP (*enriched*) upstream of the *CCDC107* gene (*not enriched*). **f,** lncRNA SLERT, which is processed from the two H/ACA snoRNAs embedded in the *TBRG4* gene. **g,** Example of a novel nucleolar transcript—a processed pseudogene—discovered by O-MAP-Seq

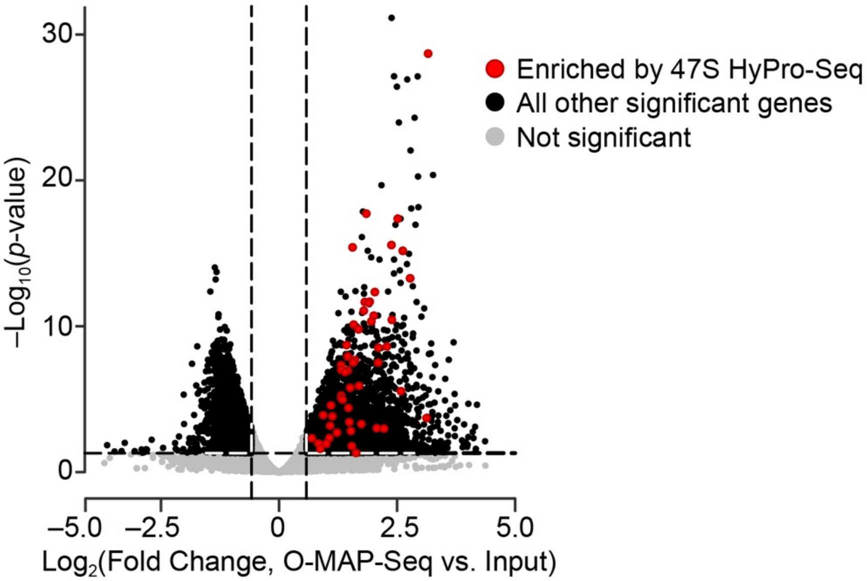


**Supplementary Figure 14**. **47S O-MAP-Seq and HyPro-Seq enrich common transcripts**. Volcano plot of 47S O-MAP-Seq data, merging both coding and noncoding genes (**Fig. 3b**). Transcripts that were reportedly enriched by 47S HyPro-Seq (PMID: 34741808) are indicated (*red*). Note that differences in transcript annotation complicated systematic comparison to HyPro-Seq data. In all, only 53 of the significantly enriched transcripts reported by HyPro-Seq were detected in our data, all of which were also enriched by 47S O-MAP-Seq.

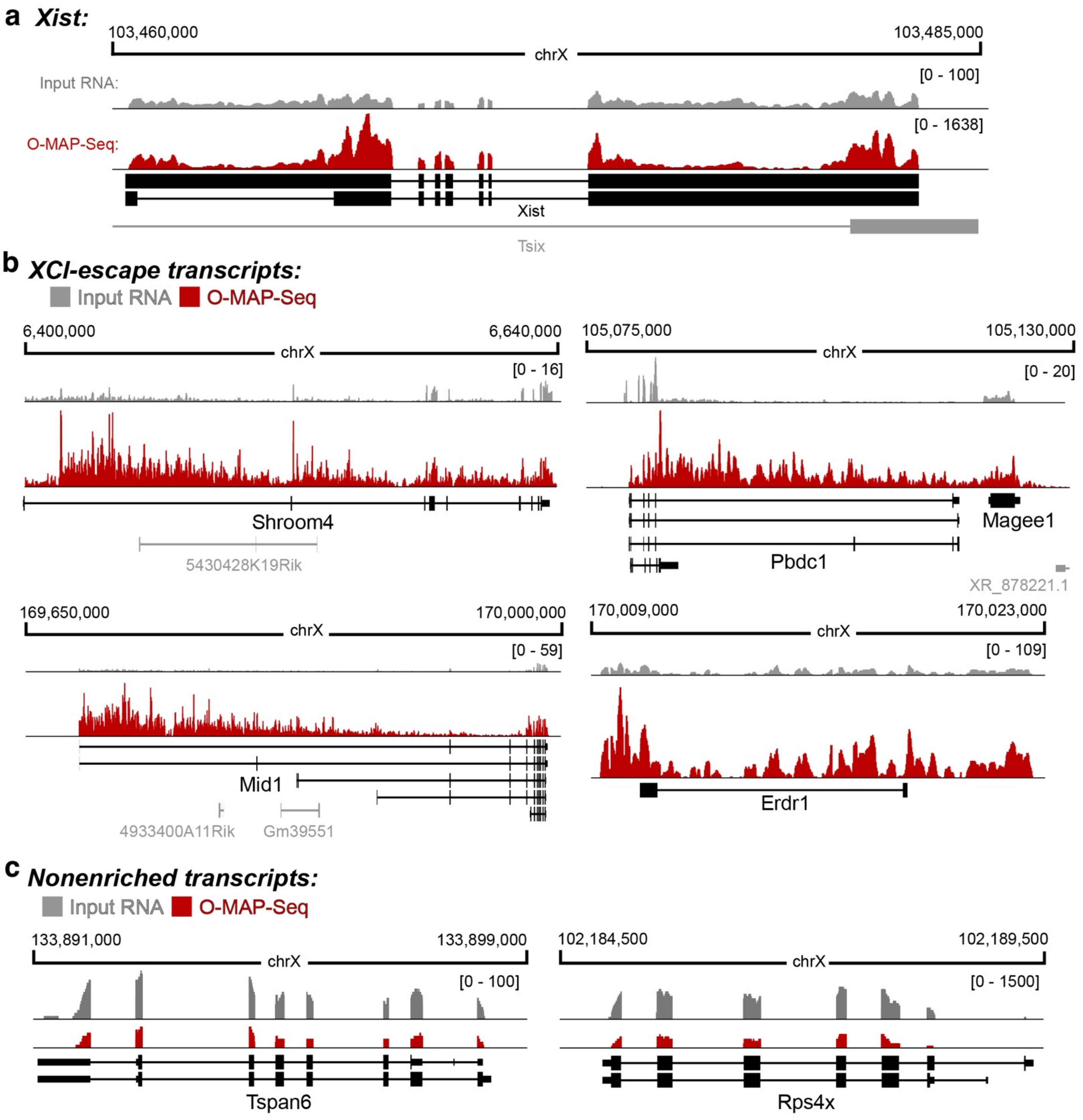


**Supplementary Figure 15. *Xist* O-MAP-Seq enriches nascent transcripts of XCI-escape genes**. **a,** Enrichment of the *Xist* gene itself. Note different scales for input RNA and O-MAP-Seq tracks. The lack of intronic reads suggests that O-MAP-Seq has predominantly targeted and captured the mature *Xist* transcript. The absence of reads mapping to the antisense noncoding RNA *TsiX,* a lncRNA that is monoallelically expressed from the Xa (*gray*), confirms that O-MAP-Seq is precisely labeling the Xi. **b,** Enriched XCI-escape genes appear to be nascent transcripts. In all cases, read densities for both Input RNA (gray) and O-MAP-Seq (red) are shown, using matched scales. Note prominent intronic read density for all XCI-escape genes (*Shroom4*, *Pbdc1*, *Magee1, Mid1, Erdr1*). Transcript structures denoting the most prominent isoforms are displayed; other nearby genes not known to escape XCI are denoted in gray. **c,** *Xist* O-MAP-Seq does not appear to preferentially capture nascent transcripts of non-XCI-escape genes. Two examples (*Tspan6* and *Rps4x*) are shown, neither of which was enriched by *Xist* O-MAP-Seq. Note the absence of prominent intronic read density, suggesting that the intronic signatures observed in (**b**) are not general artifacts of the O-MAP-Seq pipeline.

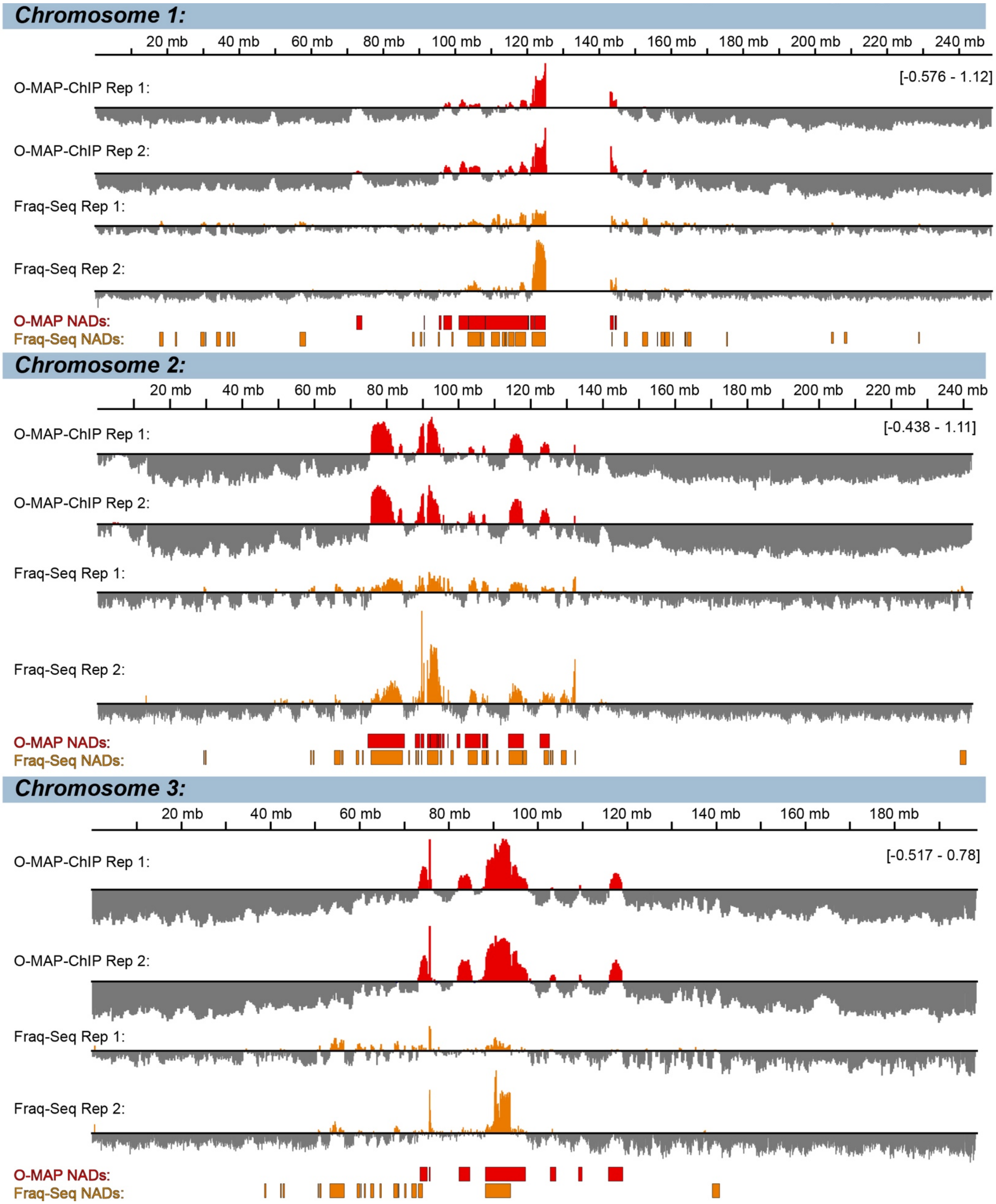


**Supplementary Figure 16. Genomic maps of HeLa Nucleolar-Associated Domains (NADs)**. Data are $Log_2$(O-MAP-ChIP/Input). NADs were called by merging peaks from EDD and EPIC2 (*red blocks*). The same pipeline was also applied to HT1080 NADs, probed by fractionation-sequencing (Fraq-Seq; PMID 20826608). Although all replicates are shown, high noise/low information in Fraq-Seq replicate 2 resulted in EDD failure after 10,000 Monte Carlo simulations; only replicate 1 was used for NAD calls (*orange blocks*). Figure continued on the next seven pages.

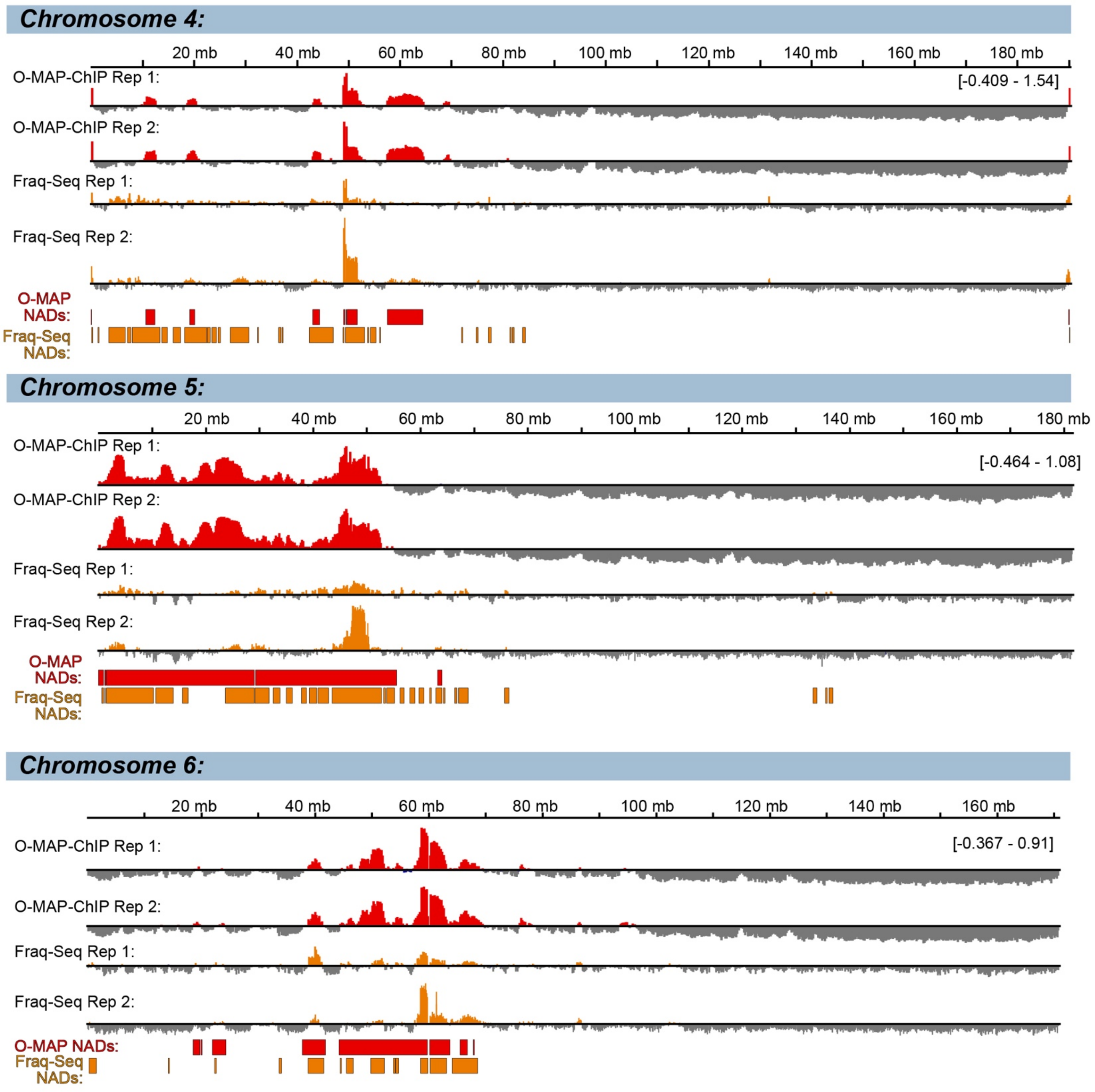


**Supplementary Figure 16,** *continued***.**

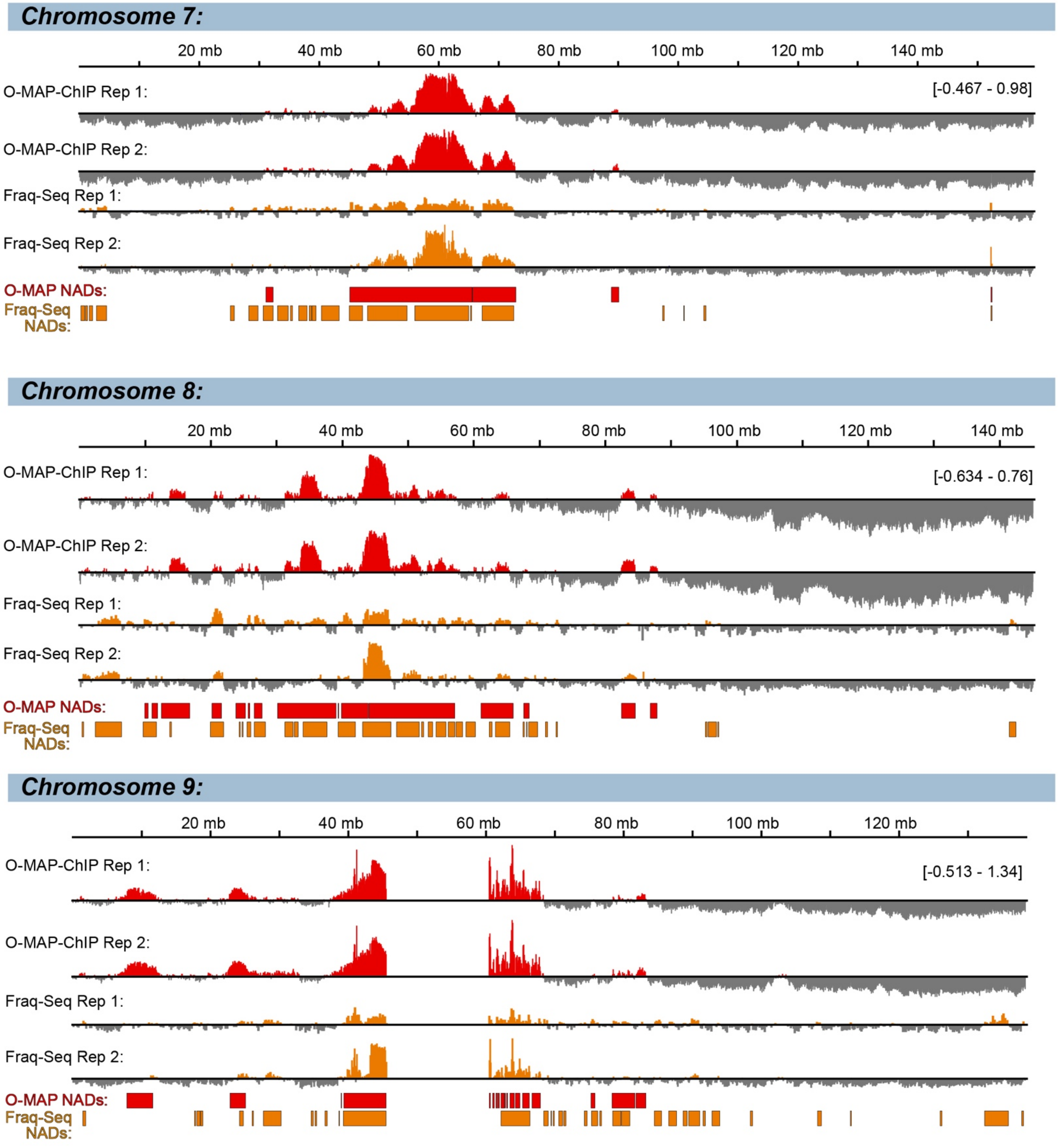


**Supplementary Figure 16**, *continued*.

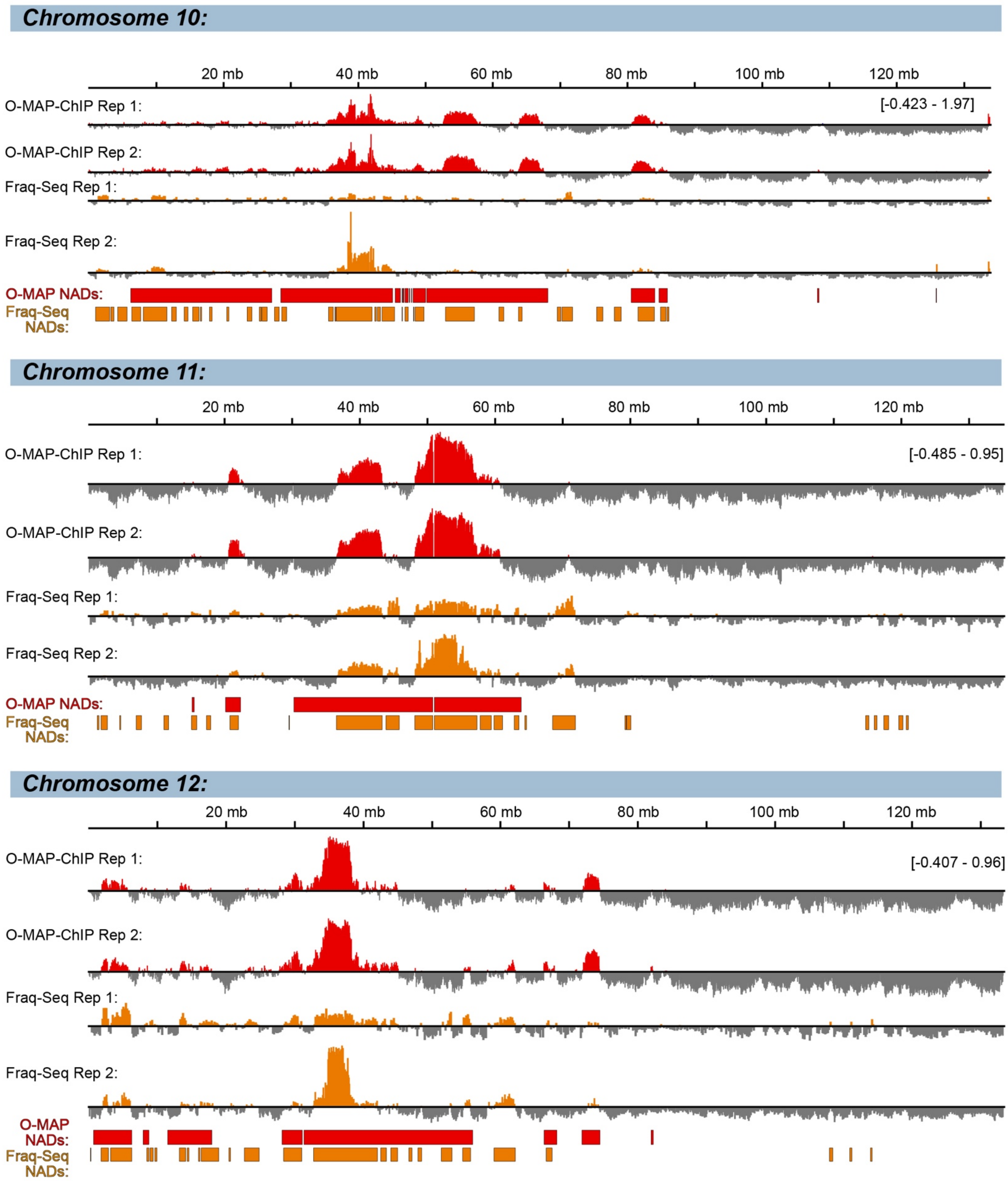


**Supplementary Figure 16,** *continued.*

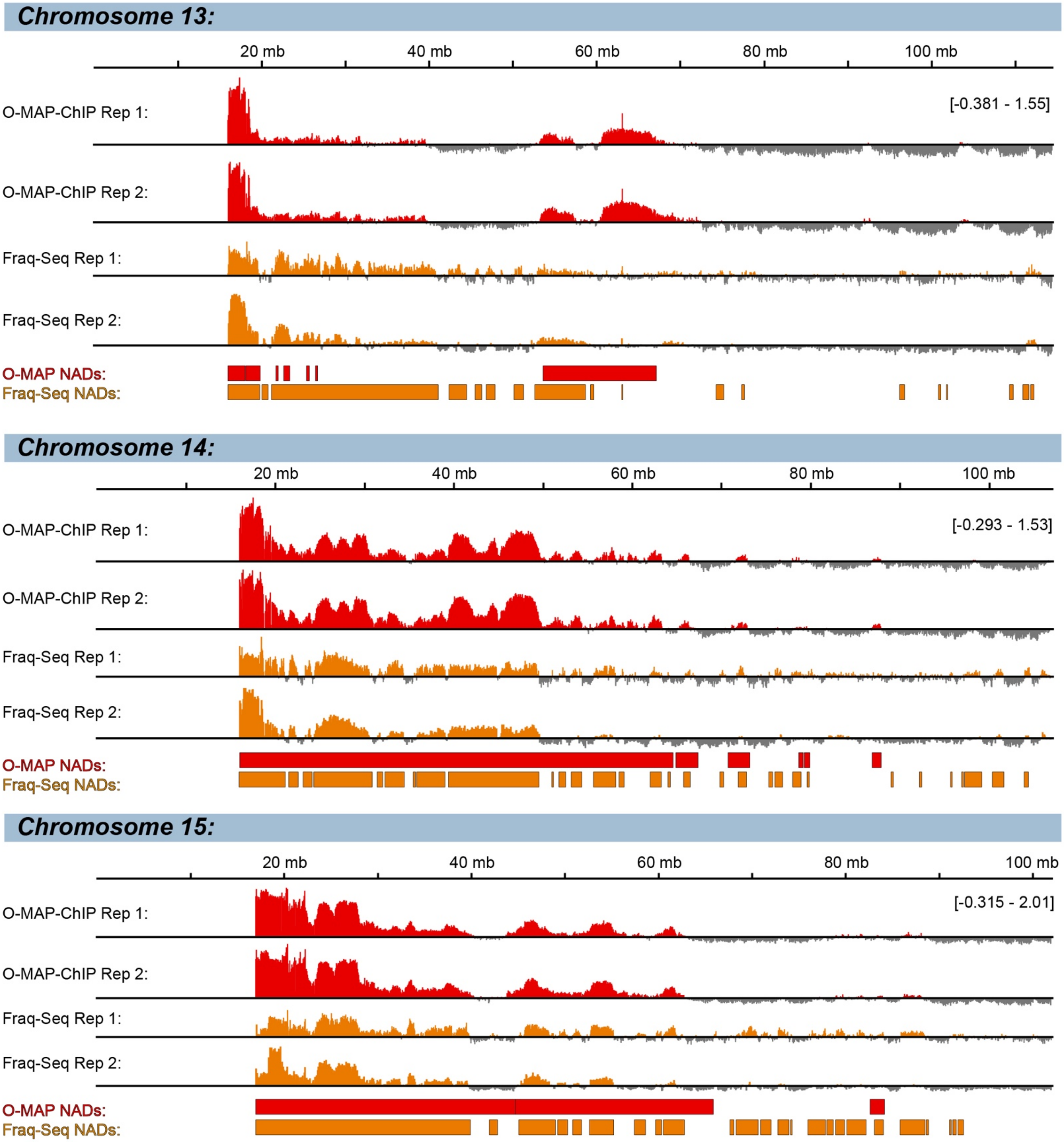


**Supplementary Figure 16,** *continued.*

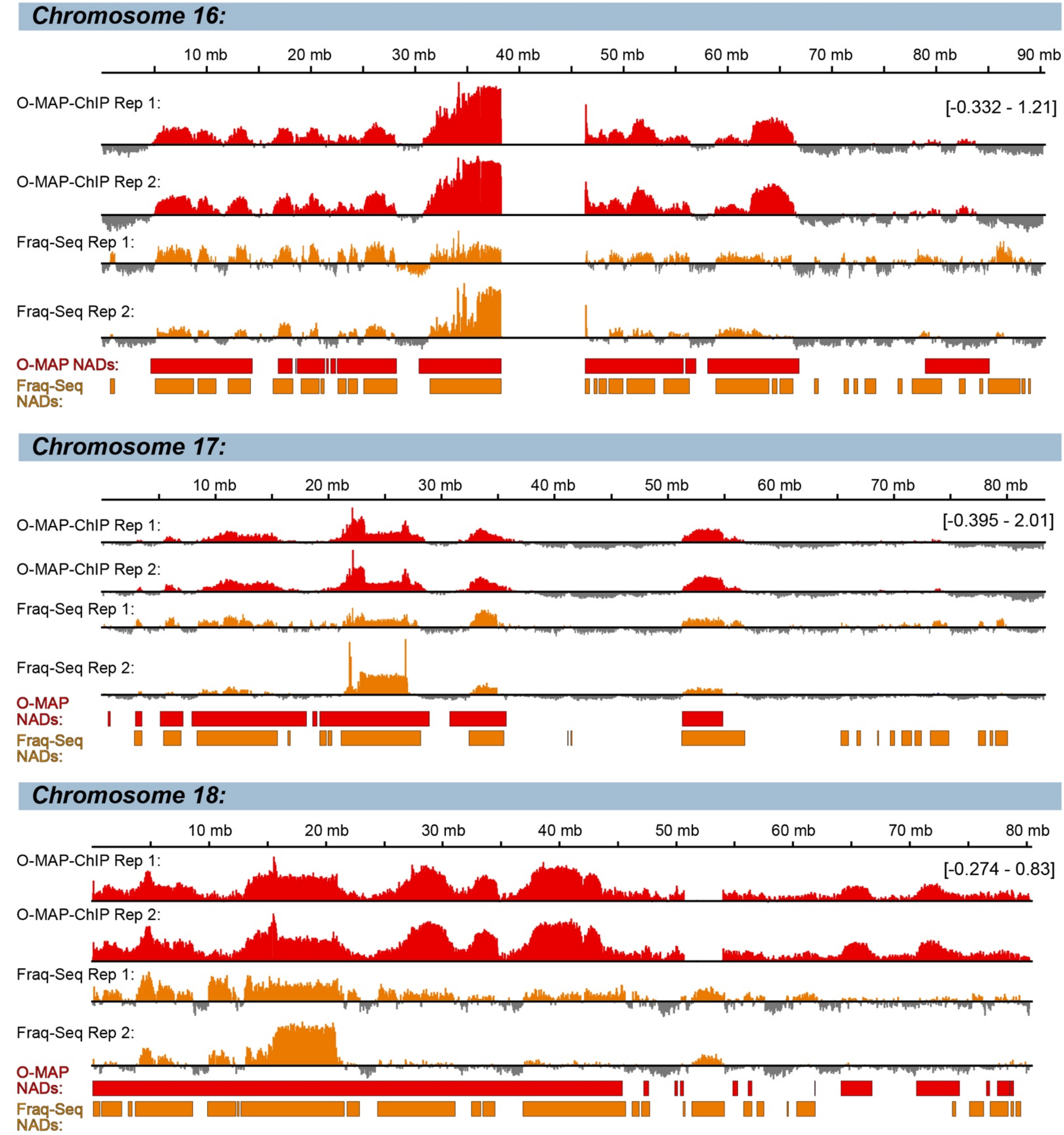


**Supplementary Figure 16,** *continued.*

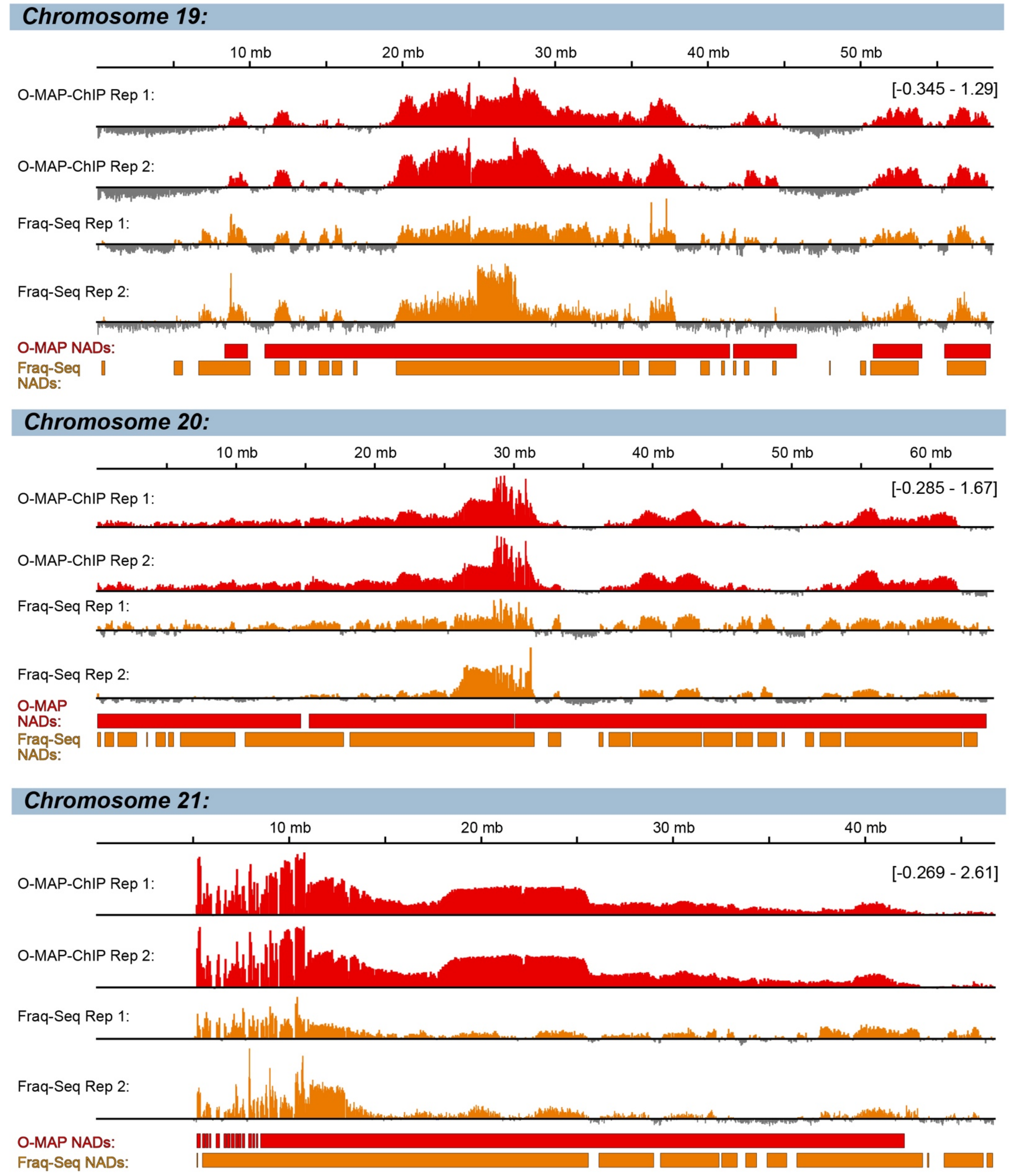


**Supplementary Figure 16,** *continued.*

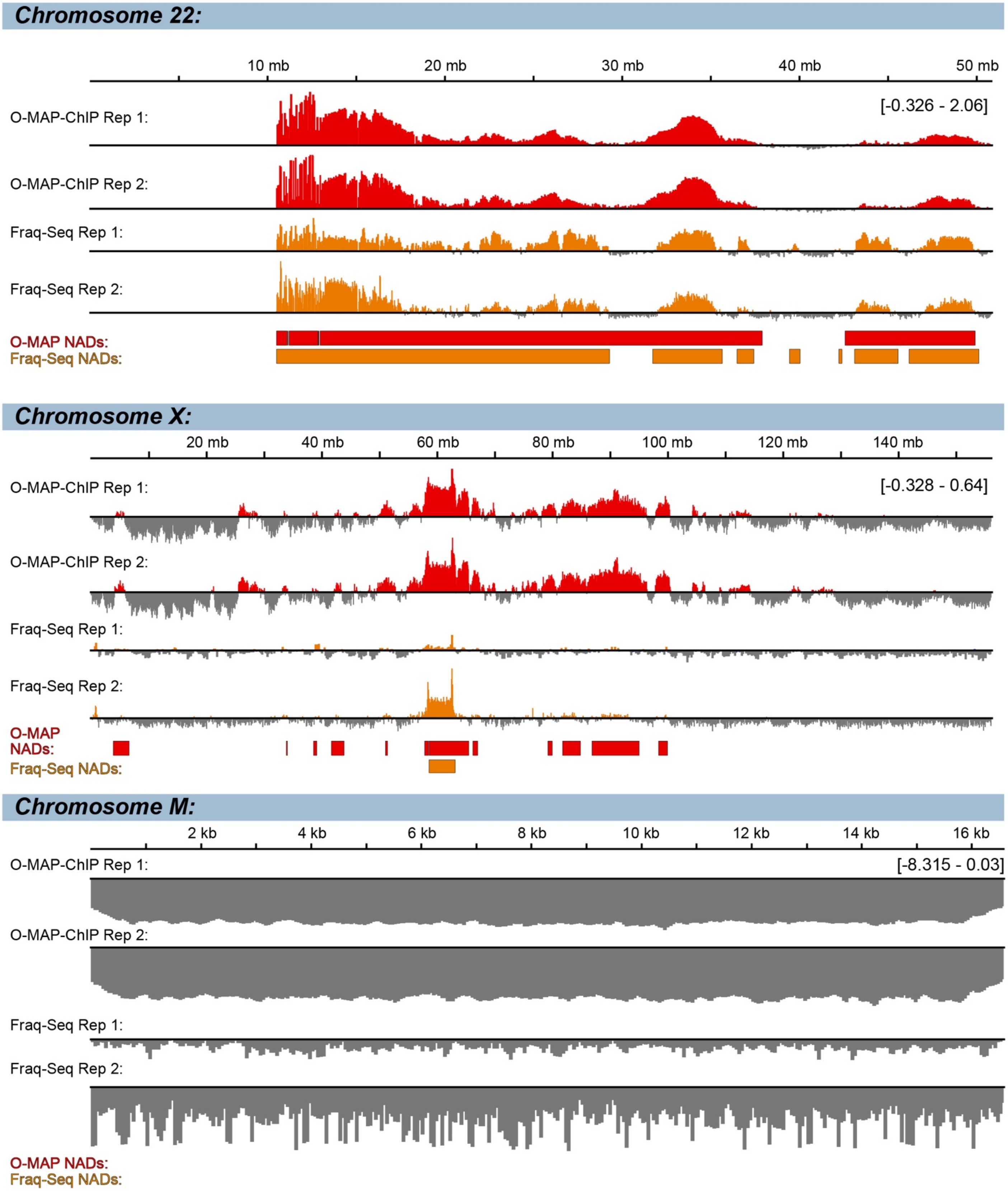


**Supplementary Figure 16,** *continued*. The mitochondrial genome exhibits a complete absence of nucleolar interactions, as expected. Note that HeLa (O-MAP-ChIP) and HT1080 (Fraq-Seq) cells were derived from patients of different genders, which may explain the divergence in NAD architecture on the X-chromosome.

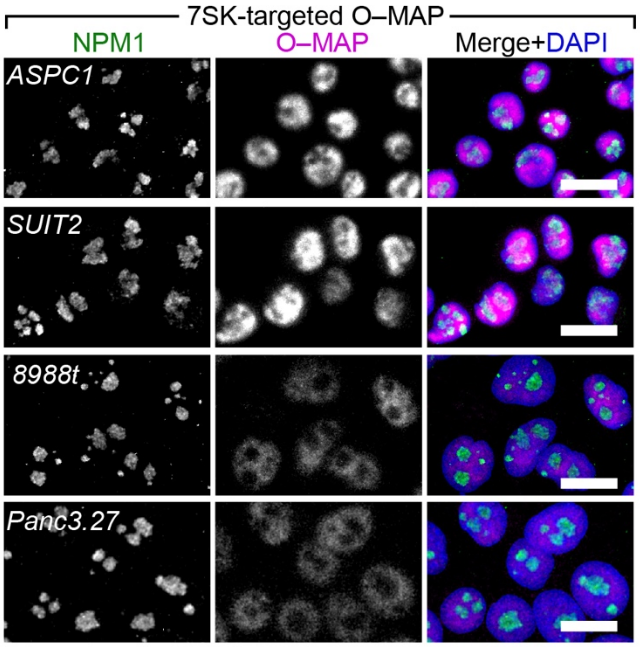


**Supplementary Figure 17. 7SK O-MAP in cultured Pancreatic Ductal Adenocarcinoma (PDA) cell lines.** O-MAP was visualized using a fluorescent neutravidin conjugate; *NPM1* by immunofluorescence, as in (**Fig. 5**). Scale bars: 20 µm.
